# Perturbative formulation of general continuous-time Markov model of sequence evolution via insertions/deletions, Part II: Perturbation analyses

**DOI:** 10.1101/023606

**Authors:** Kiyoshi Ezawa, Dan Graur, Giddy Landan

## Abstract

**Background:** Insertions and deletions (indels) account for more nucleotide differences between two related DNA sequences than substitutions do, and thus it is imperative to develop a stochastic evolutionary model that enables us to reliably calculate the probability of the sequence evolution through indel processes. In a separate paper (Ezawa, Graur and Landan 2015a), we established a theoretical basis of our *ab initio* perturbative formulation of a *genuine* evolutionary model, more specifically, a continuous-time Markov model of the evolution of an *entire* sequence via insertions and deletions. And we showed that, under some conditions, the *ab initio* probability of an alignment can be factorized into the product of an overall factor and contributions from regions (or local alignments) separated by gapless columns.

**Results:** This paper describes how our *ab initio* perturbative formulation can be concretely used to approximately calculate the probabilities of all types of local pairwise alignments (PWAs) and some typical types of local multiple sequence alignments (MSAs). For each local alignment type, we calculated the fewest-indel contribution and the next-fewest-indel contribution to its probability, and we compared them under various conditions. We also derived a system of integral equations that can be numerically solved to give “exact solutions” for some common types of local PWAs. And we compared the obtained “exact solutions” with the fewest-indel contributions. The results indicated that even the fewest-indel terms alone can quite accurately approximate the probabilities of local alignments, as long as the segments and the branches in the tree are of modest lengths. Moreover, in the light of our formulation, we examined parameter regions where other indel models can safely approximate the correct evolutionary probabilities. The analyses also suggested some modifications necessary for these models to improve the accuracy of their probability estimations.

**Conclusions:** At least under modest conditions, our *ab initio* perturbative formulation can quite accurately calculate alignment probabilities under biologically realistic indel models. It also provides a sound reference point that other indel models can be compared to. [This paper and three other papers (Ezawa, Graur and Landan 2015a,b,c) describe a series of our efforts to develop, apply, and extend the *ab initio* perturbative formulation of a general continuous-time Markov model of indels.]

## Introduction

The evolution of DNA, RNA, and protein sequences is driven by mutations such as base substitutions, insertions and deletions (indels), recombination, and other genomic rearrangements (*e.g.*, Graur and Li 2000; Gascuel 2005; Lynch 2007). Thus far, analyses on substitutions have predominated in the field of molecular evolutionary study, in particular using the probabilistic (or likelihood) theory of substitutions that is now widely accepted (e.g., Felsenstein 1981, 2004; Yang 2006). However, some recent comparative genomic analyses have revealed that indels account for more base differences between the genomes of closely related species than substitutions (*e.g*., Britten 2002; Britten et al. 2003; Kent *et al*. 2003; The International Chimpanzee Chromosome 22 Consortium 2004; The Chimpanzee Sequencing and Analysis Consortium 2005). It is therefore imperative to develop a stochastic model that enables us to reliably calculate the probability of sequence evolution via mutations including insertions and deletions.

Since the groundbreaking works by Bishop and Thompson (1986) and by Thorne, Kishino and Felsenstein (1991), many studies have been done to develop and apply methods to calculate the probabilities of pairwise alignments (PWAs) and multiple sequence alignments (MSAs) under the probabilistic models aiming to incorporate the effects of indels, and such methods have greatly improved in terms of the computational efficiency and the scope of application. See excellent reviews for details on this topic (e.g., Rivas 2005; Bradley and Holmes 2007; Miklós et al. 2009). A majority of these studies are based on hidden Markov models (HMMs) or transducer theories. Both of them calculate the indel component of an alignment probability as a product of inter-column transition probabilities or of block-wise contributions. Unfortunately, they have two fundamental problems, one regarding the theoretical grounds and the other regarding the biological realism. Regarding the theoretical grounds, it is unclear whether or not a HMM or a transducer is related with any *genuine* evolutionary model, which describes the evolution of an *entire* sequence via indels along the time axis. Regarding the biological realism, the standard HMMs or transducers can at best handle geometric distributions of indel lengths, whereas many empirical studies showed that the real indel lengths are distributed according to the power-law (*e.g*., Gonnet et al. 1992; Benner et al. 1993; Gu and Li 1995; Kent et al. 2003; Zhang and Gerstein 2003; Chang and Benner 2004; The international Chimpanzee Chromosome 22 Consortium 2004; Yamane et al. 2006; Fan et al. 2007). Besides, it is very hard for the previous methods to incorporate the indel rate variation across regions, which were often observed empirically (*e.g*., Gu et al. 2008). See the “background” section in part I (Ezawa, Graur and Landan 2015a) for more details on these problems.

In part I of this series of study (Ezawa, Graur and Landan 2015a), we established an *ab initio* formulation of a *genuine* stochastic evolutionary model, that is, a general continuous-time Markov model of sequence evolution via indels. Our evolutionary model allows any indel rate parameters including length distributions, but it does not impose any unnatural restrictions on indels. Thus, the model is naturally devoid of the aforementioned two problems. Aided by some techniques of the perturbation theory in physics (Dirac 1958; Messiah 1961a,b), we formally expanded the probability of an alignment into a series of terms with different numbers of indels. This expansion theoretically underpinned the stochastic evolutionary simulation method of Gillespie (1977). And we also showed that, if the indel model parameters satisfy a certain set of conditions, the *ab initio* probability of an alignment is indeed factorable into the product of an overall factor and contributions from local alignments delimited by preserved ancestral sites (PASs). This result reconfirmed and generalized the conjecture that Miklós et al. (2004) made under their spatially and temporally homogeneous indel model.

In this paper, we turn our attention to more concrete problems. In other words, we focus on how to concretely calculate the contribution from each local alignment, assuming that the indel model satisfies the conditions for the factorability of alignment probabilities.

In Section 1 of Results & Discussion, we demonstrate how our *ab initio* perturbative formulation can be concretely used to approximately calculate the contributions to the alignment probabilities from regions separated by gapless columns (*i.e*., local alignments). We examine all types of local pairwise alignments (PWAs) and some typical types of local multiple sequence alignments (MSAs). For each local alignment type, we calculate the fewest-indel contribution and the next-fewest-indel contribution to its probability. In this section, we also derive a system of integral equations that can be numerically solved to give “exact solutions” for some common types of local PWAs. Then, by comparing the fewest-indel contribution with the next-fewest-indel contribution, or with the “exact” solution, we examine the parameter region in which the fewest-indel contribution can approximate the alignment probability quite accurately. In Section 2 of Results & Discussion, we examine some representative models that were used in the past studies, including the HMM of Kim and Sinha (2007), in the light of our *ab initio* perturbative formulation. We attempt to delimit parameter regions where these models can safely approximate the *ab initio* probabilities under the *genuine* evolutionary model. The analyses will also suggest some modifications to these models that would enhance their reliability. And Appendix describes details on some perturbation calculations.

This paper is part II of a series of our papers that documents our efforts to develop, apply, and extend the *ab initio* perturbative formulation of the general continuous-time Markov model of sequence evolution via indels. Part I (Ezawa, Graur and Landan 2015a) gives the theoretical basis of this entire study. Part II (this paper) describes concrete perturbation calculations and examines the applicable ranges of other probabilistic models of indels. Part III (Ezawa, Graur and Landan 2015b) describes our algorithm to calculate the first approximation of the probability of a given MSA and simulation analyses to validate the algorithm. Finally, part IV (Ezawa, Graur and Landan 2015c) discusses how our formulation can incorporate substitutions and other mutations, such as duplications and inversions.

This paper basically uses the same conventions as used in part I (Ezawa, Graur and Landan 2015a). Briefly, a sequence state *s* (∈ *S*) is represented as an array of sites, each of which is either blank or equipped with some specific attributes. And each indel event is represented as an operator acting on the bra-vector, 〈*s* |, representing a sequence state. More specifically, the operator 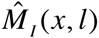 denotes the insertion of *l* sites between the *x* th and (*x* +1) th sites, and the operator 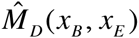 denotes the deletion of a sub-array between (and including) the *x*_*B*_ th and the *x*_*E*_ th sites. See Section 2 of part I for more details.

And, also as in part I, the following terminology is used. The term “an indel process” means a series of successive indel events with both the order and the specific timings specified, and the term “an indel history” means a series of successive indel events with only the order specified. And, throughout this paper, the union symbol, such as in *A ⋃ B* and 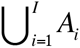, should be regarded as the union of *mutually disjoint* sets (*i.e.,* those satisfying *A* ∩ *B* = ∅ and *A*_*i*_ ∩ *A*_*j*_ = ∅ for *i* ≠ *j* (∈ {1,…, *I*}), respectively, where is an empty set), unless otherwise stated.

## Results & Discussion

### 1. Perturbation approximations of probabilities

1.1 Perturbation expansion of multiplication factor contributed from local indel histories

In Section 4 of part I (Ezawa, Graur and Landan 2015a), we showed that the alignment probability can be factorized into the product of an overall factor and contributions from regions separated by gapless columns (as in Eqs.(4.1.8a,b) of part I for a PWA and Eqs.(4.2.9a,b,c) of part I for a MSA), if the following conditions are satisfied.

Condition (i): The indel rate parameters are independent of the portion of the sequence state outside of the region in question.

Condition (ii): The increment of the exit rate due to an indel event is independent of the portion of the sequence state outside of the region in question.

Condition (iii) (necessary only for MSAs): The probability of the root sequence state is factorable into the product of an overall factor and regional contributions. (This condition is represented as Eq.(4.2.8) of part I.)

Each regional contribution of the probability was symbolically represented as 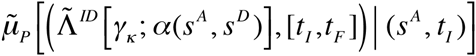 (in Eq.(4.1.8b) of part I) for a PWA and 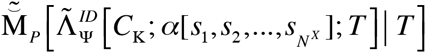 (in Eq.(4.2.9c) of part I) for a MSA. (See below for the definitions of the symbols for the arguments.) Thus, as long as we can concretely calculate these regional contributions, we can obtain the specific value of the probability of a given alignment. However, because each of these factors is in general a summation of contributions from an infinite number of local indel histories, we usually need to approximate it by a summation over a finite number of histories. In this section, we will do this, again based on the perturbation expansion as unfolded in Section 3 of part I.

For a PWA, α (*s*^*A*^, *s*^*D*^), we decompose 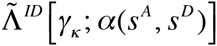, the set of local indel histories consistent with the local PWA confined in the region γ_*k*_, as: 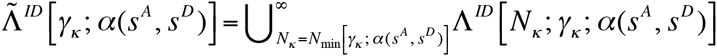. Here *Λ*^ID^ [*N*_*k*_; γ_*k*_; α(*s*^*A*^, *s*^*D*^)] is the subset of PWA-consistent histories in γ_*κ*_, each of which consists of *N*_*κ*_ indels, and *N*_min_ [γ_*k*_; α(*s*^*A*^, *s*^*D*^)] is the minimum number of indels required to give rise to the local PWA of α(*s*^*A*^, *s*^*D*^) within γ_*κ*_. Then, the “perturbation expansion” of the multiplication factor, 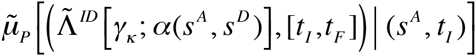, for the local PWA probability produced during time interval [*t*_*I*_, *t*_*F*_], is given by:

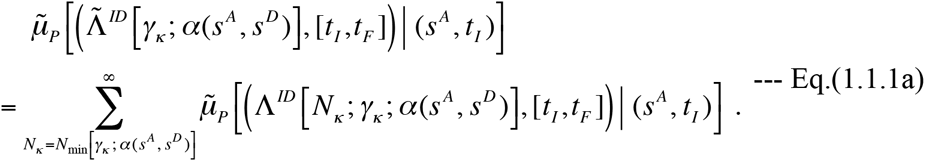

Here

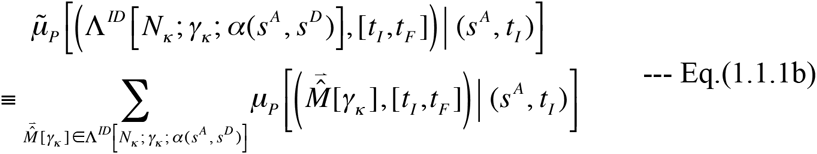

is the multiplication factor contributed from all PWA-consistent *N*_k_-event local indel histories.

For a MSA, α[*s*_1_, *s*_2_,…, *s*_*N*_*X*], we decompose 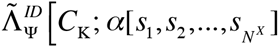, the set of local indel histories along the tree (*T*) consistent with the local MSA confined in the region *C*_k_, as: 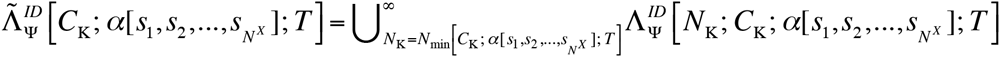.

Here, 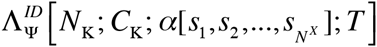 is the subset consisting of all MSA-consistent *N*_k_-event local indel histories along the tree, and *N*_min_ *C*_k_; α[*s*_1_, *s*_2_,…, *s*_*N*_*x*]; *T]* is the minimum possible number of indels that can produce the sub-MSA within *C*_k_. Using this, the “perturbation expansion” of the multiplication factor, 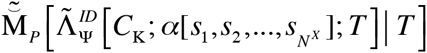, is given by:

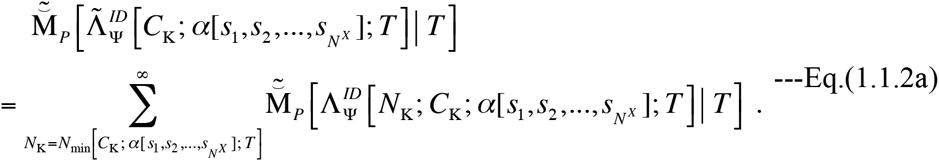

Here

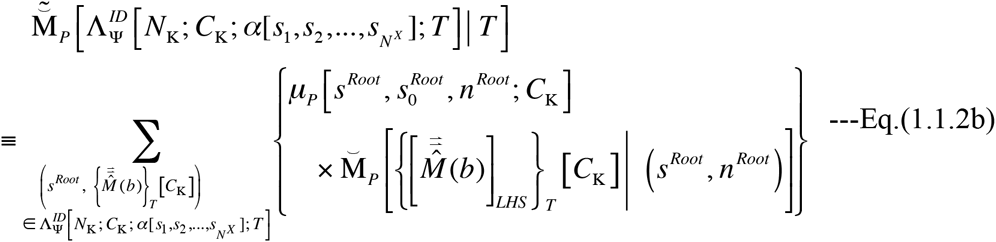

is the multiplication factor contributed from all local histories in 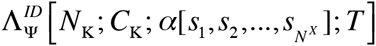.

Thus, the problems were reduced to those of how to enumerate the local indel histories in the subset, Λ^*ID*^ [*N*_*k*_ γ_*k*_; α(*s*^*A*^, *s*^*D*^)] (for a PWA) or 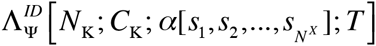 (for a MSA), up to a particular number of events, *N*_k_or *N*_k_, that was chosen to achieve a desired level of accuracy. Once the relevant local histories are enumerated, then, we can calculate the contribution from each of them according to the general formula. (The general formulas are in Eq.(4.1.1b) of part I accompanied by Eq.(3.1.8b) of part I for a local PWA, and in Eq.(4.2.6b) of part I accompanied by Eq.(4.2.4b) of part I for a local MSA.) In Subsections 1.2 and 1.3, we explicitly calculate the portions of the multiplication factors, 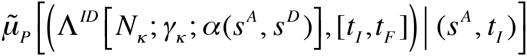 and 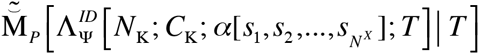, for some typical gap-configurations of the local PWAs and MSAs, respectively, that are contributed from the parsimonious local indel histories. Then we compare them with the portions contributed from the next-parsimonious local indel histories. Moreover, for a few simplest but commonest types of local PWAs, we also derive integral equation systems that give the “exact solutions” for the total multiplication factors, Eq.(1.1.1a). Then, we see how well the contributions from parsimonious indel histories alone can approximate the actual occurrence frequencies of the gap-configurations of local MSAs.

In this section, we assume that we are working with a model whose alignment probabilities are factorable, and that we are focusing on calculating the multiplication factor that comes from a single local alignment flanked by a pair of gapless columns (*i.e*., a pair of PASs). We will mainly work in the state space *S* = *S*^*II*^ (see Section 2 of part I for the definition of the space). This means that we will focus on calculating the probability of the *homology structure* of each local alignment (see, *e.g*., Lunter et al. 2005). Let ∈*L*(*s*) be the number of sites that a sequence *s* ∈ *S*^*II*^ has between the pair of PASs. We will re-assign the site numbers so that the left-and right-flanking PASs are numbered 1 and ∈*L*(*s*) ∈ 2, respectively, and the sites in between them are numbered 2,…, ∈*L*(*s*) ∈1. This will make it easy to apply the theory formulated thus far to the current situation. We will re-assign the ancestries *v*(1) *L* and *v*(∈*L*(*s*) ∈ 2) = *R* to the left-and right-flanking PASs, respectively. And we will usually re-assign the ancestries *v*(*x*) *x* − 1 to the sites, *x* = 2,…, ∈*L*(*s*) ∈1, of the ancestral sequence *s* = *s*^*A*^ (or the root sequence *s* = *s*^*Root*^ for a MSA) in between the PASs (see Figure 1 as an illustration).

**Figure 1.**
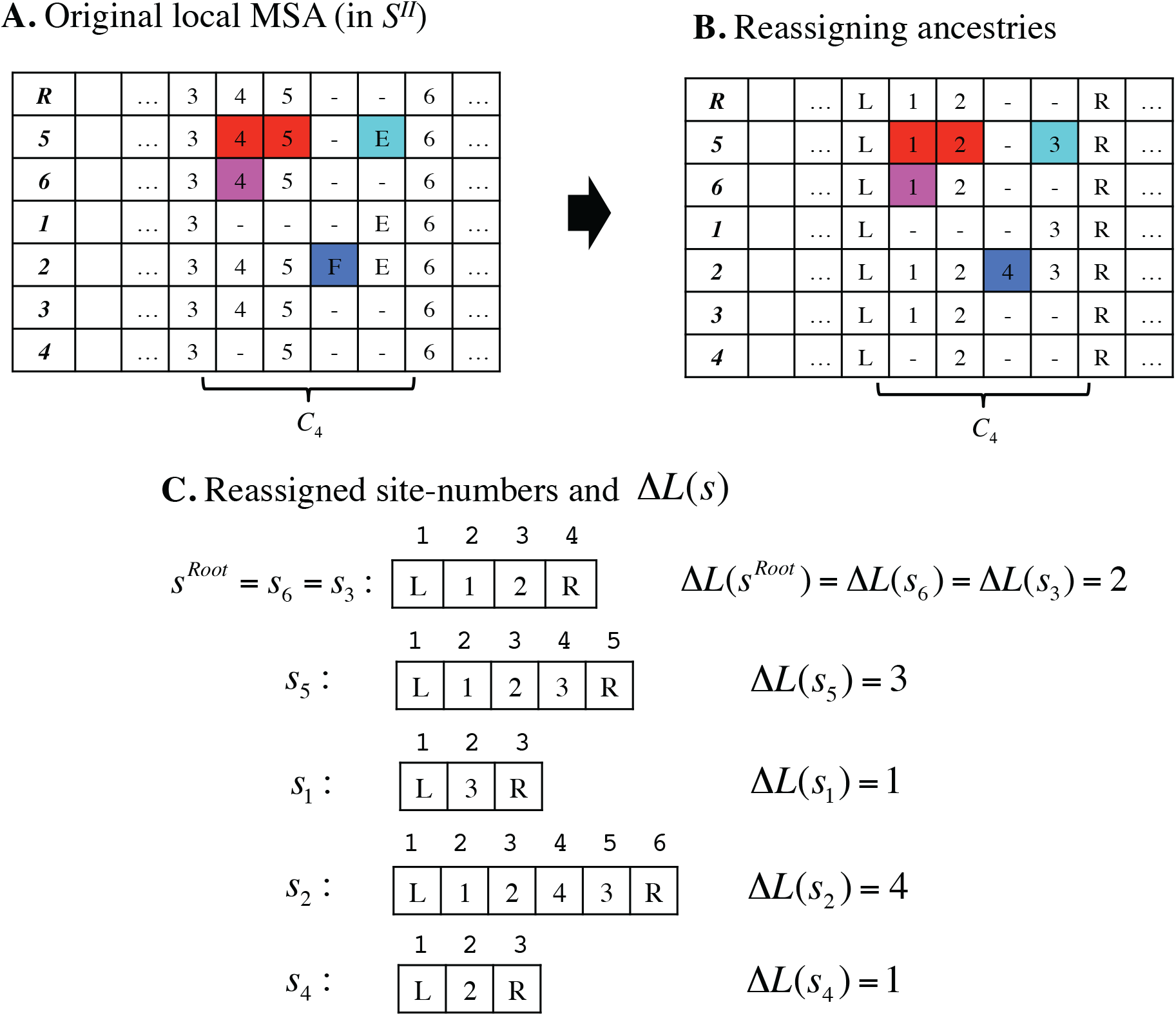
Typical situation and notation considered in Section 1 of Results. **A.** As an illustration, we use the local MSA confined in the region *C*_4_ of the MSA in the Figure 11 B of part I (Ezawa, Graur and Landan 2015a), and re-assign the ancestries as shown in panel **B**. The ancestries in between the PASs (with the re-assigned ancestries *L* and *R*) are just an example and not always assigned in this way. **C.** Sequences extracted from the MSA. Shown above each site is the site number (*i.e*., spatial coordinate) assigned to it. And shown on the right of each sequence is the count of sites in between the PASs. In panels A and B, as in Figure 5 of part I, the boldface characters in the leftmost columns stand for the sequence IDs. (More precisely, the number ‘*i*’ stands for the sequence *s*_*i*_, and the ‘*R*’ stands for the root sequence (*s*^*Root*^).)

In the following subsections, we will omit the symbol “ []_*LHS*_ “ that denoted a local history set (LHS) in part I. Because we will consider a PWA or MSA region that accommodates only one local history, the resulting LHS equivalence classes are trivial classes, each of which consists only of a single history. We will also omit the appended “ [*C*_k_] “ to indicate the MSA region when it is obvious.

### 1.2. Perturbation analyses on local PWA

In a PWA, α(*s*^*A*^, *s*^*D*^), a gap-configuration flanked by a pair of PASs corresponds to the portion of α(*s*^*A*^, *s*^*D*^) confined in γ_*k*_. If we are interested only in the homology relationships among sites (*i.e*., homology structures (Lunter et al. 2005)), there are four broad types of gap-configurations (see Figure 2; see also Subsection 3.3 of part I (Ezawa, Graur and Landan 2015a) for complexities concerning this issue). (i) Neither the ancestral nor the descendant sequence has even a single site in between the pair of PASs (panel A of Figure 2). (ii) The ancestor has one or more site(s) but the descendant has no site in between the PASs (panel B). (iii) The ancestor has no site but the descendant has one or more sites in between (panel C). And (iv) both the ancestor and the descendant have one or more site(s) in between, but no ancestral site is related to any of the descendant sites (panel D). We will consider these cases in turn.

**Figure 2.**
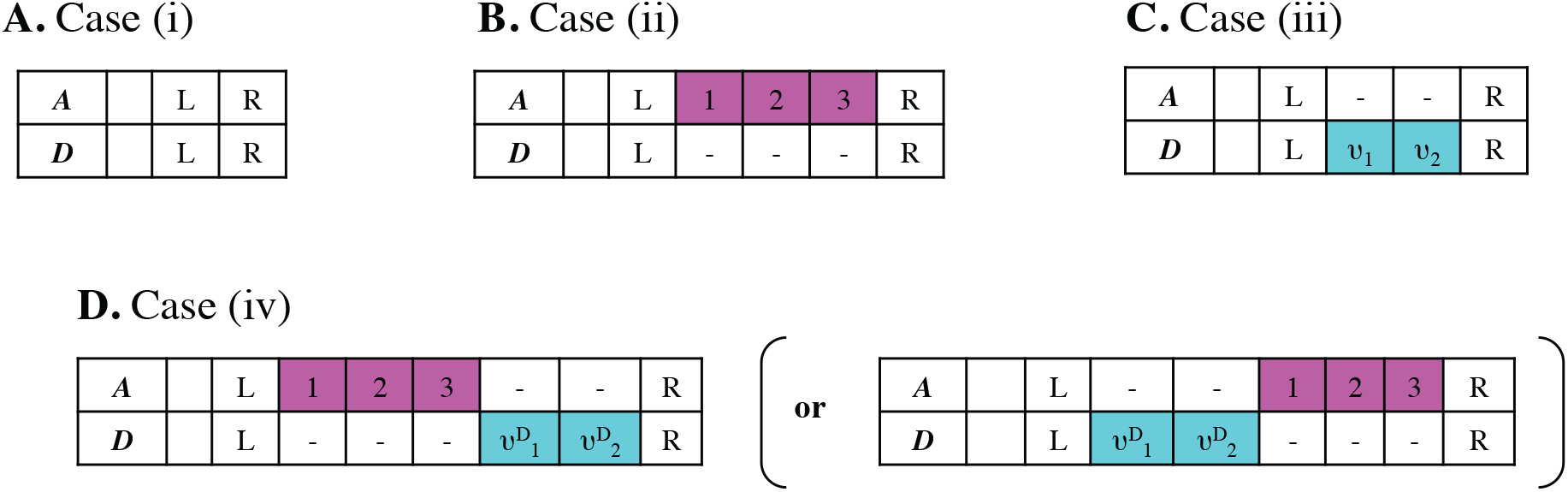
Examples of four types of local gap configurations in PWA between ancestral and descendant sequences. **A.** Case (i). **B.** Case (ii) with Δ*L*^*A*^ = 3. **C.** Case (iii) with Δ*L*^*D*^ = 2. **D**. Case (iv) with Δ*L*^*A*^ = 3 and Δ*L*^*D*^ = 2. In each PWA, each site (a cell) is assigned an ancestry. In the leftmost column of each PWA, the boldface italic ‘A’ and ‘D’ stand for an ancestor (*s*^*A*^) and a descendant (*s*^*D*^), respectively. The magenta boxes and the cyan boxes represent unpreserved ancestral sites and inserted descendant sites, respectively. In panel D, the PWA on the right (in parentheses) is equivalent to the PWA on the left, as far as the homology structure alone is concerned.

In case (i), the sequence states could be represented as *s*^*A*^ = *s*^*D*^ = [*L*, *R*]. In this case, *N*_min_ [γ_*k*_; α(*s*^*A*^, *s*^*D*^)] = 0, and thus there is only one fewest-indel history, [], where no indel event takes place. Therefore, in this case, the portion of the multiplication factor contributed by the fewest-indel history is:

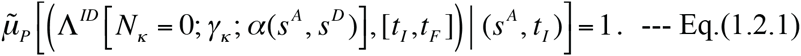

In this case, there is no history consisting only of one indel event. And each next-fewest-indel history should be a two-event history of the form, 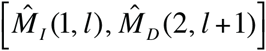 with *l* = 1,…, *L*^*CO*^, where 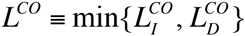. Thus, the portion contributed by the next-fewest-indel histories is:

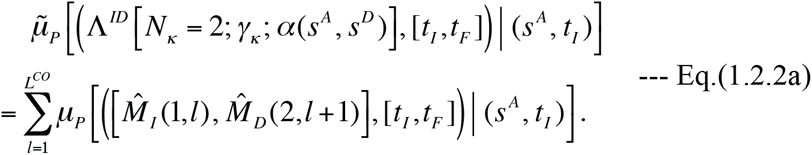

Let 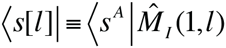 be the state resulting from the action of an insertion of *l* sites on *s*^*A*^. Then, using Eq.(4.1.1b) and Eq.(3.1.8b) of part I, each summand is calculated as:

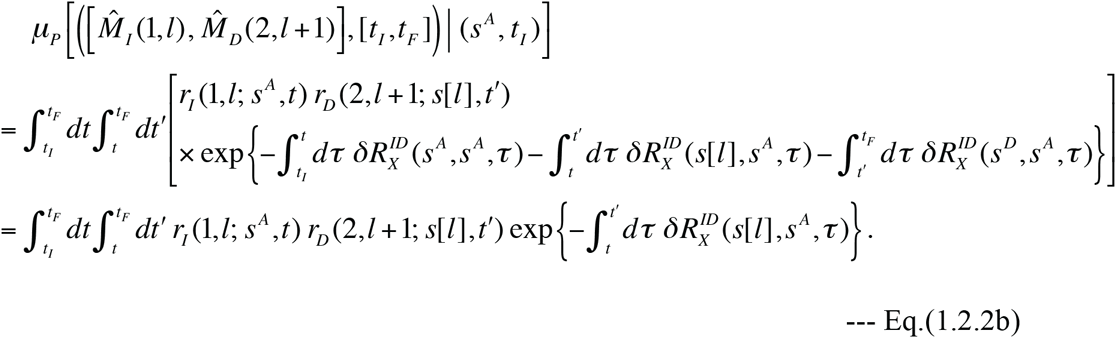

Here, as in Subsection 4.1 of part I, 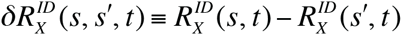 is the increment of the exit rate, which in turn is given by Eq.(2.4.1b’) of part I. The second equation of Eq.(1.2.2b) follows from 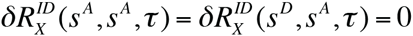. We could at least numerically calculate Eq.(1.2.2b) once the specific functional forms of the indel rates and the exit rates are given. For example, in a space-time-homogeneous model, like Dawg’s indel model (Cartwright 2005), Eqs.(2.4.4a,b) of part I, we have 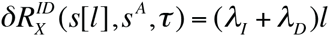, and Eq.(1.2.2b) is calculated as:

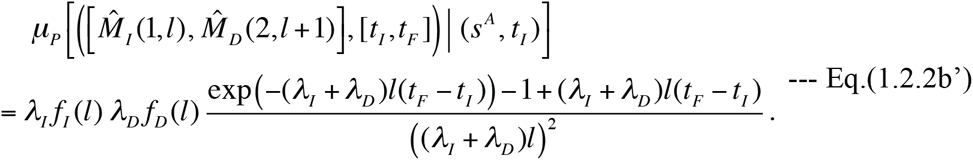

Exactly the same expression can be obtained also from the “long indel” model, Eqs.(2.4.5a,b) of part I, if we notice the correspondence, Eqs.(2.4.7a,b,c,d) of part I.) Eq.(1.2.2b’) (or Eq.(1.2.2b) itself) indicates that 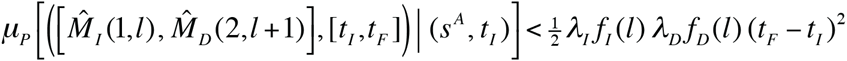 for each *l* = 1,…, *L*^*CO*^. Applying this inequality to Eq.(1.2.2a) and using another inequality, 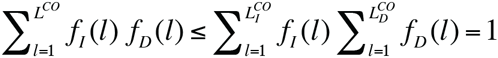, we have:

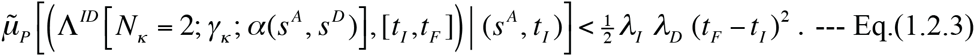

Empirically, the rate of indels (λ_*I*_ + λ_*D*_) is estimated to be at most on the order of 1/10 of the substitution rate (Lunter 2007; Cartwright 2009). Eq.(1.2.3) indicates that, in case (i), even if the elapsed time (*t*_*F*_ − *t*_*I*_) is such that the substitution process is nearly saturated, e.g., λ_*S*_ (*t*_*F*_ − *t*_*I*_) ≈ 4, where λ_*S*_ is the total substitution rate per site, the total contribution from the next-fewest-indel histories, Eq.(1.2.2a), is at most on the order of 1/10 of the contribution by the fewest-indel history, Eq.(1.2.1). Thus, we expect that the fewest-indel contribution should approximate the entire multiplication factor (Eq.(4.1.8b) in part I) very well in case (i).

Incidentally, taking advantage of the result for case (i), we can calculate the multiplication factor for a gapless PWA segment consisting of *L*^*P*^ (> 2) PASs. For clarity, let us consider an indel model as discussed in Subsection V-1. Then, in each of the *L*^*P*^ − 1 inter-PAS positions this segment contains, a null indel history could occur independently of those in other positions. Let 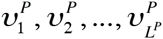 be the ancestries assigned to the *L*^*P*^ PASs in this segment (in spatial order). Focusing on the segment, we have 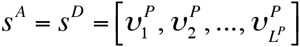. Then, excluding the contributions from both ends, the total multiplication factor for this segment is expressed as:

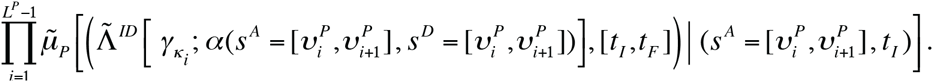

Here 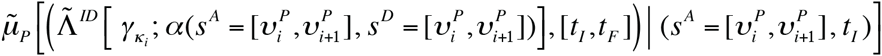 is the total multiplication factor for case (i) with 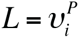 and 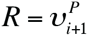. Under a space-homogeneous model, we can further simplify the above product as: 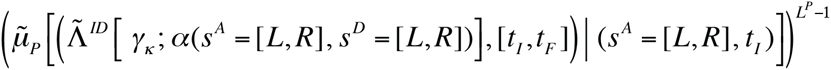. Here we used the uniform multiplication factor,

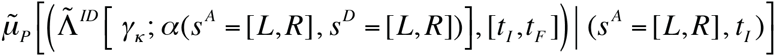

In case (ii), we assume that the ancestral state has ∈*L*^*A*^ sites in between the flanking PASs. Thus, the ancestral and descendant states could be represented as [*s*^*A*^ Δ*L*, 1,…, Δ*L*^*A*^, *R*] and *s*^*D*^ = [*L*, *R*], respectively. As long as 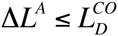, *N*_min_ [γ_k_; α(*s*^*A*^, *s*^*D*^)]= 1, and there is only one fewest-indel history, 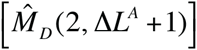 which consists of a single event that deletes the ancestral sites in between the PASs. Therefore, the contribution by the fewest-indel history is:

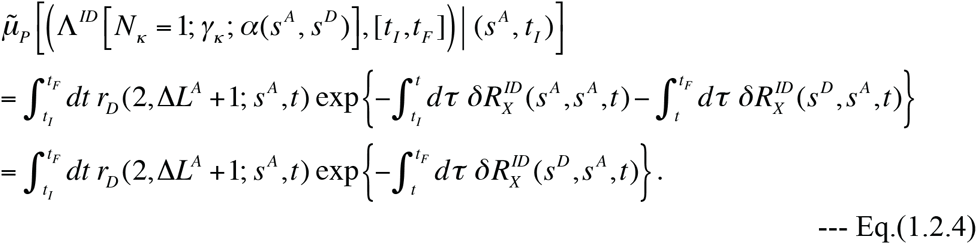

Each next-fewest indel history is composed of two indel events. There are two types. (a) Two successive deletions, 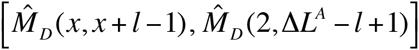 with *l* = 1,…, ∈*L*^*A*^ − 1 and *x* = 2,…, ∈*L*^*A*^ − *l* ∈ 2. And (b) an insertion followed by a deletion, 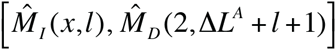 with *l* = 1,…, 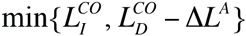 and *x* = 1,…,∈ *L*^*A*^ ∈1. Thus, in this case, the total contribution by the next-fewest indel histories is given by:

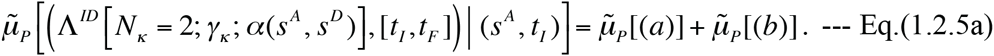

Here,

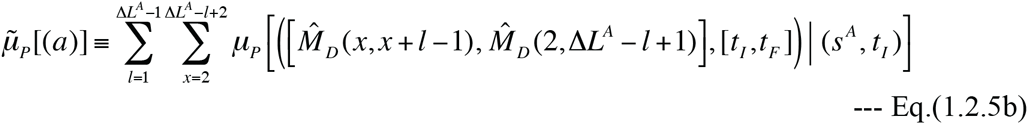

is the sum of contributions from the histories of type (a). And

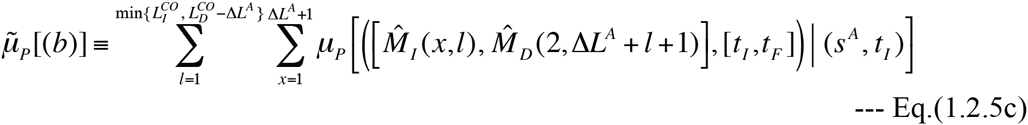

is the sum of contributions from the histories of type (b). Let 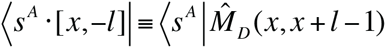 be the intermediate state in each type (a) history. Then, the history’s contribution is calculated as:

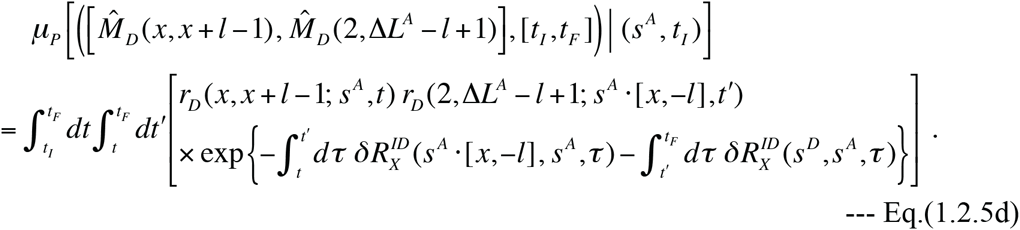

Similarly, let 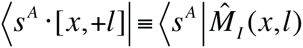 be the intermediate state in each type (b) history. Then, the history’s contribution is calculated as:

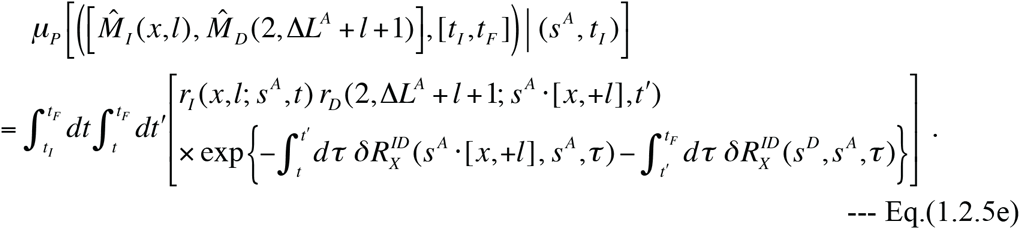

Eq.(1.2.4) and Eqs.(1.2.5a-e) can indeed be calculated at least numerically once the specific functional forms of the indel rates and the exit rates are given. For example, under Dawg’s indel model (Eqs.(2.4.4a,b) in part I) or the “long indel” model as noted in case (i), we have 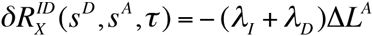, and Eq.(1.2.4) becomes:

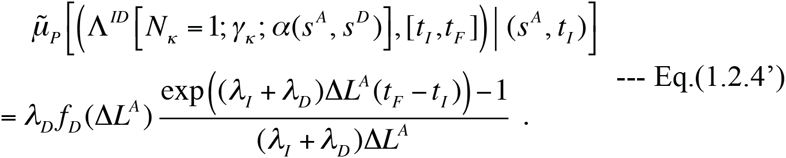

Similarly, using 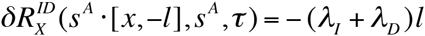 and 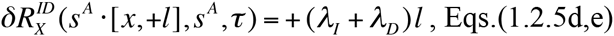 in Dawg’s model (and also in the “long indel” model) are calculated, respectively, as:

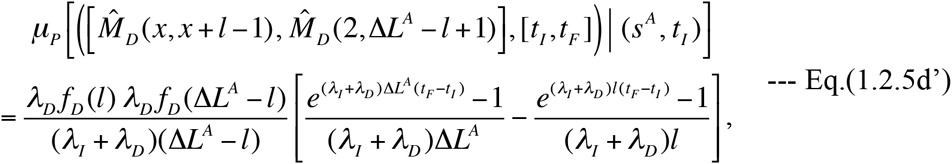

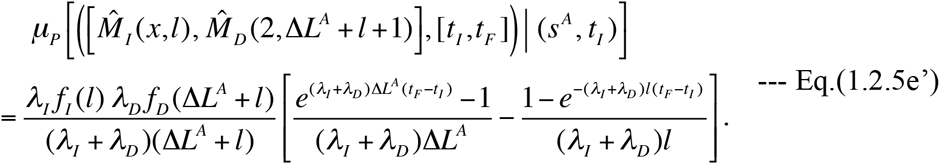

Substituting Eqs.(1.2.5d’,e’) into Eqs.(1.2.5a,b,c), we can concretely calculate the total contribution of the next-fewest-indel histories. Figure 3 A shows the ratio of the total next-fewest-indel contribution, Eq.(1.2.5a), to the fewest-indel contribution, Eq.(1.2.4), as a function of the number of deleted ancestral sites (∈*L*^*A*^) and the expected number of indels per site ((λ_*I*_ + λ_*D*_)(*t*_*F*_ − *t*_*I*_)). For the figure, we used the following parameters: 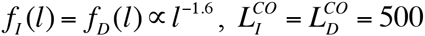, and λ_I_/λ_D_ = 1. The power-law behavior of the indel length distributions broadly conforms to the past empirical observations (*e.g*., Gonnet et al. 1992; Benner et al. 1993; Gu and Li 1995; Kent et al. 2003; Zhang and Gerstein 2003; Chang and Benner 2004; Yamane et al. 2006; Fan et al. 2007). Here, the exact overall indel rate (λ_*I*_ + λ_*D*_) does not matter, because the probabilities are invariant under the simultaneous rescaling of the rate and the time interval (*t*_*F*_ − *t*_*I*_) that keeps the expected number of indels per site (*i.e.*, (λ_*I*_ + λ_*D*_)(*t*_*F*_ − *t*_*I*_)) unchanged. Moreover, we confirmed that the results remain almost the same even if we use 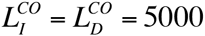. Now, as indicated by Figure 3A, each curve for a fixed (λ_*I*_ + λ_*D*_)(*t*_*F*_ −*t*_*I*_) reaches an asymptotic value slightly above 1 when Δ*L*^*A*^ is sufficiently large. Thus, to define a threshold within which the fewest-indel histories alone are likely to give a decent approximation of the probability, using the point at which the ratio is 1 (unity) is risky. Here, we tentatively define the threshold as the value of Δ*L*^*A*^ at which the ratio is 0.5. With this definition, the tentative threshold, 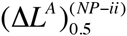, is around 128, 31, 12, 6 and 3 when (λ_I_ + λ_*D*_)(*t*_*F*_ − *t*_*I*_) is 0.01, 0.04, 0.1, 0.2 and 0.4, respectively (Table 1). Hence, we have a rough inversely proportional relationship: 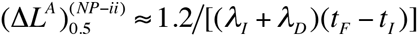, under the parameter setting used here.

**Table 1.** Various “threshold gap lengths” in case (ii) of local PWAs.

| $X = (\lambda_I + \lambda_D)(t_F - t_I)^a$ | $(\Delta L^A)_{0.5}^{(NP-ii)}{}^b$ | $(\Delta L^A)_{0.5}^{(1-ii)}{}^c$ | $(\Delta L^A)_{0.5}^{(2-ii)}{}^c$ | $(\Delta L^A)_{0.5}^{(5-ii)}{}^c$ |
| --- | --- | --- | --- | --- |
| 0.01 indels/site | 128 | 160 | > 300 | > 300 |
| 0.04 indels/site | 31 | 41 | 99 | 272 |
| 0.1 indels/site | 12 | 17 | 42 | 119 |
| 0.2 indels/site | 6 | 8 | 22 | 66 |
| Approximate relation<sup>d</sup> | $Y \approx 1.2 / X$ | $Y \approx 1.6 / X$ | $Y \sim 4 / X$ | $Y \sim 11 / X (?)$ |

NOTE: The parameters used for this analysis are: $\lambda_I = \lambda_D = 0.1$ , $L_I^{CO} = L_D^{CO} = 500$ , and $f_I(l) = f_D(l) = l^{-1.6} / \left( \sum_{k=1}^{500} k^{-1.6} \right)$ . For the iteration analysis, we used
$\Delta L_{\max}^A = \Delta L_{\max}^D = 300$ and $(\lambda_I + \lambda_D)(t_F - t_I)/N_P = 0.002$ as well. Because of the symmetry under the time reversal, the identical results apply also to the local PWAs in case (iii), if we replace $(\Delta L^A)_{0.5}^{(xx-ii)}$ with $(\Delta L^D)_{0.5}^{(xx-iii)}$ .
<sup>a</sup> The expected number of indels per site.
<sup>b</sup> The number of ancestral sites in between the PASs, *i.e.*, $\Delta L^A$ , at which the total probability of the next-parsimonious local indel histories is 1/2 (=0.5) of that of the parsimonious local indel histories.
<sup>c</sup> $(\Delta L^A)_{0.5}^{(N_D-ii)}$ is the value of $\Delta L^A$ at which the local histories involving up to (and including) $N_D$ indels each account for 1/2 (=0.5) of the total probability of all local histories consistent with the configuration. We used the probability calculated up to (and including) $N_D = 50$ indel steps as a proxy of the total probability.
<sup>d</sup> A rough (inversely proportional) relationship between each threshold gap length ( $Y$ ) and the expected number of indels per site ( $X$ ).

**Figure 3.**
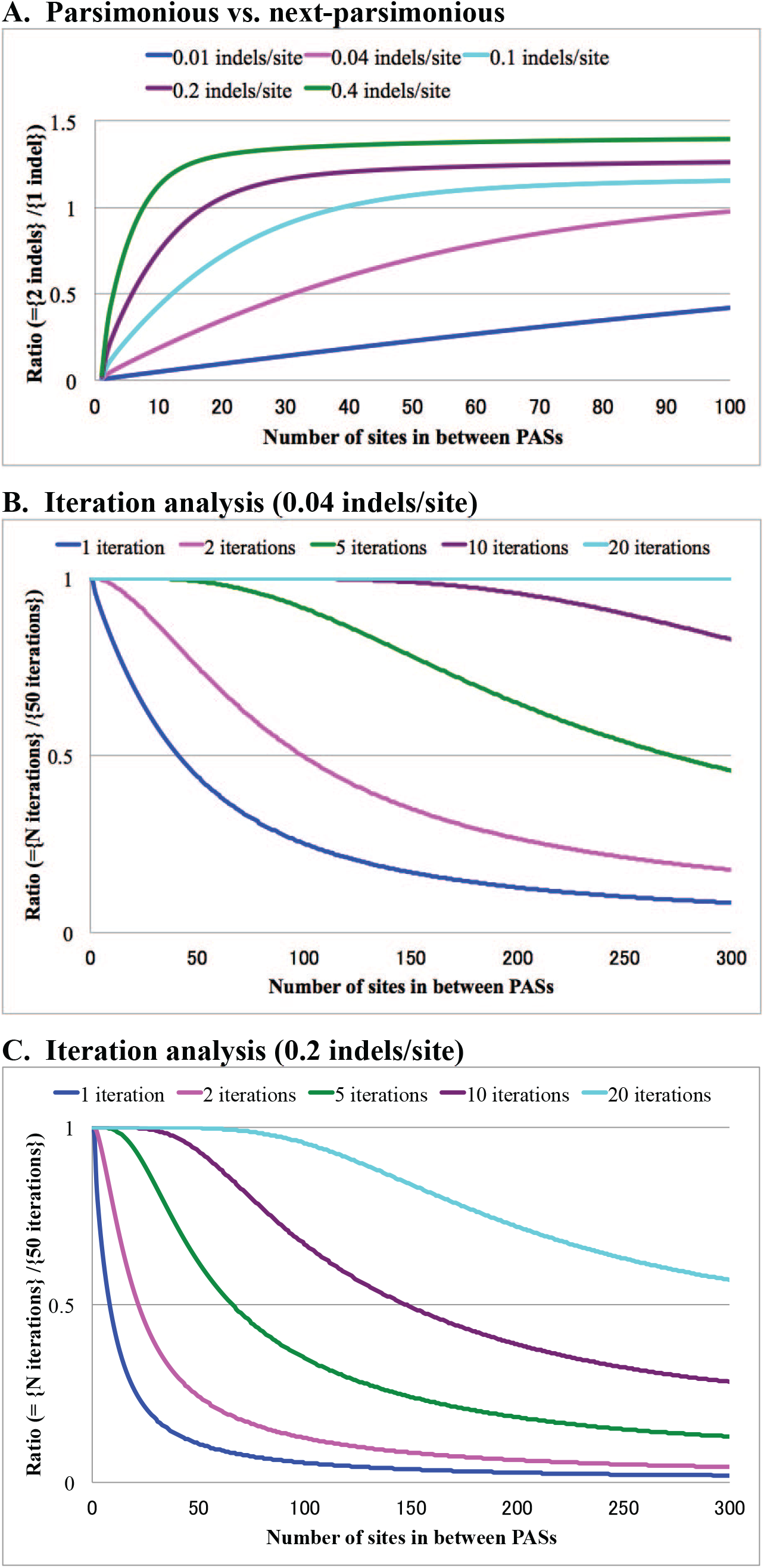
Perturbation analyses on local PWA probabilities. The results shown in all panels of this figure apply to both case (ii) and case (iii) local PWAs. In all panels, the abscissa is Δ*L*^*A*^ in case (ii) and Δ*L*^*D*^ in case (iii). Panel **A** shows the ratio of the total probability of the next-fewest-indel histories to that of the fewest-indel history. The blue, magenta, cyan, purple and green curves show the ratios when the expected numbers of indels per site, *i.e*., the values of (*λ*_*I*_ + *λ*_*D*_)(*t*_*F*_ – *t*_*I*_), are 0.01, 0.04, 0.1, 0.2 and 0.4, respectively. Panels **B** and **C** show the ratios of the multiplication factors calculated *via* the iteration formulas (either Eq.(1.2.8) or Eq.(A1.3.3) in Appendix A1), including histories with up to *N*_*ID*_ = 1 (blue), 2 (magenta), 5 (green), 10 (purple), and 20 (cyan) indels, to the factor including histories with up to *N*_*ID*_ = 50 indels. Panel B is for (*λ*_*I*_ + *λ*_*D*_)(*t*_*F*_ – *t*_*I*_) = 0.04 indels per site, and panel C is for (*λ*_*I*_ + *λ*_*D*_)(*t*_*F*_ – *t*_*I*_) = 0.2 indels per site. The following parameters were used for all panels: *f*_*l*_(*l*) = *f*_*D*_(*l*) ∝ *l*^−1.6^, 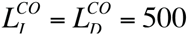 and *λ*_*I*_/*λ*_*D*_ = 1. For panels B and C, we also set the following parameters: 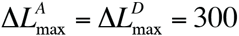 and (*λ*_*I*_ = *λ*_*D*_)(*t*_*F*_ – *t*_*I*_)/*N*_*P*_ = 0.002.

In case (iii), we assume that the descendant state has Δ*L*^*D*^ sites in between the flanking PASs. Thus, the ancestral and descendant states could be represented as 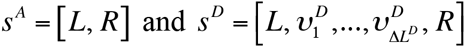, respectively. The ancestries satisfy 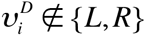 for every *i* = 1,…, Δ*L*^*D*^, and 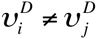 for every pair (*i*, *j*) with *i* ≠ *j*, and their details depend on the responsible indel history. (Actually, such details don’t matter in the state space *S*^*II*^, as explained in Subsection 2.1 of part I.) As long as 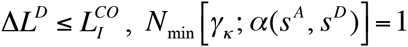, and there is only one fewest-indel history, 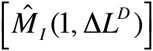. The history consists of a single event that inserts the descendant sites in between the PASs. As in case (ii), each next-fewest indel history is composed of two indel events, and classified into two types. (c) Two successive insertions, 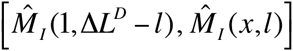 with *l* = 1,…, Δ*L*^*D*^ − 1 and *x* = 1,…, Δ*L*^*D*^ − *l* +1. And (d) an insertion followed by a deletion, 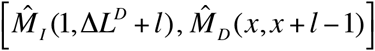 with *l* = 1,…, 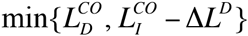 and *x* = 2,…, Δ*L*^*D*^ + 2. The sum of contributions by the fewest-indel histories and that by the next-fewest-indel histories can be calculated as in case (ii). And their calculations are detailed in Appendix A1.1. If calculated under the same setting as used for Figure 3 A and with the same value of (λ_*I*_ + λ_*D*_)(*t*_*F*_ − *t*_*I*_), their ratio with *L*^*D*^ = *L* is identical to that in case (ii) with Δ*L*^*A*^ = *L*, because of the symmetry of the probabilities under the time reversal. Thus, the same conclusions can be drawn from Figure 3 A also on this case.

In case (iv), we assume that the ancestral and the descendant states have Δ*L*^*A*^ and [*L*^*D*^ sites, respectively, in between the flanking PASs. Thus, the ancestral and descendant states could be represented as *s*^*A*^ = [*L*, 1,…, Δ*L*^*A*^, *R]* and 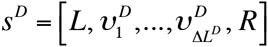, respectively. Here, the descendant ancestries satisfy 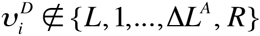 for every *i* = 1,…, Δ*L*^*D*^, and 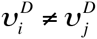 for every pair (*i*, *j*) with *i* ≠ *j*, and their details depend on the responsible indel history. (Again, the details don’t matter in the space *S*^*II*^.) As long as 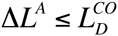 and 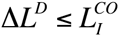, *N*_min_ γ_*k*_; α(*s*^*A*^, *s*^*D*^) 2. As indicated by Eqs.(A1.3c’,d’) in Appendix A1 of part I, there are three types of fewest-indel histories. (e) The deletion of the ancestral sites followed by an insertion of Δ*L*^*D*^ sites, 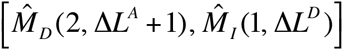. (f) An insertion immediately on the right of the ancestral sites to be deleted, followed by the deletion, 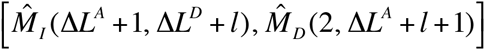 with 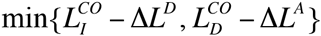. And (g) an insertion immediately on the left of the ancestral sites to be deleted, followed by the deletion, 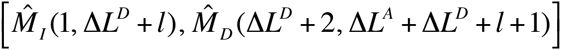 also with *l* = 0,…, 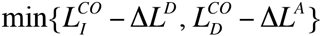. In this case, each next-fewest-indel history is composed of three indel events, and classified into one of 6 broad types: (h) two successive deletions followed by an insertion; (i) a deletion, followed by an insertion, followed by a deletion; (j) an insertion followed by two successive deletions; (k) a deletion followed by two successive insertions; (l) an insertion, followed by a deletion, followed by an insertion; and (m) two successive insertions followed by a deletion.

And these six broad types can be further sub-classified into 24 sub-types, as described in Appendix A1.2. The calculations of the sum of contributions from the fewest-indel local histories and that from the next-fewest-indel local histories are also detailed in Appendix A1.2. Table 2 shows their ratios calculated for some typical configurations under the same setting as for Figure 3 A. As the table indicates, the approximation by the fewest-indel histories alone is likely to be decent as long as the expected number of indels (*i.e*., (λ_*I*_ + λ_*D*_)(*t*_*F*_ − *t*_*I*_)) and the gap lengths (i.e., Δ*L*^*A*^ and Δ*L*^*D*^) are at most moderate.

**Table 2.** Perturbation analysis on local PWA probabilities in case (iv)

| $(\Delta L^A, \Delta L^D)$ | 0.01 indels/site | 0.04 indels/site | 0.1 indels/site | 0.2 indels/site |
| --- | --- | --- | --- | --- |
| (1, 1) | **0.003** | **0.010** | **0.024** | **0.045** |
| (3, 1) | **0.021** | **0.084** | **0.204** | **0.393** |
| (3, 3) | **0.042** | **0.166** | **0.402** | 0.768 |
| (5, 5) | **0.073** | **0.283** | 0.672 | 1.256 |
| (10, 1) | **0.064** | **0.246** | 0.572 | 1.013 |
| (10, 10) | **0.149** | 0.561 | 1.292 | 2.288 |
| (25, 1) | **0.151** | 0.547 | 1.112 | 1.541 |
| (25, 4) | **0.198** | 0.723 | 1.519 | 2.234 |
| (30, 10) | **0.288** | 1.038 | 2.164 | 3.072 |
| (100, 1) | 0.537 | 1.333 | 1.507 | 1.574 |
| (100, 3) | 0.607 | 1.593 | 1.894 | 2.033 |
| (300, 1) | 1.165 | 1.394 | 1.427 | 1.527 |

NOTE: Each cell shows the ratio of the total probability of the next-fewest indel histories to that of the fewest indel histories, when there are $\Delta L^A$ ancestral sites and $\Delta L^D$ descendant sites in between the PASs. Each column is labeled with the expected number of indels per site (*i.e.*, $(\lambda_I + \lambda_D)(t_F - t_I)$ ). The parameters used for this analysis are: $\lambda_I = \lambda_D = 0.1$ , $L_I^{CO} = L_D^{CO} = 500$ , and $f_I(l) = f_D(l) = l^{-1.6} / \left( \sum_{k=1}^{500} k^{-1.6} \right)$ .
Because of the symmetry of the probabilities under the time reversal, the ratio for $(\Delta L^A, \Delta L^D) = (L_1, L_2)$ is identical to that for $(\Delta L^A, \Delta L^D) = (L_2, L_1)$ . Thus we only showed the results for $\Delta L^A \geq \Delta L^D$ . The ratios that are less than 0.5 are shown in boldface.

It is difficult to calculate the summed contributions from local histories involving more indels, especially in case (iv). We could calculate the contribution from a single local history involving any number of indels if we use the algorithm for a “trajectory likelihood” given by Miklós et al. (2004). As we exemplified in Appendix A1.2, however, it is already quite hard to enumerate all possible local indel histories even up to the next-to-minimum number of involved indels. Nevertheless, if we consider only cases (i), (ii), and (iii) under a (locally) spatially homogeneous model, we can work out systems of “exact” integral equations that could in principle provide the numerical solutions for the total sum of contributions to a multiplication factor up to a desired level of accuracy (at some time and memory expenses). In the following, we derive a system of integral equations to give multiplication factors for cases (i) and (ii). Another system of integral equations, which gives multiplication factors for cases (i) and (iii), is derived in Appendix A1.3.

Here, we assume that the indel rates are locally homogeneous, which means that the rates do not depend on the exact positions that the indels hit, *as long as* they are confined in the region that accommodates the local history. Thus, we assume that the indel rate is *locally* homogeneous and the exit rate is *locally* an affine function of the (local) sequence length, but that they may be non-homogeneous *globally*. (In terms of equations, we *locally* assume Eqs.(5.1.1a,b) of part I for the indel rates and Eq.(5.2.4) of part I for the *local* exit rate, but we assume something like Eqs.(5.3.2a,b,c) of part I for the *global* exit rate.) We are now considering only cases (i) and (ii), in which (local) ancestral and descendant states should be *s*^*A*^ = [*L*, 1,…, Δ*L*^*A*^, *R]* and *s*^*D*^ = [*L*, *R*], respectively, with Δ*L*^*A*^ = 0,1, 2,…. Because ofthe local homogeneity, Eqs.(V-1.1a,b), the exit rate 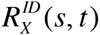 of a state *s* (∈ *S*) in this context depends only on the (local) sequence length, *L*(*s*) = 2 + Δ*L*(*s*). Thus, ∈*L*(*s*) adequately represents the local sequence state *s*, and we let 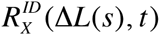 denote its (local) exit rate. The starting point of the equation system is the fundamental integral equation (Eq.(3.1.4) of part I) for the stochastic indel evolution operator 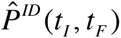. *P*We sandwich the fundamental integral equation with 〈 *s*^*A*^ | and *s*^*D*^ 〉, and expand the instantaneous mutation operator 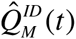 using its definition (*i.e*., Eq.(3.1.1c) of part I supplemented by Eqs.(2.4.2b’,c’) of part I). Because we know that the flanking PASs, which are assigned the ancestries *L* and *R*, have never been struck by any indels, we can ignore the effects of indels that hit the PASs. And, because we are now focusing on the local alignment, we will also ignore the indels completely outside of the region delimited by the PASs. Thus, we have:

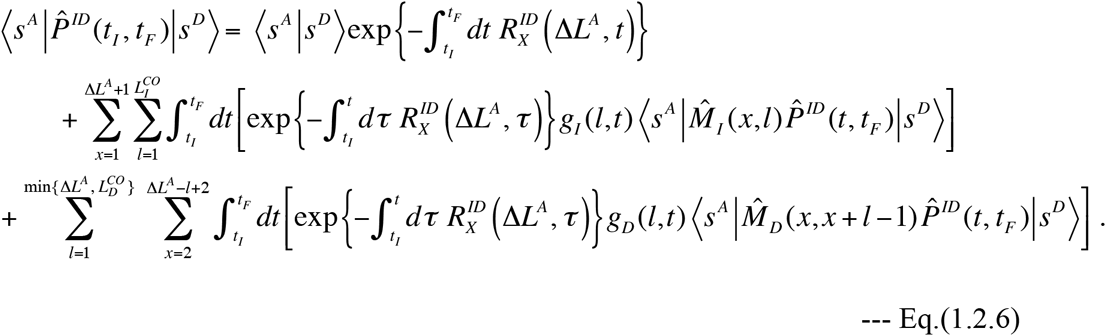

In the present setting, the number of sites between the PASs, ∈*L*(*s*), uniquely determines the local sequence state *s*. Thus, we let the local states denoted as 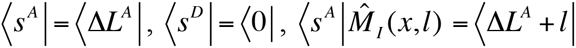, and 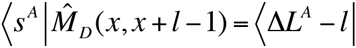. We also introduce the notation, 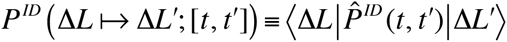, for the probability that the local state with Δ*L* sites in between the PASs at time *t* became a state with Δ*L*′ sites in between at time *t*‵. Then, taking advantage of the independence of the indel rates, exit rates and *P*^*ID*^ (∈*L* ↦ Δ*L*‵); [*t*, *t*‵] on the position of each indel (*x*), we have:

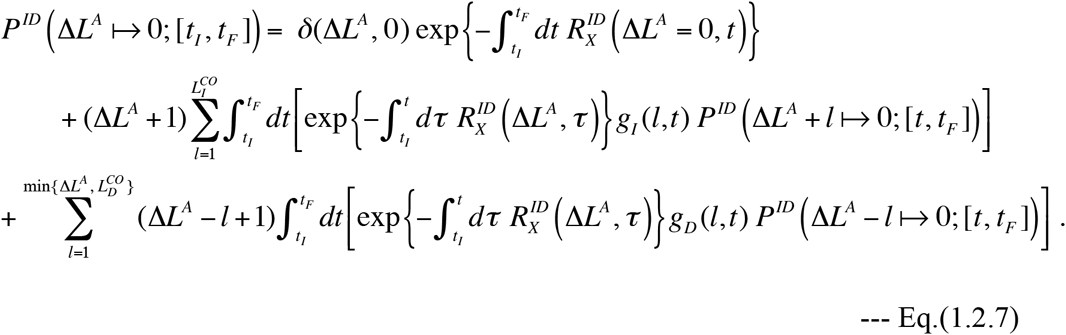

Here δ(Δ*L*, Δ*L*‵) is Kronecker’s delta, which equals 1 if Δ*L* ‵Δ*L*‵, and 0 if Δ*L* = *L*‵∈. Eq.(1.2.7) gives the desired system of integral equations for the “exact” probabilities, *P*^*ID*^(Δ*L*^*A*^ ↦ 0; [*t*, *t*]), with non-negative integers Δ*L*^*A*^ = 0,1, 2,…. This equation holds for every non-negative integer Δ*L*^*A*^ and even if we replace the initial time *t*_I_ with any time in the closed interval [*t*_*I*_, *t*_*F*_]. Thus, the equations can be solved iteratively, starting with the “zero-event approximation” of the probability, 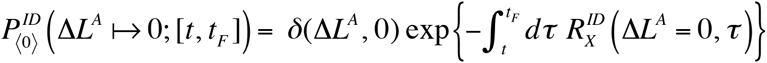, and calculating the approximation at the *n*_*S*_ th step, 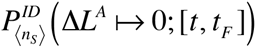, from the approximation at the previous step via the recursion relation:

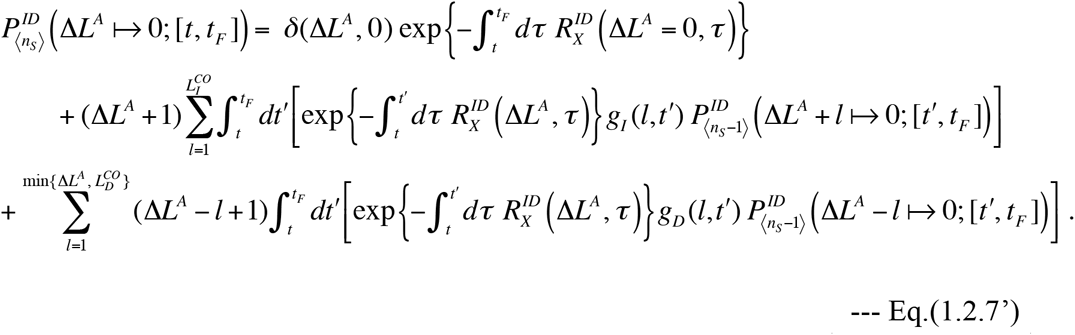

If *N*_*ID*_ iteration steps are performed, the resulting probability, 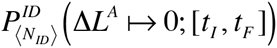, is the summation of the probabilities over all possibly responsible local histories consisting of up to (and including) *N*_*ID*_ indel events. After the iteration is finished, the multiplication factor will be obtained by following its definition (*i.e*., Eq.(4.1.1b) in part I). We have:

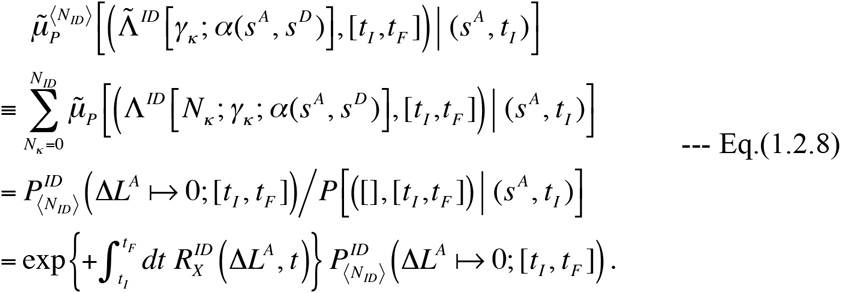

The accuracy of the numerical solutions will depend on how finely we partition the time interval [*t*_*I*_, *t*_*F*_]. If the interval is partitioned into *N*_*P*_ equal-sized sub-intervals, we could in principle achieve an accuracy of *O* ((*N*_*P*_)^4^) under Simpson’s rule (*e.g*., Press et al. 1992). However, as the number of sub-intervals increases, it would take longer to complete the calculation. A naïve implementation of the aforementioned numerical iteration would have the time complexity of *O* (*N* (*L*^*CO*^)^2^ (*N*)^2^) and the space complexity of *O* (*L*^*CO*^ *N*), when we want to obtain the total probabilities of local histories composed of up to *N*_*ID*_ indels each, with 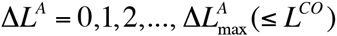. Here we set 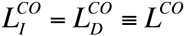. This becomes impractically slow when either *L*^*CO*^ or *N*_*P*_ is large, *e.g*., around 1000 or greater. It is likely that *N*_*P*_ does not have to be this large, as it would be usually enough to set *N* around 100 or smaller. However, *L*^*CO*^ will often be around 1000 or greater, indeed making the naïve algorithm too slow to be practical. Fortunately, we can avoid this problem by rewriting the recursion equation, Eq.(1.2.7’), as:

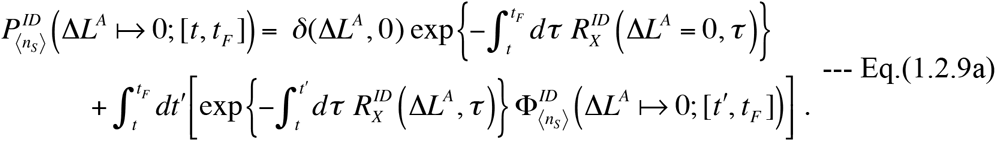

Here, the “auxiliary function,” 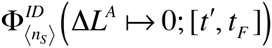 is given by:

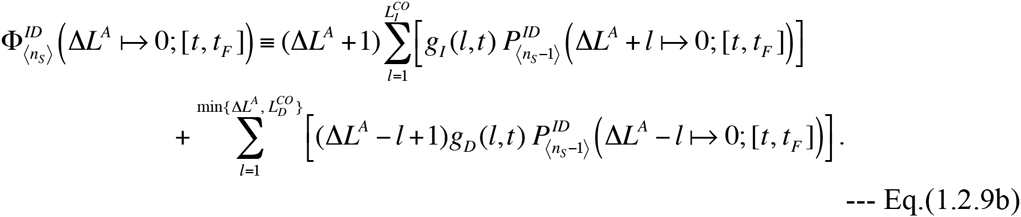

Consider the following “two-sub-step” algorithm. In the first sub-step (in each iteration step), it calculates 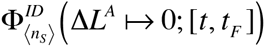’s via Eq.(1.2.9b) and stores them for all Δ*L*^*A*^ = 0,1, 2,…, *L*^*CO*^ at all time points, 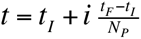 with *i* = 0,1,…, *N*_*P*_. And, in the second sub-step, it uses them to calculate the probabilities 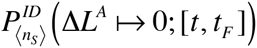 via Eq.(1.2.9a) for the same values of Δ*L*^*A*^ and *t*. This algorithm can reduce the time-complexity to *O* (*N L*^*CO*^ (*L*^*CO*^ ∈ *N*)*N*) while keeping the space complexity to be *O* (*L*^*CO*^ *N*). This algorithm does finish in a practical amount of time (typically on the order of an hour or shorter when implemented in Perl). But it may still be too slow to perform each time we evaluate the probability of the gap configuration of a local MSA. Good news is that a single run of the iteration algorithm *inevitably* calculates the probabilities for all 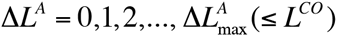 at all temporal partitioning points, 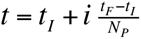, as well as at *t* = *t*_*I*_ and *t* = *t*_*F*_. Thus, once we calculate the probabilities with a fixed set of model parameters, we could use them to calculate the probabilities of various alignments (under various phylogenetic trees), as long as the model parameters remain unchanged. In any case, the time and space complexities might be further reduced without considerably compromising the accuracy by a clever beforehand discarding of terms that are unlikely to make significant contributions to the final probabilities. Panels B and C of Figure 3 show the ratios of the multiplication factors, Eq.(1.2.8) at *N*_*ID*_ = 1, 2, 5, 10, 20 iteration steps, to that at *N*_*ID*_ = 50 steps. They were calculated with 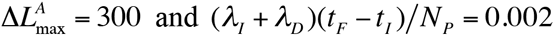, and under the same setting as that used for Figure 3 A. (We confirmed that results remain almost the same even if *L*^*CO*^ is greater than 1000, instead of *L*^*CO*^ = 500 used for the figure.) When *N*_*ID*_ ≥ 2, we actually started from *N*_*ID*_ = 2, at which the probabilities were calculated using Eqs.(1.2.2a,b’), Eq.(1.2.4’) and Eqs.(1.2.5a,b,c,d’,e’), in stead of from *N*_*ID*_ = 0 as mentioned above, in order to enhance the accuracy of the approximation. As indicated by the panels B and C, the accuracy of the probabilities improves in a step-wise manner as the number of iterations increases. Now that the “exact” probabilities are available in cases (i) and (ii), we can use them to define more precise threshold values of Δ*L*^*A*^ within which the approximate probabilities are expected to be quite accurate. For example, let 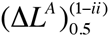 be the value of Δ*L*^*A*^ at which the approximation by the 1-event local indel history alone accounts for 50% of the total probability of the local PWA. As shown in Table 1, 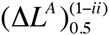 is slightly larger than the aforementioned “tentative threshold,” 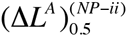. Indeed, we can observe a rough inversely proportional relationship, 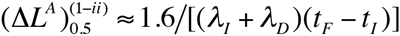. Thus, we expect that the tentative threshold 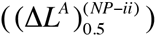 should work as a slightly conservative criterion for the decency of the approximation by the fewest-indel histories alone. We could also define the threshold, 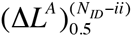, at which the probability calculated up to (and including) *N*_*ID*_ iterations, *i.e*., 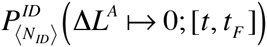, is 1/2 (=0.5) of the “exact solution.” As indicated by Table 1, 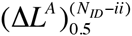 is over *N*_ID_ times as large as 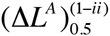. This implies that, even when the fewest-indel histories alone give a poor approximation of the probability, taking account of the next-fewest-indel histories, or of the histories involving yet more indels, will considerably expand the range of Δ*L*^*A*^ where the approximate probability is quite accurate.

Following the similar procedures, this time starting from the integral equation, Eq.(III-1.2), we can also derive a system of integral equations for the multiplication factors for cases (i) and (iii), as described in Appendix A1.3. Thanks to the symmetry of the probabilities under the time reversal, panels B and C of Figure 3 can also be interpreted as the results of numerical calculations of this equation system, under the same setting as above except that 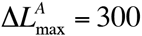 is replaced by 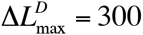.

### 1.3. Perturbation analyses on local MSA

Compared to contributions to local PWAs, those to local MSAs are much more complex. In this subsection, we consider some simple but common patterns, under a tree *T* with three OTUs, corresponding to the external nodes, *n*_1_, *n*_2_ and *n*_3_. Here, we regard its single internal node as the root node *n*^*Root*^ for simplicity (panel A of Figure 4). Let *b*_*m*_(*m* = 1, 2, 3) be the branch that connects the nodes *n*^*Root*^ and *n*_m_. Let *s*_*m*_ ∈ *S* (*m* = 1, 2, 3) be the (local) sequence state at node *n*_*m*_. Then, we consider the gap-configurations of the MSAs of the three sequences, α[*s*_1_, *s*_2_, *s*_3_], as well as the consistent sequence states *s*^*Root*^ ∈ *S* at the root node *n*^*Root*^. As in the previous subsections, we focus on the portion of MSAs delimited by a pair of PASs, whose ancestries are denoted as *L* and *R*. Here we consider four typical situations (see Figure 4; see also Subsection 3.4 of part I (Ezawa, Graur and Landan 2015a) for complexities concerning this issue). (I) None of {*s*_1_, *s*_2_, *s*_3_} have any site in between the PASs (Figure 4, panel B). (II) *s*_1_ and *s*_2_ share the identical set of sites in between the PASs, but *s*_3_ has no site in between (panel C). (III) *s*_1_ has a set of sites in between the PASs, but neither *s*_2_ nor *s*_3_ has even a single site in between (panel D). And (IV) *s*_1_ has a set of sites in between the PASs, but *s*_3_ has no site in between, and *s*_2_ lacks a run of some, but not all, of contiguous sites of *s*_1_ in between the PASs (panel E).

**Figure 4.**
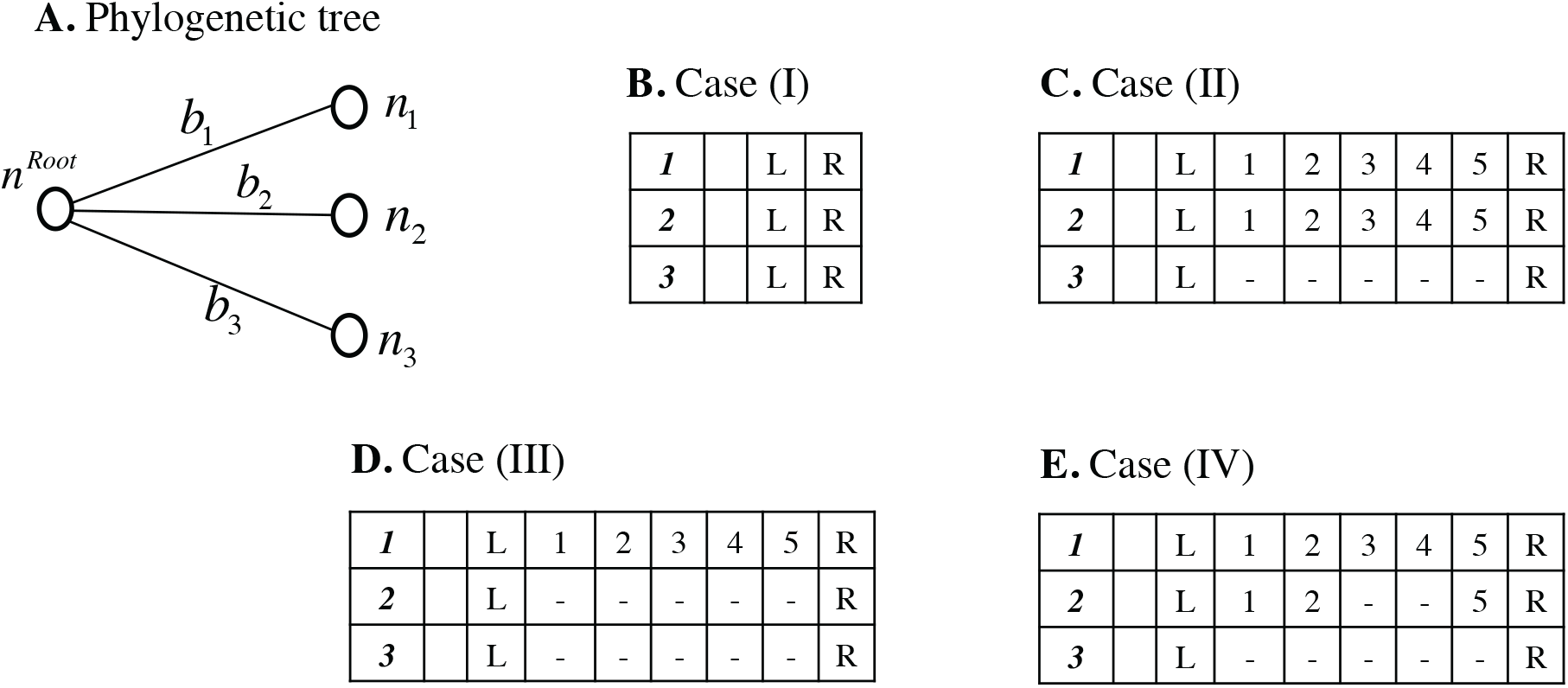
Typical local gap configurations in MSAs considered for perturbation analyses in Subsection 1.3 of Results. **A.** The 3-OTU tree used in the perturbation analyses and the notation. A node (open circle) is labeled *n*_*i*_ (external) or *n*^*Root*^ (root). A branch is labeled *b*_*i*_. **B.** Local gap configuration in case (I). **C.** Case (II) with Δ*L*^*D*12^ = 5. **D.** Case (III) with Δ*L*^*D*1^ = 5. **E.** Case (IV) with Δ*L*^*D*1^ = 5, *i* = 2 and *j* = 5. In this figure, none of the sites are colored, to take account of the possibilities of their preservations, insertions and deletions.

These situations are not restricted to the 3-OTU trees but widely applicable to each portion surrounding a trivalent node of any tree topology, although they never exhaust all gap configurations. The time at *n*^*Root*^ will be represented as *t*_I_, and the time at node *n*_*m*_ will be represented as *t*_*F*:*m*_. The indel parameters along branch *b*_*m*_ will be indicated by the subscript “:*m*.”

Case (I) is represented by the external sequence states *s*_1_ = *s*_2_ = *s*_3_ = [*L*, *R*]. In this case, we have *N*_min_ [*C*_k_; α[*s*_1_, *s*_2_, *s*3]; *T*] = 0. And the set of fewest-indel local histories, 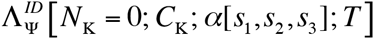, is composed only of a no-indel history:

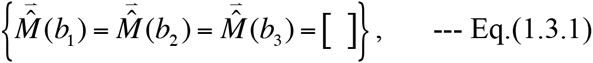

with 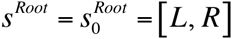. Thus, according to Eq.(1.1.2b), supplemented by Eqs.(4.2.4b,6b,8) of part I, the portion of the multiplication factor contributed by the fewest-indel local history is:

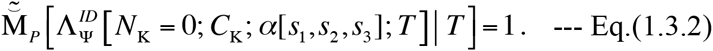

Here we used 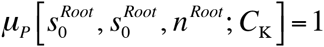. No single-event local history can result in the gap configuration in this case. Each next-fewest-indel history consists of two indels, and it can be represented as:

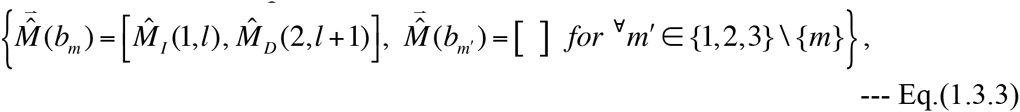

with 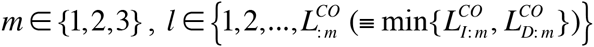, and with 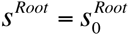 again. Thus, the total contribution to the multiplication factor by the next-fewest-indel histories can be calculated similarly to Eqs.(1.2.2a,b). We have:

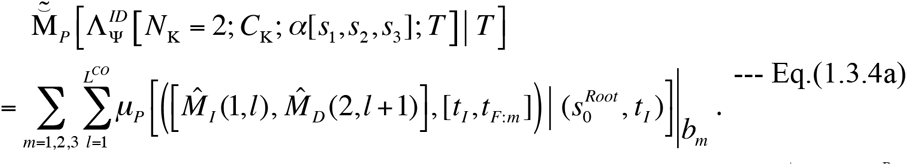

Each summand can be calculated from Eq.(1.2.2b), by replacing *s*^*A*^ with 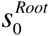 and also replacing the time and rate parameters with those assigned to each branch. Especially, under Dawg’s model, each summand is calculated as:

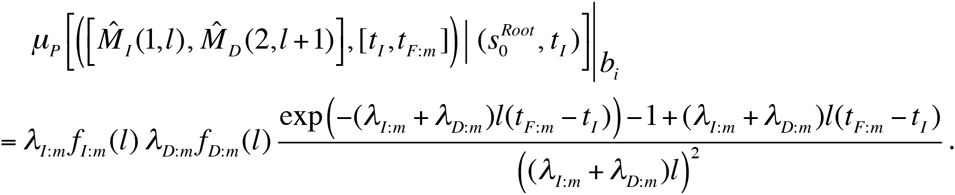

---Eq.(1.3.4b)

If the three branches share the same time interval and the indel rate parameters, the summed contribution from the next-fewest-indel histories, Eq.(1.3.4a), is exactly three times Eq.(1.2.2a) for a PWA. Indeed, this total contribution on a general tree can be calculated by summing Eq.(1.2.2a) (with appropriate parameters) over all branches of the tree. Following the same line of reasoning as around Eq.(1.2.3), this total contribution is roughly proportional to the summation of the squared branch lengths over all branches. This means that a richer sampling of the homologous sequences will not significantly increase, or might rather slightly decrease, this total contribution, as long as the maximum evolutionary distance remains at a similar level. Incidentally, any root sequence state of the type *s*^*Root*^ = [*L*, 1,…, Δ*L*^*Root*^, *R]* is also consistent with α[*s*_1_, *s*_2_, *s*_3_] in this case. Such a state, however, would require at least three indels, in order to delete the extra sites, 1,…, Δ*L*^*Root*^, independently along the three branches. Thus, the contributions from the local indel histories with such root states would be smaller in general.

Case (II) is represented by the external sequence states *s* = *s* = [*L*, 1,…, Δ*L*^*D*12^, *R*] and *s*3 = [*L*, *R*]. In this case, the “phylogenetic correctness"condition (see, *e.g*., Chindelevitch et al. 2006; Section 3.4 of part I (Ezawa, Graur and Landan 2015a)) dictates that the root state *s*^*Root*^ must have all the sites with ancestries 1,…, Δ*L*^*D*12^. In this case, we have *N*_min_*C* _k_; α[*s*_1_, *s*_2_, *s*_3_]; *T* = = 1. And the set of fewest-indel local histories, 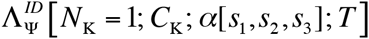, consists of a single element:

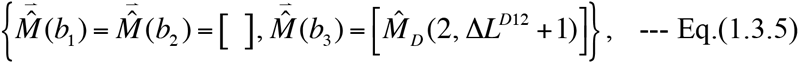

with 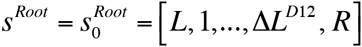. Again, according to Eq.(1.1.2b), supplemented by Eqs.(4.2.4b,6b,8) of part I, the contribution by the fewest-indel local history turns out to be exactly the same as Eq.(1.2.4) for case (ii) of PWAs, with the parameters replaced with those assigned to the branch *b*3, and with Δ*L*^^*A*^^ replaced with ∈*L.D*12 Especially, under Dawg’s model, we have:

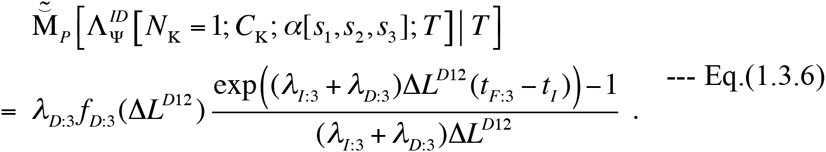

As in case (ii) of PWAs, each next-fewest-indel history is composed of two indel events, and there are two types of such histories. One is based on type (a) in case (ii):

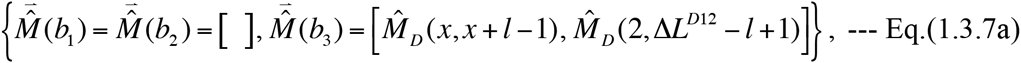

with *l* = 1,…, Δ*L*^*D*12^)1, *x* = 2,…, Δ*L*^*D*12^) *l* ∈ 2, and also with 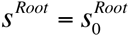. And the other is based on type (b) in case (ii):

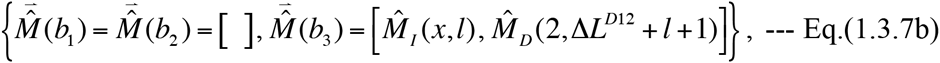

with 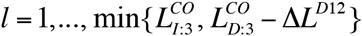, *x* = 1,…, Δ*L*^*D*12^ +1, and also with 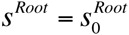 again. Thus the summed contributions over the next-fewest-indel histories is given by:

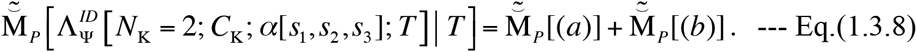

Here 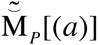 and 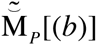 are given by exactly the same equations as Eqs.(1.2.5b,d) and Eqs.(1.2.5c,e), respectively, with the parameters replaced by those assigned to branch *b*3, and with Δ*L* replaced by Δ*L*. Especially, under Dawg’s model, the contributions by the individual next-fewest-indel histories are given by Eqs.(1.2.5d’,e’) with an appropriate replacement of the parameters and the ancestral state. Thus, panel A of Figure 3 can also be interpreted exactly as the comparison between the fewest-indel history’s contribution and the next-fewest-indel histories’ total contribution in the current case. Incidentally, root sequences with some additional ancestral sites in between the PASs of 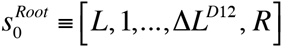 are also consistent with α[*s*_1_, *s*_2_, *s*_3_] in this case. However, such root sequence states require at least three indels each to give rise to α[*s*_1_, *s*_2_, *s*_3_]. This is because the additional ancestral sites need to be deleted independently along *b*_1_ and *b*_2_, even if they are deleted simultaneously with the sites with the ancestries 1,…, Δ*L*^*D*12^ along *b*_3_. Thus, in general, the indel histories consistent with such root states are expected to contribute much less to the multiplication factor.

Case (III) is represented by the external sequence states *s* = [*L*, 1,…, [*L*^*D*1^, *R*] and *s*_2_ = *s*_3_ = [*L*, *R*]. In this case, the phylogenetic correctness condition does not require the root state to have any site in between the PASs. Thus we have 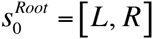. As in case (II), we have *N*_min_ *C*_k_; α[*s*_1_, *s*_2_, *s*_3_]; *T* = = 1. And the set of fewest-indel local histories, 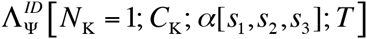, consists of a single element:

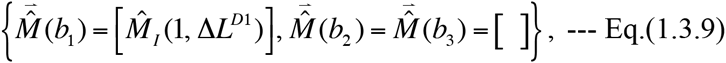

with 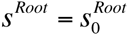. Again, as in case (II), the contribution by this local history turns out to be exactly the same as Eq.(A1.1.1) (in Appendix A1.1) for case (iii) of PWAs, with the parameters replaced with those assigned to the branch *b*_1_, and with Δ*L*^*D*^ replaced with Δ*L*^*D*1^. Especially, under Dawg’s model, we have:

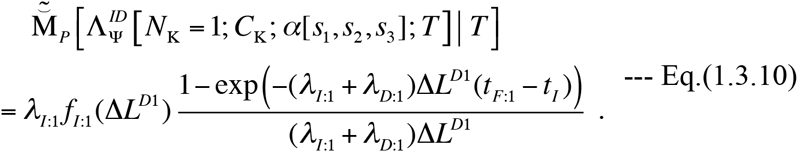

As in case (iii) of PWAs, each next-fewest-indel history is composed of two indel events. Unlike case (iii) of PWAs, however, there are three types of such histories; two of them are similar to those in case (iii), but the other one is totally new. Specifically, the first one is based on type (c) in case (iii):

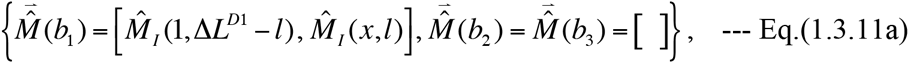

with *l* = 1,…, Δ*L*^*D*1^)1, *x* = 1,…, Δ*L*^*D*1^) *l* ∈1, and also with 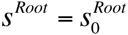. The second one is based on type (d) in case (iii):

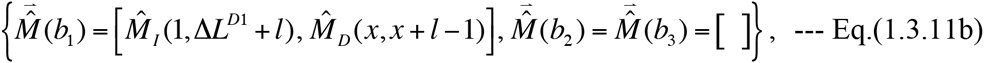

with 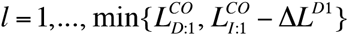, *x* = 2,…, Δ*L*^*D*1^ + 2, and also with 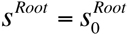 again. The third one involves events along *b*_2_ and *b*_3_, instead of along *b*_1_:

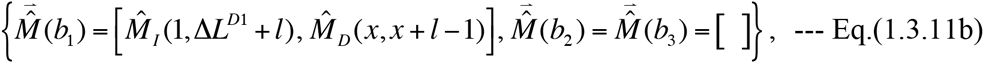

It is consistent with the root state *s*^*Root*^ = *s*_1_ = [*L*, 1,…, Δ*L*^*D*1^,*R*] instead of 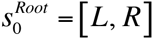. It should be noted that there is only one local history of the third type. In this case, therefore, the summed contribution of the next-fewest-indel local histories is given by:

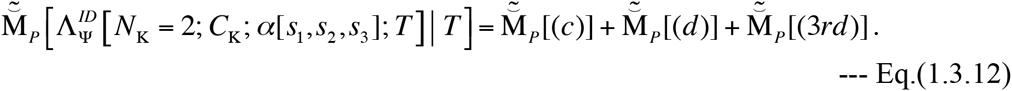

Here, 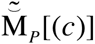 and 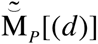 are the summed contributions of the type (c)-based and type (d)-based histories, respectively. They are given by exactly the same equations as Eqs.(A1.1.2b,d) and Eqs.(A1.1.2c,e), respectively, with the parameters replaced by those assigned to branch *b*_1_, and with Δ*L* replaced by Δ*L*^*D*1^. Under Dawg’s model, these two terms are given by summations of Eqs.(A1.1.2d’,e’) with an appropriate replacement of the parameters and the descendant states. Thus, panel A of Figure 3 can also be interpreted as the comparison between the total contribution of these two types of next-fewest-indel histories and the fewest-indel history’s contribution.

Meanwhile, 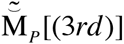 is the contribution from the unique next-fewest-indel history of the 3rd type, Eq.(1.3.11c). According to the definition, Eq.(1.1.2b) supplemented by Eqs.(1.2.4b,6b,8), it is expressed as:

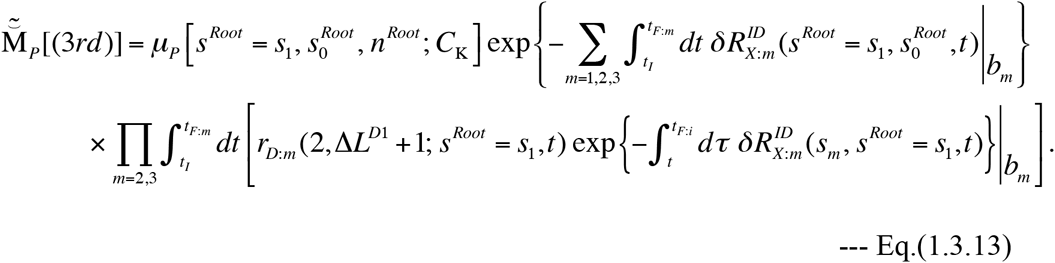

Under Dawg’s model, we have 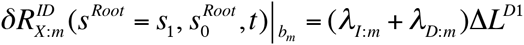 for *m* = 1,2,3, and 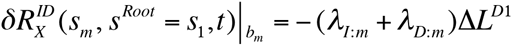. Moreover, if we assume the uniform distribution of the root sequence length, we have 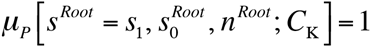 Thus, Eq.(1.3.13) is reduced to:

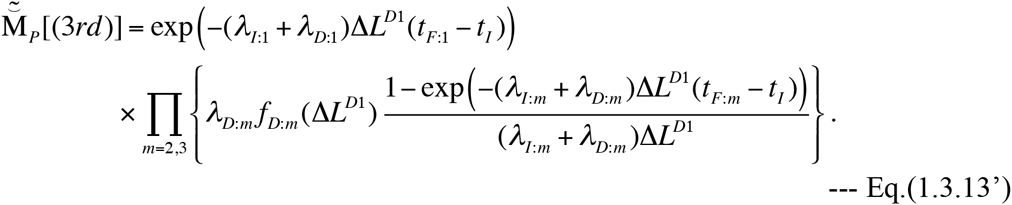

Figure 5 shows the ratio of 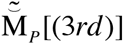 to the fewest-indel history’s contribution, Eq.(1.3.10), when all three branches have the same length and are assigned the same indel model as that used for Figure 3. Because the ratio compares the multiplication factors concerning the indel events along different branches, its value actually depends on several factors. It would be convenient to keep in mind that the ratio could be approximated by 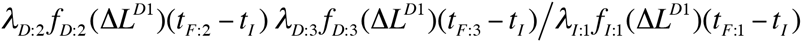 When (*λ*_*I*:*m*_+ *λ D*:*m*)Δ*L*^*D*^(*t* _*F*:*m*_*t*_*I*_)’s are sufficiently smaller than 1 for all *m* = 1, 2, 3. In general, as Δ*L*^*D1*^ gets larger, the ratio is expected to decrease, because the relative frequencies of long indels (*f*_*I*:*m*_ (Δ*L*_*D*1_) and (*f*_*D*:*m*_ (Δ*L*^*D*1^)) are small in general. The ratio is expected to be much smaller than 1 in general. However, it may become quite large when the relative frequency of deletions compared to insertions (*i.e*., the ratio *λ*^*D*:*m*^ *λ*_*I*:*m*_) is considerably larger than 1, or when the lengths of *b*_2_ and *b*_3_ are much larger than that of *b*_1_ (*i.e.*, *t*_*F*:2_ – *t*_*I*_, *t*_*F*:3_ – *t*_*I*_ –s *t*_*F*:1_ – *t*_*I*_). Such situations are similar to those causing the “Felsenstein zone” regarding a substitution model, where a non-parsimonious substitution history at a site is most likely to occur along a tree (see, *e.g*., Chapter 9 of Felsenstein 2004). Under the conditions used to draw Figure 5, an indel history of the 3rd type has a probability much smaller than that of the fewest-indel history. The former is less than 5% of the latter even when as much as 0.4 indels per site are expected to occur. This is probably because the history of the 3rd type requires an exquisite spatial coordination of deletions along different branches. And the result implies that the “Felsenstein zone” of indels should generally be quite narrow, consisting of the cases where a node is connected with branches with extremely unequal lengths.

**Figure 5.**
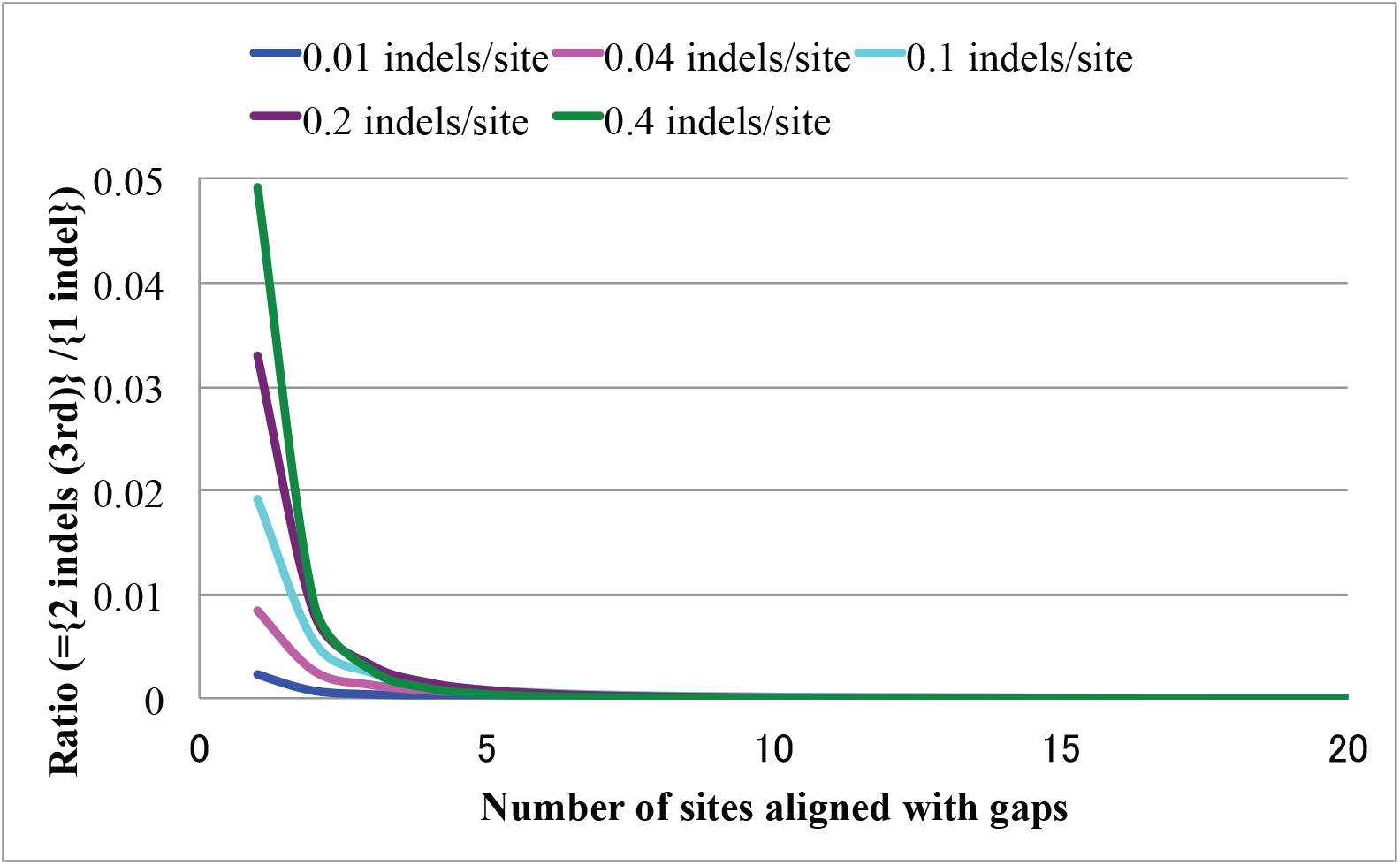
Perturbation analysis on local MSA probabilities in case (III). The graph shows the ratio of the probability of the history with two deletions giving rise to a case (III) local MSA (Eq.(1.3.13’)) to that of the single-insertion history (Eq.(1.3.10)). The ratio is shown as a function of the length of the gapped segment (Δ*L*_*D*1_) and the expected number of indels per site along each branch ((*λ*_*I:m*_ + *λ*_*D:m*_)(*t*_*F:m*_ – *t*_*I*_), identical for all *m* = 1, 2, 3). The blue, magenta, cyan, purple, and green curves show the ratios when the values of (*λ*_*I:m*_ + *λ*_*D:m*_)(*t*_*F:m*_ – *t*_*I*_) are 0.01, 0.04, 0.1, 0.2 and 0.4 indels/site, respectively. We used a 3-OTU tree whose three branches are equally long. For each branch, we used the same parameters as those used for Figure 3.

Case (IV) is represented by the external sequence states, *s*_1_ = [*L*, 1,…, Δ*L*^*DI*^, *R*], *s*_2_ = [*L*, 1,…,, *i*, *j*,…, Δ*L*^*DI*^, *R*], and *s*_3_ = [*L*, *R*], with 1 ≤ *i* + 1 < *j* ≤ Δ*L*^*DI*^ + 1 but (*i*, *j*) ≠(0, Δ*L*^*DI*^ +1) (Here “1,…, 0 ” and “ Δ*L*^*DI*^ +1,…, Δ*L*^*DI*^ ” should be considered to be empty.)In this case, the phylogenetic correctness condition requires the root state to have sites with ancestries 1,…, *i* and *j*,…, Δ*L*^*DI*^, on top of the PASs. Thus, we have 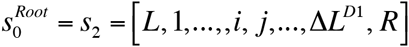. Here, the minimum number of indels is 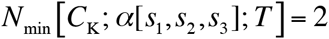. And the set of fewest indel local histories, 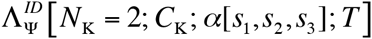 consists of *two* histories. One starts with the root state 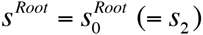, and is represented as:

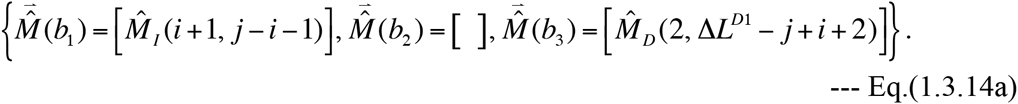

The other starts with the root state *S*^*Root*^ = *s*1 =[*L*, 1,…, Δ*L*^*DI*^, *R*] which differs from 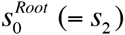 It is represented as:

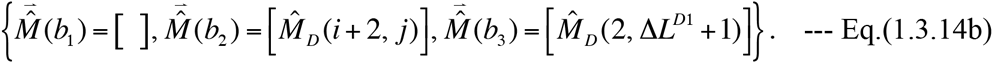

The portion of the multiplication factor contributed from these two fewest-indel local histories, and that contributed from the next-fewest-indel local histories, are calculated in Appendix A2. Table 3 shows the ratio of the total contribution of the next-fewest-indel local histories to that of the fewest-indel local histories for typical configurations in this case. The table indicates that the fewest-indel local histories give a decent approximation of the local MSA probability, as long as the number of descendant sites and the expected number of indels per site are at most moderate.

**Table 3.**
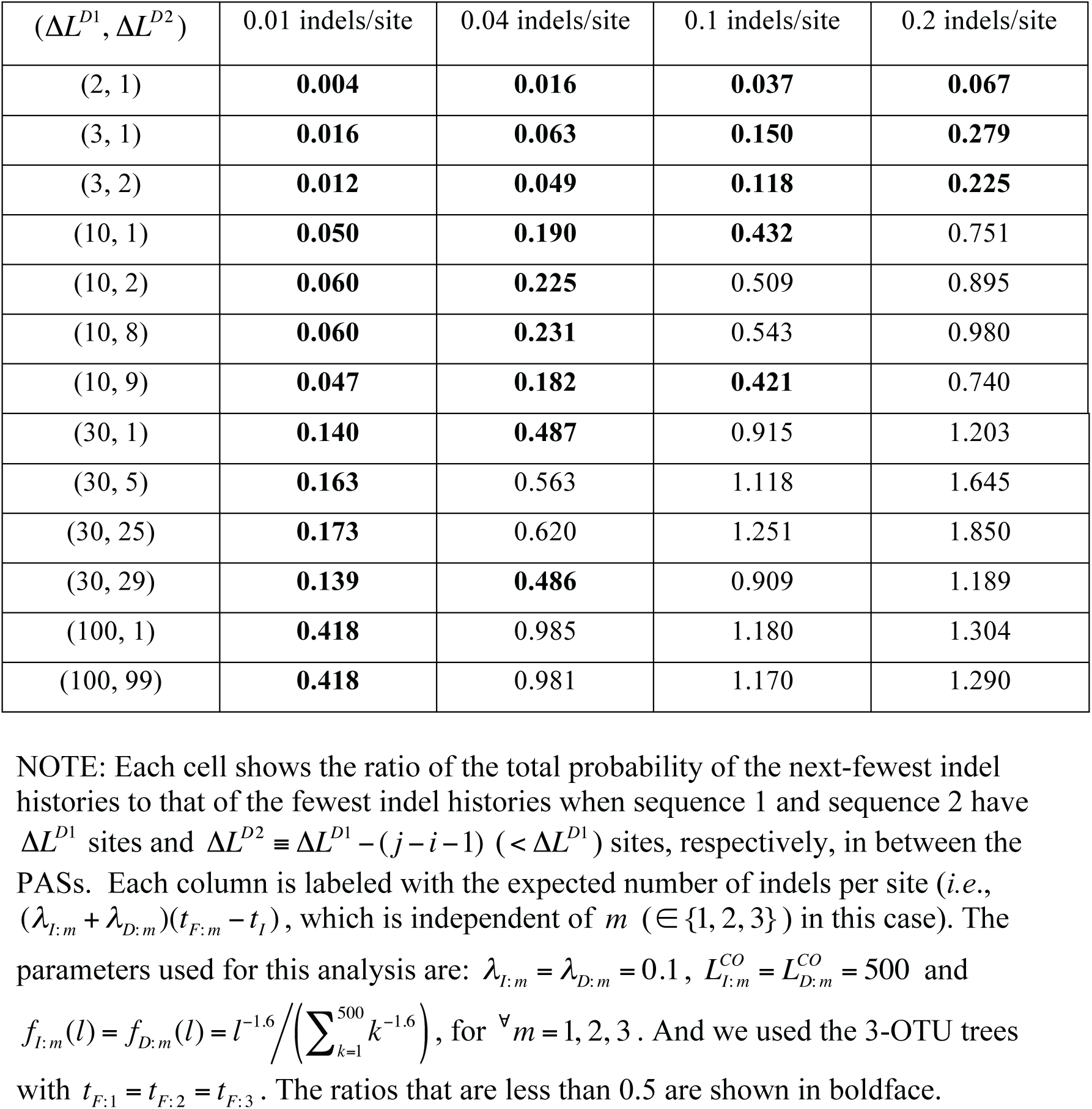
Perturbation analysis on local MSA probabilities in case (IV)

## 2. Goodness of approximation by past indel models

In part I (Ezawa, Graur and Landan 2015a), as well as in Section 1 of this series of paper, we established a theoretical formulation that can calculate the probabilities of given alignments (PWAs or MSAs) under a general continuous-time Markov model, which is a *genuine* evolutionary model describing the stochastic sequence evolution along the time axis via insertions and deletions. The Markov model we dealt with can accommodate quite biologically realistic settings, such as overlapping indels, power-law indel length distributions, and indel rate variation across regions. As shown in Subsection 3.1.1 of part I, the probabilities calculated via our theoretical formulation also fulfill, up to a desired order of the perturbation expansion, the Chapman-Kolmogorov (CK) equation, which is a crucial consistency condition for a *genuine* evolutionary model. To the best of our knowledge, the CK equation has never been satisfied by any previous probabilistic theories allowing for indels of multiple residues. Moreover, the results in Section 5 of part I indicated that even the lowest-order approximation based on the fewest-indel histories alone can approximate the probabilities of the local gap configurations of the alignments considerably well, as long as the indel lengths and the branch lengths are at most moderate. Given these ideal properties, the *ab initio* perturbation theory that we formulated in this series of study can be used as a sound “reference point.” More precisely, other probabilistic models can be compared to it in order to examine how well they can approximate the biologically realistic probabilities of the alignments, and the conditions under which they can. In this section we will examine the “goodness of approximation” by some representative models of indels in the light of our *ab initio* formulation.

### 2.1 Geometric indel length distributions

As mentioned in Introduction, a majority of indel probabilistic models that have been used thus far are based on HMMs or transducer theories that calculates the indel component of the probability of an alignment as a product of column-to-column transition probabilities. Such models can only accommodate geometric indel length distributions, or at most linear combinations of two or more geometric distributions. However, empirical analyses revealed that the lengths of inserted/deleted sequences are distributed according to power-laws (*e.g*., Gonnet et al. 1992; Benner et al. 1993; Gu and Li 1995; Kent et al. 2003; Zhang and Gerstein 2003; Chang and Benner 2004; The international Chimpanzee Chromosome 22 Consortium 2004; Yamane et al. 2006; Fan et al. 2007). Therefore, a question naturally arises as to up to what length a geometric distribution can decently approximate a biologically realistic power-law distribution. Here we address this question. As a reference distribution, we used a power-law distribution:

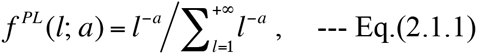

with an empirically typical exponent of *a* = 1.6 (panel A of Figure 6). Then we fitted a scaled geometric distribution:

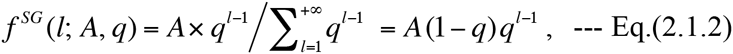

**Figure 6.**
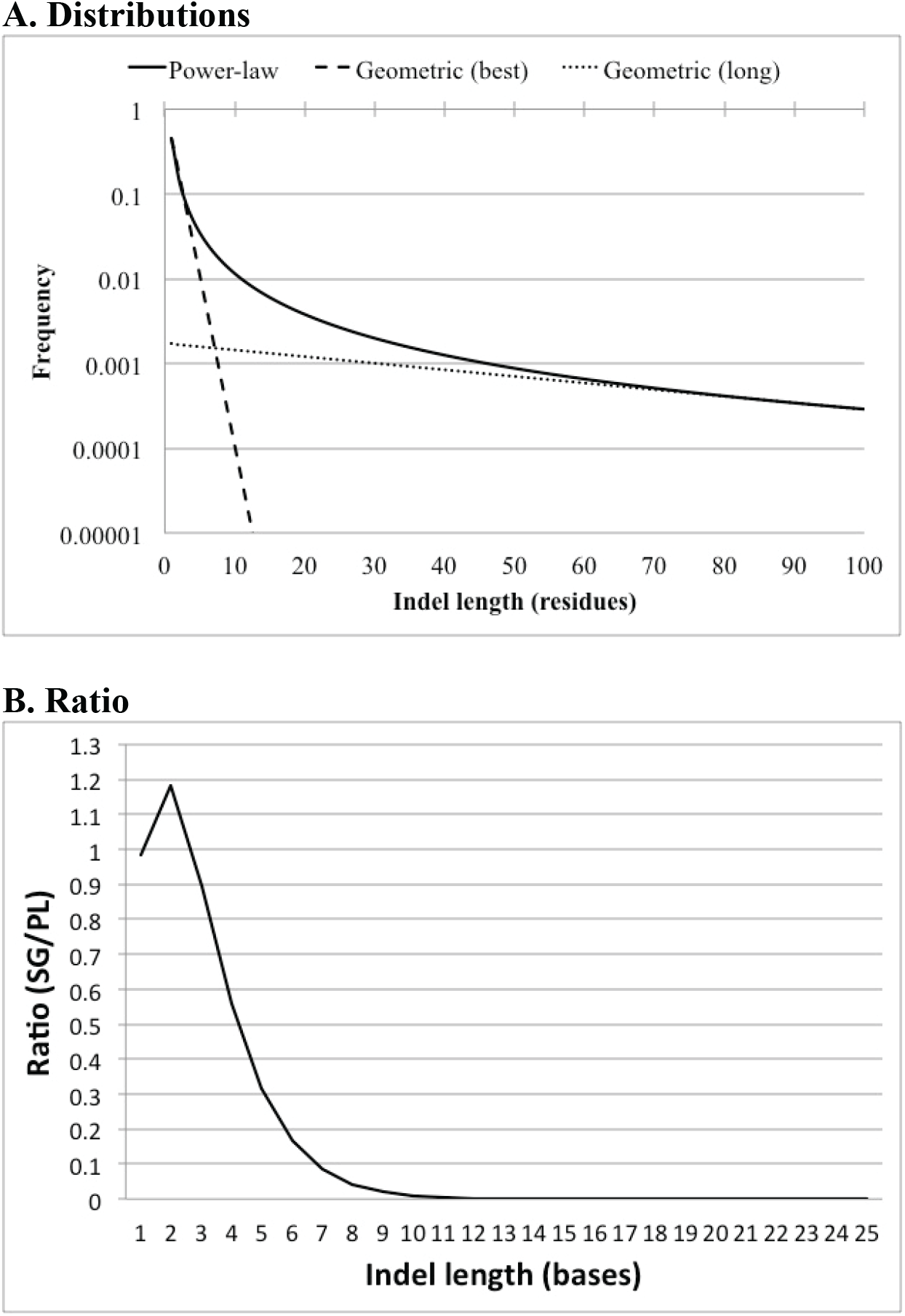
Power-law indel length distribution and its approximation by geometric distributions. **A.** Visual comparison of the distributions. The continuous line is the power-law distribution, Eq.(2.1.1) with *a* = 1.6. The dashed line is the scaled geometric distribution that fits the power-law the best according to the least-square criterion (*i.e*., Eq.(2.1.2) with *A* = *A*_*LS*_ (1.6) ≡ 0.7125 and *q* = *q*_*LS*_ (1.6) ≡ 0.3957). The dotted line is the scaled geometric distribution that well approximates the behavior of the power-law around 100 residues (*i.e.*, Eq.(2.1.2) with *A* = 0.09538 and *q* = 0.9822). Note the logarithmic scale for the ordinate. **B.** The ratio of the best-fit scaled geometric distribution (SG) to the power-law distribution (PL).

to the reference, Eq.(2.1.1), via a least-square fitting across the lengths *l* = 1, 2,…, 100.

We found that the best-fitting parameters are *A* = *A*_*LS*_ (1.6) ≡ 0.7125 and *q* = *q*_*LS*_ (1.6) ≡ 0.3957. (The distribution is drawn in the dashed line in panel A.) Then, we calculated the ratios:

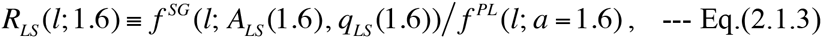

for *l* = 1, 2,…, 25 (panel B of Figure 6). As shown in the figure, *R*_*LS*_ (*l*; 1.6) is less than 1/3 at *l* = 5, less than 1/10 at *l* = 7, and less than 1/1000 at *l* = 13. This indicates that the probabilistic model of indels based on the geometric distribution is somewhat reliable for indels that are at most 4 bases long, which account for 70.5% of all indels if we assume this power-law distribution for all indel lengths. In other words, the geometric distribution substantially underestimates the probabilities of 30% of indels that actually occurred. Although a larger value of *q* could prevent a rapid damping of the estimated frequency, this then causes the mis-estimation of relative frequencies among commonly occurring indels with, *e.g*., *l* = 1, 2, 3 (the dotted line in panel A of Figure 6). Hence, this simple analysis clearly demonstrates how important it is to incorporate biologically realistic indel length distributions into the evolutionary model.

### 2.2 Models pursuing biological realism

Some existing probabilistic models of indels, such as the “long indel” model (Miklós et al. 2004) and the model of Kim and Sinha (2007), aim to pursue more biological realism by accommodating indel length distributions that are biologically more realistic than geometric distributions. As noted in Subsection 5.1 of part I (and proved in Appendix A6 of part I), as far as each LHS equivalence class is concerned, the probability given by the recipe of Miklós et al. (2004) equals the probability calculated by our formulation under the space-and time-homogeneous model parameters (given in Eqs.(2.4.5a,b) of part I). Thus, regarding the probability of a PWA, the two methods should give the identical sum of contributions from the fewest-indel histories. And, *provided that* both methods can correctly enumerate all possible local histories of any given number of indels consistent with each local gap configuration, they will also give the identical sum of contributions from the histories with up to a desired number of indels. Fulfilling this last condition, however, will be an important task left for future studies. (At present, we cannot tell whether or not the algorithm of Miklós et al. (2004) can correctly enumerate indel histories, because their paper has no explicit description on this topic. So, we will not discuss their method further in this section.)

Here, we specifically examine the model of Kim and Sinha (2007) in the light of our theoretical formulation. Their model is a block-wise HMM, and calculates the probability of a PWA between the ancestral and descendant sequences along a branch as a product of block-wise probabilities. In their HMM, a block is either a column of a PAS, a run of gaps in the ancestor aligned with a run of residues in the descendant, or a run of gaps in the descendant aligned with a run of residues in the ancestor. Each PWA is actually a part of a MSA of given sequences at the external nodes and one of alternative sets of sequences at internal nodes. Ancestral gaps aligned with descendant gaps are removed before evaluating the probability of a PWA. Because their purpose is to find a most likely indel history and a resulting set of consistent ancestral sequence states at internal nodes, they are not interested in an indel event that begins and/or ends in the middle of a block. Thus, they only consider those events that insert/delete the entire blocks in single steps.

We now calculate the probabilities of the local PWAs that were considered in the cases (i)-(iv) in Subsection 1.2, via the model of Kim and Sinha (2007). And we compare the results to those via our theoretical formulation under Dawg’s parameters (Eqs.(2.4.4a,b) of part I). In the following, the probabilities via Kim and Sinha (2007) will be calculated according to Eq.(2) and Figure 1C of their paper, and the probabilities via our formulation will be calculated according to the prescriptions in Subsection 1.2 (and in Appendix A1). We set *t*_*F*_ – *t*_*I*_ = | *b* | in the following calculations. (Here | *b* | denotes the length of branch *b*). Via the model of Kim and Sinha (2007), the PWA probability in case (i) is calculated as:

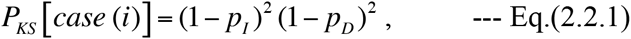

Where *p*_*I*_ and *p*_*D*_ are the “transition probabilities of the ‘Insertion’ and ‘Begin deletion’ states,” respectively. Via our formulation, the PWA probability is:

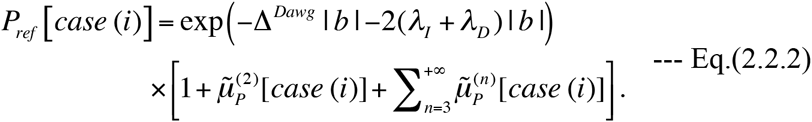

Here, Δ^*Dawg*^ is the abbreviation of the “universal” factor for the indel exit rate (*i.e*., Δ^*Dawg*^[*λ*_*I*_*, λ*_*D*_, *f*_*D*_ (.)] in Eq.(2.4.4c) of part I). And 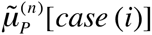 is short for 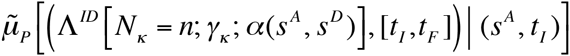, *i.e.,* the summed contribution from the *n* -event local histories to the multiplication factor in case (i). Especially, 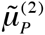 [*case*(*i*)] is concretely expressed in Eq.(1.2.2a). Now, assuming that (*λ*_*i*_+*λ* _*D*_)| *b* | is sufficiently small, we expand the expression in the square brackets into the power series in *λ*_*I*_ | *b* | and *λ*_*D*_ | *b* |, which will collectively be denoted as *λ* | *b* | when considering the order of magnitude. From Eqs.(1.2.2a,b’), we get 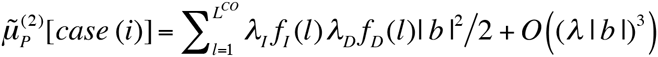 Moreover, the expansion of 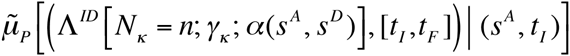 generally starts with *O* (*λ* | *b* |)^*n*^ terms. Thus, we have:

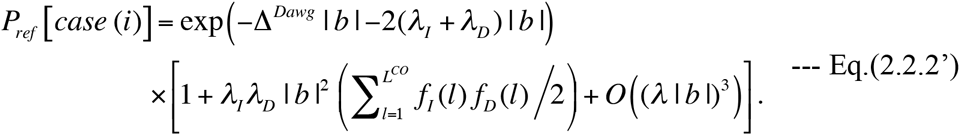

This and Eq.(2.2.1) will provide the baseline when examining the probabilities in Kim and Sinha’s model in other cases.

In case (ii), the PWA probability under the HMM of Kim and Sinha (2007) is calculated as:

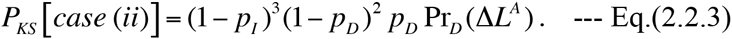

Here Pr_*D*_ (*l*) is the “probability distribution on the deletion length (*l*),” which is assumed as shared among different branches. To facilitate the comparison, we consider the ratio of the probability in case (ii) to that in case (i), which yields:

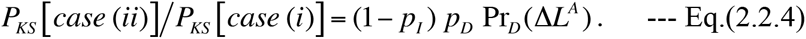

Meanwhile, the probability via our formulation is:

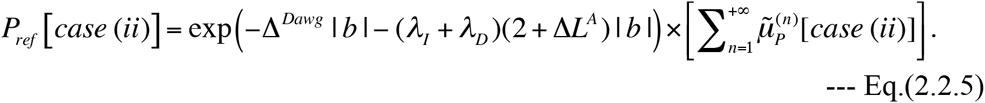

Here, 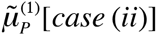 is given by Eq.(1.2.4’), and 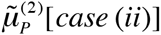 is given by Eq.(1.2.5a) supplemented with Eqs.(1.2.5b,c,d’e’). The ratio of Eq.(2.2.5) to Eq.(2.2.2) is:

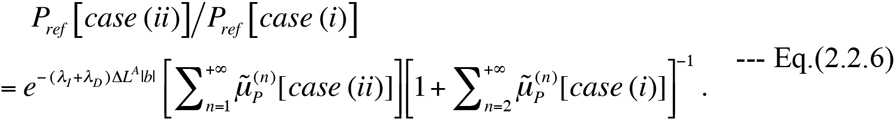

Expanding this expression into the power series in *λ* | *b* |, we have:

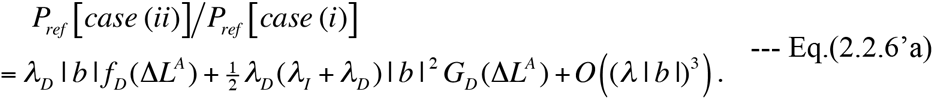

Here *G* (Δ*L*^*D*^) is defined as:

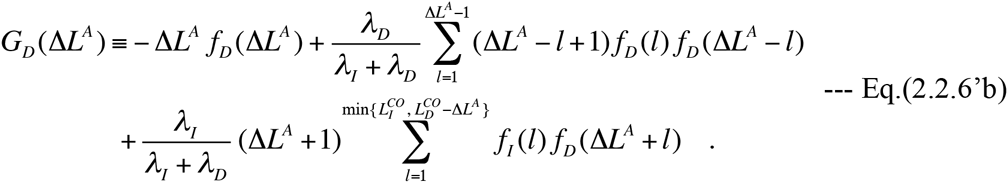

Figure 7 shows the ratio *G*_*D*_(Δ*L*^*A*^)/*f*_*D*_(Δ*L*^*A*^) as a function of Δ*L*^*A*^.

Similarly, according the HMM of Kim and Sinha (2007), the ratio of the PWA probability in case (iii) to that in case (i) is expressed as:

**Figure 7.**
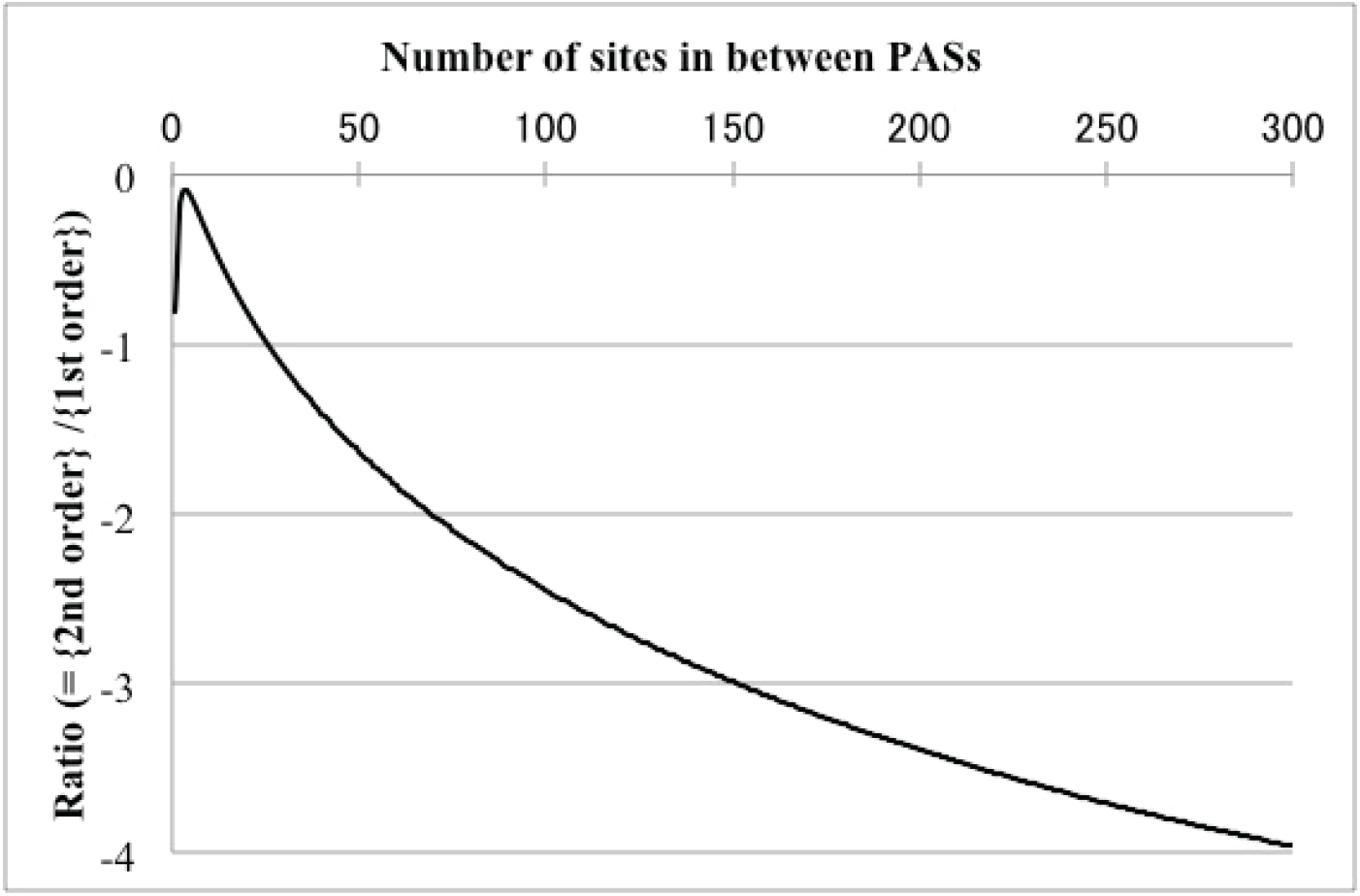
Coefficient of 2nd-order term of PWA probability in cases (ii) and (iii). The curve shows the ratio *G*_*D*_(Δ*L*^*A*^)/*f*_*D*_(Δ*L*^*A*^) as a function of the number of sites in between the PASs (Δ*L*^*A*^) in a local PWA belonging to case (ii). Here *G*_*D*_(Δ*L*^*A*^) is the coefficient of the 2nd-order term in the expected number of indels (*λ* |*b*|) given in Eq.(2.2.6’b) in Results. And *f*_*D*_(Δ*L*^*A*^) is the coefficient of the 1st-order term (see Eq.(2.2.6’a)). The ratio was calculated with the following parameters: *λ*_*I*_ = *λ*_*D*_ = 0.1, 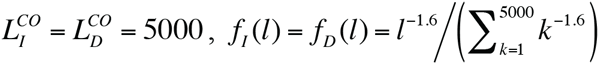. Under the same parameter setting, this graph also gives the ratio for a case (iii) local PWA as a function of Δ*L*^*D*^ (*G*_1_(Δ*L*^*D*^)/*f*_*I*_(Δ*L*^*D*^). See Eqs.(2.2.8a,b)). The value, 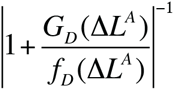, gives a rough estimate of the “threshold” value of (*λ*_*I*_ + *λ*_*D*_)|*b*| beyond which the violation of the Chapman-Kolmogorov equation (Eq.(3.1.1.1) in part I) may severely undermine the HMM of Kim and Sinha (2007).

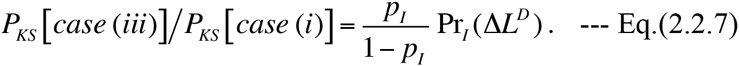

Here Pr_*I*_ (*l*) is the “probability distribution on the insertion length (*l*),” which also is assumed as shared among different branches. The ratio via our formulation is obtained by the power-series expansion in *λ* |*b*| of Eq.(A1.1.1’) and Eq.(A1.1.2a) supplemented with Eqs.(A1.1.2b,c,d’,e’) (all in Appendix). The result is:

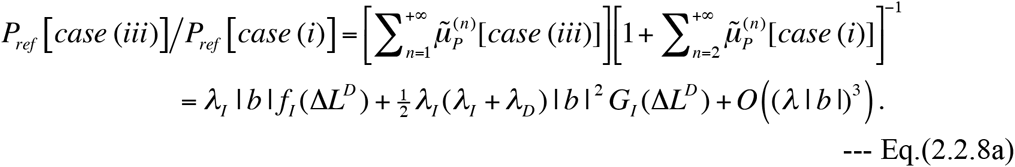

Here *G* (Δ*L*^*DI*^) is defined as:

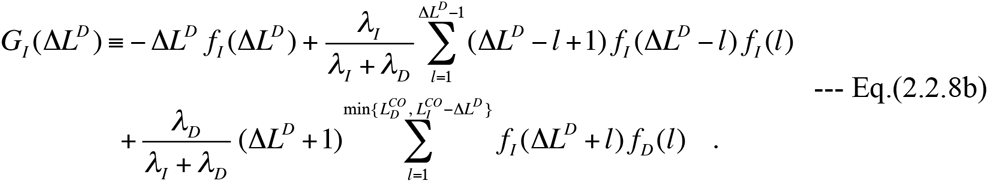

Thanks to the symmetry of the probabilities under the time reversal, Figure 7 also gives the ratio *G*_*I*_(Δ*L*^*D*^)/*f*_*I*_(Δ*L*^*D*^) as a function of Δ*L*^*D*^, when calculated under same setting.

Now we compare the results, Eq.(2.2.4) and Eq.(2.2.7), under Kim and Sinha’s model with the corresponding results, Eq.(2.2.6’a) and Eq.(2.2.8a), obtained via our formulation. In the method of Kim and Sinha (2007), the substitutions *p*_*I*_ = *c*_*I*_ |*b*| and *p*_*D*_ = *c*_*D*_ |*b*| are made first. Then *c*_*I*_ and *c*_*D*_ are estimated from the total frequencies of insertions and deletions, respectively, along the external branches observed from the input MSA. Similarly, Pr_*I*_(Δ*L*^*A*^) and Pr_*D*_(Δ*L*^*A*^) are estimated from the observed length histograms for insertions and deletions, respectively. Let {*b*}_*PE*_ be the set of branches used for parameter estimations. Then, using the summations of the “reference” results, Eq.(2.2.6’a) and Eq.(2.2.8a), both over {*b*}_*PE*_, we expect to have:

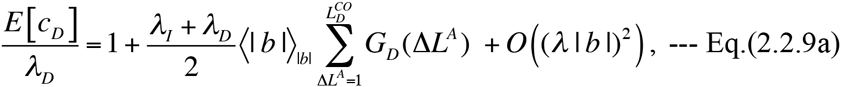

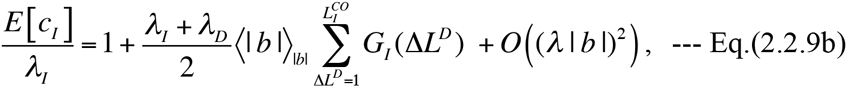

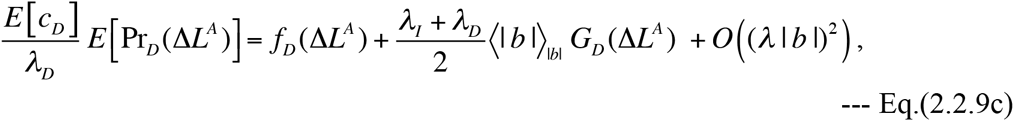

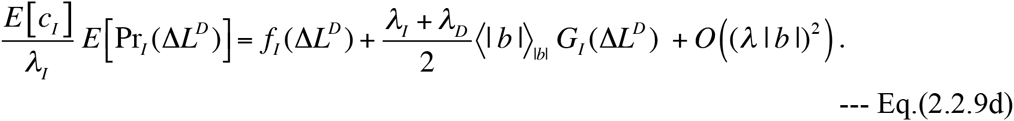

Here *E* [*X*] denotes the expected value of the estimated parameter *X*, which is the average of estimated *X* over all indel processes under the given set of conditions (the tree and model parameters). And we also used the notation, 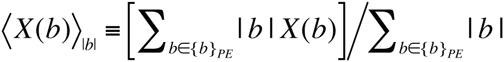 [NOTE: The actual values of *c*_*I*_ and *c*_*D*_ estimated by the method of Kim and Sinha (2007) may be slightly smaller than Eqs.(2.2.9b,c), because the denominator in their method is the total number of MSA columns, instead of the average numbers of possible indel positions in ancestral sequences.] Usually, 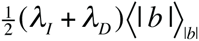 is quite small, at most *O* (10^−1^) and typically *O* (10^−2^). Thus, as long as the actual parameters, *λ*_*I*_, *λ*_*D*_, *f*_*I*_ (Δ*L*^*D*^), and *f*_*D*_ (Δ*L*^*A*^), do not considerably vary across branches, and provided that the MSA is sufficiently long and accurate, the estimated values of *c*_*I*_ and *c*_*D*_, respectively, should approximate *λ*_*I*_ and *λ*_*D*_ fairly well. Also, under the same situation, the estimated values of Pr_*I*_(*l*) and Pr_*D*_(*l*), respectively, should approximate *f*_*I*_(*l*) and *f*_*D*_ (*l*) fairly well, as long as the ratios *G*_*I*_(*l*)/*f*_*I*_(*l*) and *G*_*D*_(*l*)/*f*_*D*_ (*l*) are sufficiently less than 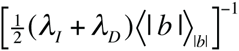. However, it does not actually matter so much whether or not the estimated Kim-Sinha parameters (*c*_*I*_, *c*_*D*_, Pr_*I*_(*l*) and Pr_*D*_(*l*)) approximate Dawg’s indel parameters (*λ*_*I*_, *λ*_*D*_, *f*_*I*_(*l*) and *f*_*D*_ (*l*)) fairly well. What actually matters is how accurately Eq.(2.2.4) and Eq.(2.2.7) approximate Eq.(2.2.6’a) and Eq.(2.2.8a), respectively, using the estimated parameters. In an extreme case where all branches have the same branch length, the approximation should be nearly perfect. This is because, in this case, |*b*| ≈ ⟨| *b* |⟩ _|*b*|_ for all branches, and thus because we can use Eqs.(2.2.9a-d) without any significant modifications to estimate the probabilities (not involving case (iv)). [It should be noted here that Eq.(2.2.4) and Eq.(2.2.7), respectively, contain extra multiplication factors, (1− *p*_*I*_) and (1− *p*_*I*_)^−1^, compared to the corresponding Eq.(2.2.6’a) and Eq.(2.2.8a). However, these factors should remain close to 1, because *p*_*I*_ = *c*_*I*_ | *b* | should normally be at most *O* (10^−1^).] In contrast, the approximations by Eq.(2.2.4) and Eq.(2.2.7) could considerably deteriorate, *e.g*., when (|*λ*_*I*_ + *λ*_*D*_)(| *b* | − 〈| *b* |〉 _|*b*|_|) times the ratios, *G*_*D*_ (Δ*L*^*A*^)/*f*_*D*_ (Δ*L*^*A*^) and *G*_*I*_ (Δ*L*^*D*^)/ *f*_*I*_ (Δ*L*^*D*^), respectively, become comparable to or greater than 1 (unity). Because *G*_*D*_ (Δ*L*^*A*^) and *G*_*I*_ (Δ*L*^*D*^) are mostly contributed from next-fewest-indel local histories containing overlapping indels, we can interpret the result as follows. “Overlapping indels start to make Kim and Sinha’s method poorly approximate the alignment probabilities when the involved gap is long and the branch lengths show a large variation.” If there is a good reason to believe that the Dawg indel parameters (*λ*_*I*_, *λ*_*D*_, *f*_*I*_ (*l*) and *f*_*D*_ (*l*)) are shared among all branches, one way to mitigate the aforementioned effects of overlapping indels may be to set:

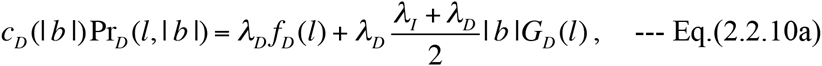

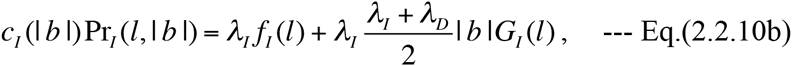

and to fit *λ*_*I*_, *λ*_*D*_, *f*_*I*_(*l*) and *f*_*D*_(*l*) according to these equations supplemented with Eqs.(2.2.6’b,8b). Now, as indicated by Figure 7, under the power-law indel length distributions, the ratios *G*_*D*_(Δ*L*^*A*^)/*f*_*D*_(Δ*L*^*A*^) and *G*_*I*_(Δ*L*^*D*^)/*f*_*I*_(Δ*L*^*D*^) are less than 4 in absolute value when the gap is 300 residues long or shorter. Therefore, the 2nd-order factors should remain close substantially influence the results when 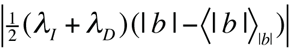 is larger than, say, 0.1. Such a situation will be quite rare in practical sequence analyses. Even if we encounter such a rare case, then local histories with more than 2 indels will begin to account for a substantial fraction of the probability. Considering this way, we expect that the method of Kim and Sinha (2007) will pretty well approximate the probabilities of local PWAs belonging to cases (ii) and (iii) at least within the threshold 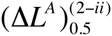 defined in Subsection 1.2.

Finally, we consider case (iv). Indel histories giving rise to the local sequence states in this category are shown, *e.g*., in Figure 5, panels F and G of Figure 3, and panel A of Figure 6 (all of them are in part I). In such a situation, an aligner will reconstruct a MSA that is like either panel B or C of Figure 5 of part I (if the reconstruction is correct), and Kim-Sinha’s method assigns a probability according to the reconstructed MSA. Whether it is like panel B or panel C, the assigned probability is the same, and its ratio to the probability of case (i) is:

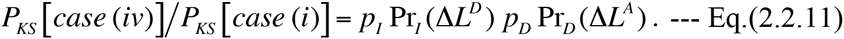

Via our formulation, how to calculate the probability in this case was briefly described in the middle of Subsection 1.2, and detailed in Appendix A1.2. In this case, each fewest-indel history consists of two indels, and each next-fewest-indel history consists of three indels. Because there are as many as 24 types of next-fewest-indel histories, here we only consider the fewest-indel histories. Then, the lowest-order contribution of the multiplication factor, 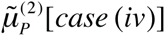, is given by Eq.(A1.2.1a), supplemented with Eqs.(A1.2.1b,c,d,e’,f’,g’). Expanding each term into a power series in *λ* | *b* |, we get the following expression for the ratio:

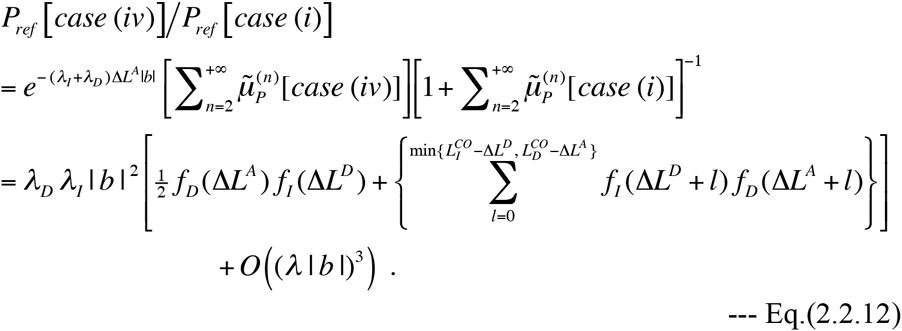

In case (iv), as opposed to in cases (ii) and (iii), the PWA probabilities via the HMM of Kim and Sinha (2007) differ considerably from that via our formulation *even when* ⟨| *b* |⟩ _|*b*|_ ≪ 1 and | *b* | ≪ 1. Under these conditions, *p*_*I*_ Pr_*I*_ (Δ*L*^*D*^) and *p*_*D*_ Pr_*D*_ (Δ*L*^*A*^) quite accurately approximate *λ*_*I*_ | *b* | *f*_*I*_ (Δ*L*^*D*^) and *λ*_*D*_ | *b* | *f*_*D*_ (Δ*L*^*A*^), respectively (see Eqs.(1.2.9c,d)), and the *O* ((*λ* | *b* |)^3^) terms in Eq.(2.2.12) can be neglected. Thus, we have:

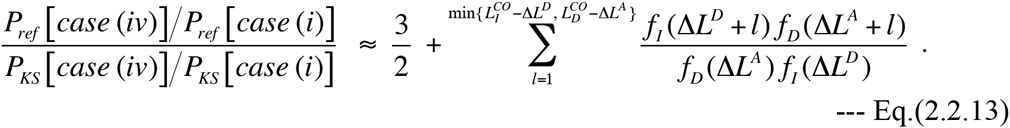

Table 4 shows this ratio for typical cases. The second term on the right hand side of Eq.(2.2.13) is the effect of overlapping indels. When Δ*L*^*A*^ = Δ*L*^*D*^ = 1, this term is expected to be quite small; for example, it is about 0.167 if *f*_*I*_(*l*) = *f*_*D*_(*l*) ∞ *l*^−1.6^. And it gets more and more influential when Δ*L*^*A*^ and/or Δ*L*^*D*^ gets larger, and it substantially exceeds 1 (unity) in some cases (Table 4). Actually, a similar effect was incorporated in the HMM of Knudsen and Miyamoto (2003). Their HMM could only accommodate geometric indel length distributions, and consequently the relevant term was independent of Δ*L*^*A*^ and Δ*L*^*D*^. Coming back to Eq.(2.2.13), the first term on the right hand side, 3/2, differs from 1 (unity) because the HMM of Kim and Sinha (2007) does not correctly take account of the non-overlapping indel histories, either. This error is actually shared by most of the standard, or nearly standard, HMMs and transducers used thus far as probabilistic models of indels (see Background of part I). Taking these results into consideration, another possible improvement on the model of Kim and Sinha (2007) would be to modify the HMM structure so that the probability of an insertion and an immediately adjacent deletion (or that of the opposite configuration) will be given by Eq.(2.2.12), or by its extension to include the terms of higher-orders in *λ* | *b* |.

**Table 4.** Comparison of Kim-Sinha’s probabilities of case (iv) local PWAs with reference probabilities.

| $(\Delta L^A, \Delta L^D)$ | Ratio (Eq.(VII-2.13)) | Overlapping <sup>a</sup> |
| --- | --- | --- |
| (1, 1) | 1.667 | 0.167 |
| (3, 1) | 1.883 | 0.383 |
| (3, 3) | 2.449 | 0.949 |
| (5, 5) | 3.325 | 1.825 |
| (10, 1) | 2.165 | 0.665 |
| (10, 10) | 5.572 | 4.072 |
| (25, 1) | 2.355 | 0.855 |
| (25, 4) | 4.714 | 3.214 |
| (30, 10) | 8.300 | 6.800 |
| (100, 1) | 2.561 | 1.061 |
| (100, 3) | 4.896 | 3.396 |
| (300, 1) | 2.659 | 1.159 |

NOTE: Each cell in the middle column gives the ratio, Eq.(2.2.13), of the reference probability in the fewest-indel approximation to the corresponding probability given by Kim and Sinha's HMM (2007), for a local PWA with $\Delta L^A$ ancestral sites and $\Delta L^D$ descendant sites in between a pair of PASSs. The parameters used for this analysis are: $\lambda_I = \lambda_D = 0.1$ , $L_I^{CO} = L_D^{CO} = 5000$ , and $f_I(l) = f_D(l) = l^{-1.6} / \left( \sum_{k=1}^{5000} k^{-1.6} \right)$ . The ratio for $(\Delta L^A, \Delta L^D) = (L_1, L_2)$ is identical to that for $(\Delta L^A, \Delta L^D) = (L_2, L_1)$ . Thus, we only showed the results for $\Delta L^A \geq \Delta L^D$ .
<sup>a</sup> The effect of overlapping indels, which is the second term on the right hand side of Eq.(2.2.13).

### 2.3 Violation of Chapman-Kolmogorov equation

The Chapman-Kolmogorov (CK) equation (Eq.(3.1.1.1) in part I),

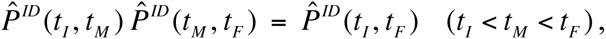

is a crucial condition that must be satisfied by *genuine* evolutionary models based on any continuous-time Markov models. Thus far, the CK equation was violated by most HMMs and transducer theories, except those directly derived from evolutionary models, *e.g*., the TKF91 model (Thorne et al. 1991). As formally shown in Appendix A3 of part I (Ezawa, Graur and Landan 2015a), our theoretical formulation could satisfy the CK equation up to any desired order of the perturbation expansion. Therefore, our formulation enables us to analytically examine the effects of the violation of the CK equation. Here, we will only consider a simplest yet non-trivial example, *i.e*., case (ii) in Subsection 1.2. And we will examine the HMM of Kim and Sinha (2007) because of its generality; most HMMs could be obtained from their HMM by slightly modifying the parameters and by fixing the indel length distributions to geometric ones, possibly with time-dependent gap-extension probabilities.

Via the HMM of Kim and Sinha (2007), the PWA probability in case (ii) was given by Eq.(VII-2.3), with the substitutions *p*_*I*_ = *c*_*I*_ | *b* | and *p*_*D*_ = *c*_*D*_ | *b* |:

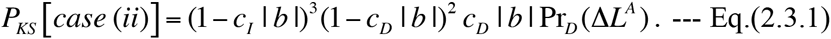

In contrast, the probability in the same case via our formulation was given by Eq.(2.2.5), which can be partially expanded into the power-series in *λ* | *b* | as:

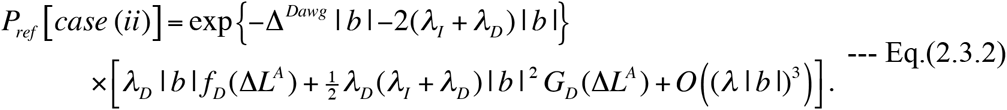

Here *G*^*D*^ (Δ*L*^*A*^) was given in Eq.(2.2.6’b). And exp{-Δ^*Dawg*^ | *b* |} is a “universal factor” that occur in the probability of any indel processes on the entire sequence (along branch *b*). Eq.(2.3.2) satisfies the CK equation up to (and including) *O* ((*λ* | *b* |)^2^). Now, compare Eq.(2.3.1) with Eq.(2.3.2), aside from the universal factor. First, if we set *c*^*I*^ = *λ*^*I*^ and *c*^*D*^ = *λ*^*D*^, (1− *c*^*I*^ | *b* |)^2^ (1 − *c*^*D*^ | *b* |)^2^ should approximate exp{−2(*λ*_*I*_ + *λ*_*D*_)| *b* |} quite well, because *λ*^*I*^ | *b* | and *λ*_*D*_ | *b* | should usually be at most *O*(10^−1^). Thus, it is sufficient to compare (1− *c*^*I*^| *b* |)*c*_*D*_ | *b* | Pr_*D*_ (Δ*L*^*A*^) with the expression in the square brackets on the right hand side of Eq.(2.3.2). If Pr_*D*_ (Δ*L*^*A*^) is considered as independent of | *b* |, it should be equal to *f*^*D*^ (Δ*L*^*A*^). Then, whether or not Eq.(2.3.1) approximately satisfies the CK equation depends on whether or not the absolute value of the difference, 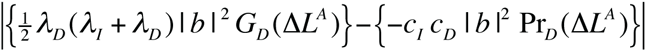 is negligible compared to *λ*^*D*^ | *b* | *f*^*D*^ (Δ*L*^*A*^). As Figure 7 indicates, this condition will hold if Δ*L*^*A*^ is within a “critical value,” which decreases as (*λ*^*I*^ + *λ*^*D*^)| *b* | increases. Once Δ*L*^*A*^ exceeds the critical value, the violation of the CK equation will considerably impact on the probability estimation, and therefore on the data analyses. Under the parameter setting used for Figure 7, the critical value is 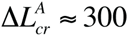 300 when (*λ*^*I*^+λ_*D*_)| *b* | ≈ 0.6. This means that, *as far as case (ii) local gap-configurations are concerned*, Kim and Sinha’s HMM practically satisfies the CK equation at least up to *O* ((*λ* | *b* |)^2^).

However, we expect that their HMM will severely violate the CK equation when it is applied to case (iii) local PWAs, in view of the results in Subsection 2.2.

Some past simulation analyses (*e.g*., Thorne et al. 1992; Knudsen and Miyamoto 2003; Metzler 2003) seem to have concluded that the violation of the CK equation, or its cause, *i.e*., the failure to accommodate overlapping indels, did not seriously impact on the results of data analyses. These results might be because they used geometric distributions of indel lengths. To see if this is indeed the case, let’s assume *f*^*I*^(*l*) = *f*^*D*^(*l*) = (1− *q*)*q*^*l*−1^. Substituting them into Eq.(2.2.6’b), we have:

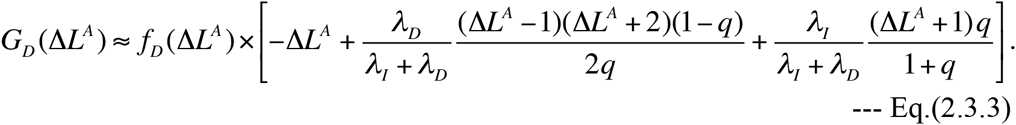

Here we used 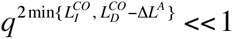 to obtain the approximation. Now, with the approximations *c*_*I*_≈ *λ*_*I*_, *c*_*D*_ ≈ *λ*_*D*_ and Pr_*D*_ (Δ*L*^*A*^) ≈ *f*_D_ (Δ*L*^*A*^), we calculate the ratio:

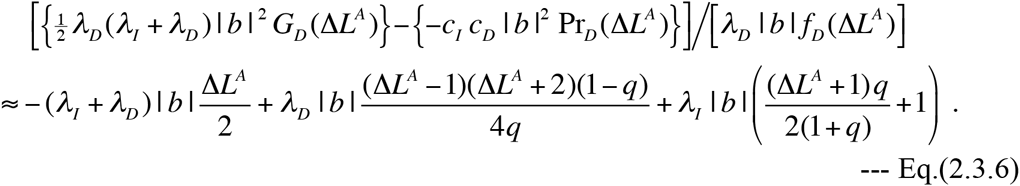

If Δ*L*^*A*^ is quite large, *e.g*., 10 or greater, the middle term on the right hand side will predominate, and the ratio could be roughly approximated as

*λ*_*D*_ |*b*| (Δ*L*^*A*^)^2^ (1− *q*) (4*q*). Thus, the “critical value” is roughly given by 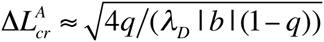. If we set, *e.g*., *λ*_D_= 0.1 and *q* = *q*_*LS*_(1.6) = 0.3957 as given above Eq.(2.1.3), the critical value would be: 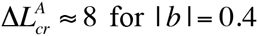, and 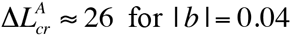. At these critical values, geometric deletion length frequencies are already pretty small; we have *f*(Δ*L*^4^-8)≈9.2×10^−4^and *f*_*D*_ (Δ*L*^*A*^ = 26) ≈ 5.2 ×10^−11^ when *q* = *q*_*LS*_ (1.6) = 0.3957. Therefore, as long as the geometric indel length distributions are used for simulations, it would be very rare to encounter indels whose lengths exceed the critical value. Moreover, as shown in Subsection 2.1, when the indels are longer than these critical values, models with geometric indel length distributions severely underestimate their frequencies, to the extent that they are deemed almost useless. With these circumstances compounded together, it would come as no surprise that the past studies did not detect significant impacts of the violation of the CK equation on their results. [NOTE: Precisely speaking, (nearly) standard HMMs and transducers can take account of *a part* of contributions from multiple deletions. For example, the total contribution of two-deletion histories that they can capture is (Δ*L*^*A*^ −1)(*λ_D_* | *b* |)^2^ (1− *q*) *f*_*D*_ (Δ*L*^*A*^). Here (Δ*L*^*A*^ −1) comes from the number of ways in which a run of gaps can be split into two segments. Thus, its ratio to the first-order term, *λ_D_* | *b* | *f*_*D*_ (Δ*L*^*A*^), is (Δ*L*^*A*^ −1)(*λ_D_* | *b* |)(1− *q*). This is roughly 4*q* Δ*L*^*A*^ times as large as the dominant term in Eq.(2.3.6). This ratio is much smaller than 1 (unity) when Δ*L*^*A*^ is around or greater than the above critical values. Therefore, such histories with two contiguous deletions have only negligible effects on the present argument.]

In contrast, if we incorporate biologically realistic indel length distributions into indel probabilistic models that are *not genuine* evolutionary models, such effects may become remarkable when dealing with long indels and/or long branches. The effects, however, might be somewhat alleviated by fitting indel rate parameters and length distributions that depend on the branch length as in Eqs.(2.2.10a,b).

## Conclusions

In a previous paper (Ezawa, Graur and Landan 2015a), we approached the fundamental problems on the probability of indels in an absolutely orthodox manner. More specifically, we established the theoretical basis of an *ab initio* perturbative formulation of a general continuous-time Markov model, which is a *genuine* stochastic model describing the evolution of an *entire* sequence via indels along the time axis. Using the formulation, we proved that, if the indel model parameters satisfy a certain set of conditions, the probability of an alignment is indeed factorable into the product of an overall factor and contributions from local alignments delimited by preserved ancestral sites (PASs).

In this paper, we concretely calculated the probability contributions from individual local alignments using perturbation analyses. The results indicated that even the fewest-indel terms alone can approximate the probabilities pretty well as long as the branch lengths and the indel lengths are at most moderate. And we also clarified the parameter regions where the alignment probabilities are safely approximated by the hidden Markov models (HMMs) of indels (and by association the transducer theories) that were often used in the past. We showed that the approximation by the geometric indel length distributions grossly deteriorates as soon as the indels become longer than 4 sites. We also showed that these HMMs violate the Chapman-Kolmogorov (CK) equation because they lack most of the non-fewest-indel terms in the perturbation expansion. This finding enables to predict the regions where the violation of the CK equation starts to compromise the reliability of these models. The analyses also suggested possible modifications to these models in order to improve their accuracy.

To summarize, by depending purely on the first principle, our *ab initio* perturbation formulation provides a sound reference point to which other indel models can be compared in order to see when and how well they can approximate the true alignment probabilities.

## Methods

### Implementation of the formulas

Most of the formulas used for the perturbation analyses were implemented in Perl. They are available as a part of the package named LOLIPOG (log-likelihood for the pattern of gaps in MSA), which in turn is available at the FTP repository of the Bioinformatics Organization (Ezawa 2013).

## Authors’ contributions

KE conceived of and mathematically formulated the theoretical framework in this paper, implemented the key algorithms, participated in designing the study, performed all the mathematical analyses, and drafted the manuscript. DG and GL participated in designing the study, helped with the interpretation of the data, and helped with the drafting of the manuscript.

## Acknowledgements

This study is dedicated to the late Dr. Keiji Kikkawa, who was a renowned theoretical physicist, one of the key pioneers of the string field theory of the elementary particle physics, and the best ever mentor of K.E. We are grateful to Dr. R. A. Cartwright at Arizona State University for his useful information and discussions that inspired this study. We appreciate the logistic support and the feedback of Dr. Tetsushi Yada at the Kyushu Institute of Technology. We would also like to thank the three anonymous referees of the predecessor manuscript entitled: “Framework that enables approximate lilelihood analysis of insertions/deletions on multiple sequence alignment.” Their comments helped drastically improve the study itself, not to mention the manuscript. This work was a part of the project, “Error Correction in Multiple Sequence Alignments,” which was funded by US National Library of Medicine [grant number LM010009-01 to Dan Graur and Giddy Landan at the University of Houston]. The later stage of this work was also supported by Grants-in-Aid No. 221S0002, which was awarded to Tetsushi Yada by the Ministry of Education, Culture, Sports, Science and Technology of Japan.

## List of abbreviations

CK: Chapman-Kolmogorov
CHMM: hidden Markov model
indel: insertion/deletion
LHS: local history set
MSA: multiple sequence alignment
PAS: preserved ancestral site
PWA: pairwise alignment

## Appendix

### A1. Perturbation calculation of multiplication factors for PWAs between ancestral and descendant sequences

#### A1.1. For case (iii) local PWAs

Here we detail the calculation of the sum of contributions from the fewest-indel events as well as that from the next-fewest-indel events in case (iii) considered in Section 1.2 of Results, that is, the case where the ancestor has no site but the descendant has one or more sites in between a pair of preserved ancestral sites (PASs) (Figure 2 C).

In case (iii), we assume that the descendant state has Δ*L*^*D*^ sites in between the flanking PASs. Thus, the ancestral and descendant states could be represented as *s*^*A*^=[*L, R*] and 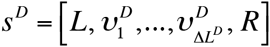, respectively. The ancestries satisfy 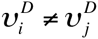 for *i* ≠ *j*, and their details depend on the responsible indel history. As long as 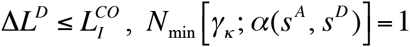, and there is only one fewest-indel history, 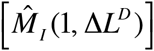, which consists of a single event that inserts the descendant sites in between the PASs. Therefore, the sum of contributions from the fewest-indel history is:

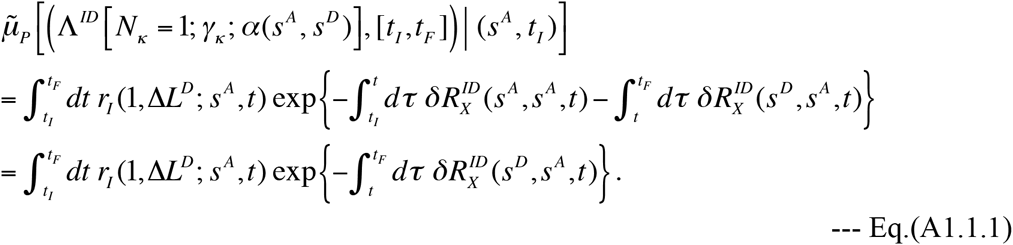

As in case (ii), each next-fewest indel history is composed of two indel events, and classified into two types: (c) two successive insertions, 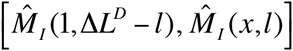 with *l* = 1,…, Δ*L*^*D*^ −1 and *x* = 1,…, Δ*L*^D^ − *l* +1; and (d) an insertion followed by a deletion, 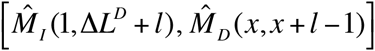 with 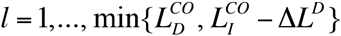 and *x* = 2,…, Δ*L*^*D*^ + 2. Thus, in this case, the portion contributed by the next-fewest indel histories is given by:

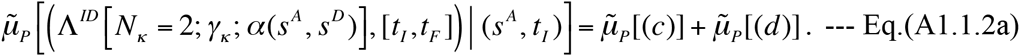

Here,

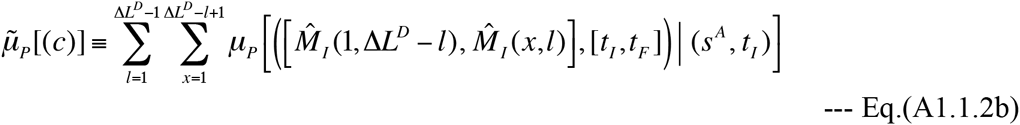

is the sum of contributions from the histories of type (c), and

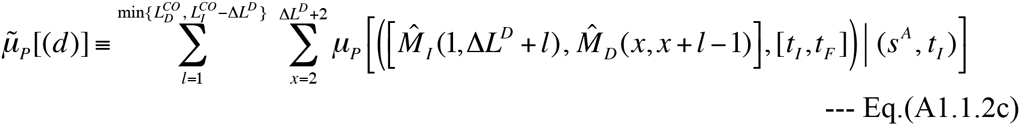

is the sum of contributions from the histories of type (d). As in case (i), let 〈[Δ*L*^*D*^−*l*]|≡〈*s*^*A*^|*M*^*I*^(1,Δ*L*^*D*^−*l*) be the intermediate state in each type (c) history. Then, the history’s contribution is calculated as:

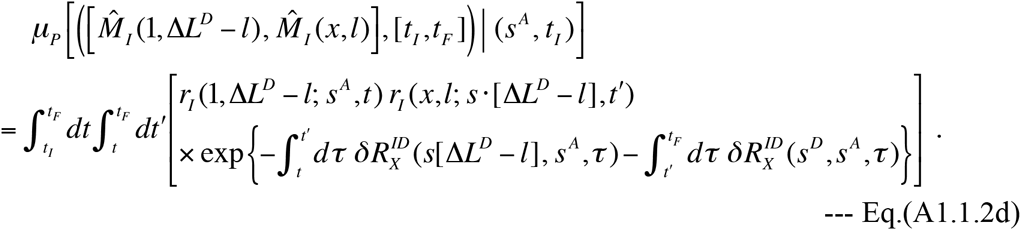

Similarly, each type (d) history’s contribution is calculated as:

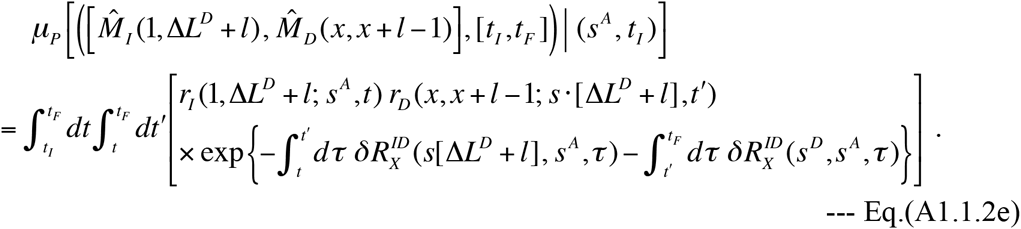

Let us now calculate Eqs.(A1.1.1,2a-e) under Dawg’s indel model or the “long indel” model, as in the previous cases in Section 1.2 of Results. In this model, we have 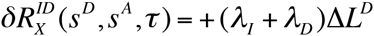 and Eq.(A1.1.1) becomes:

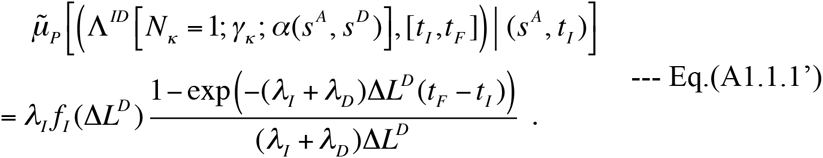

Similarly, using 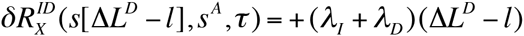 and 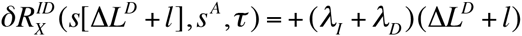 Eqs.(A1.1.2d,e) are calculated, respectively, as:

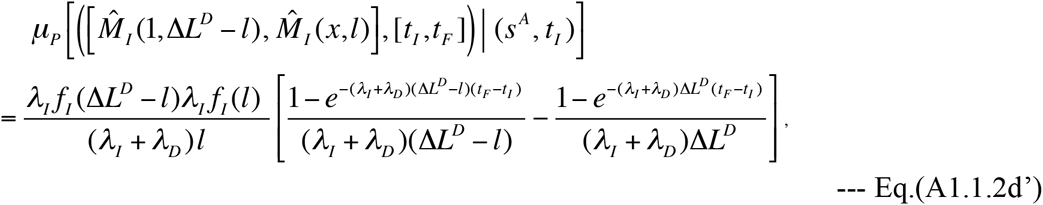

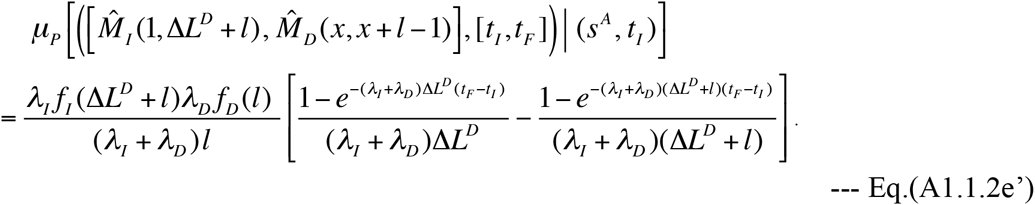

Substituting Eqs.(A1.1.2d’,e’) into Eqs.(A1.1.2a,b,c), we can concretely calculate the total contribution from next-fewest-indels.

#### A1.2. For case (iv) local PWAs

Here we detail the calculation of the sum of contributions from the fewest-indel events as well as that from the next-fewest-indel events in case (iv) considered in Section 1.2 of Results, that is, the case where both the ancestor and the descendant have one or more site(s) in between a pair of PASs, but no ancestral site is related to any of the descendant sites (Figure 2 D).

In case (iv), we assume that the ancestral and the descendant states have Δ*L*, ^*A*^ and Δ*L*^*D*^ sites, respectively, in between the flanking PASs. Thus, the ancestral and descendant states could be represented as *s*^*A*^ =;[*L*, 1,…, Δ*L*^*A*^, *R*]and 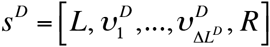 respectively. Here, the descendant ancestries satisfy 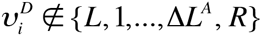 for all *i* = 1,…, Δ*L*^*D*^ and 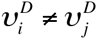 for all *i ≠ j*, and their details depend on the responsible indel history. As long as 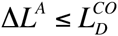 and 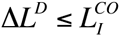, *N* _*min*_ [γ_*k*_; α(*s*^*A*^, *s*^*D*^)]=2As indicated by Eqs.(A1.3c’,d’) in Appendix A1 of part I (Ezawa, Graur and Landan 2015a), there are three types of fewest-indel histories: (e) the deletion of the ancestral sites followed by an insertion of Δ*L*^D^ sites, 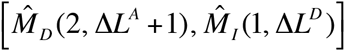 (f)an insertion immediately on the right of the ancestral sites to be deleted, followed by the deletion, 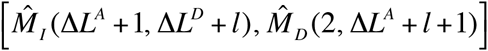 with 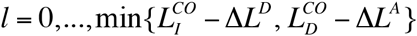; and (g) an insertion immediately on the left of the ancestral sites to be deleted, followed by the deletion, 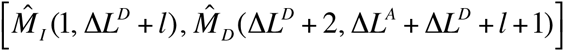 also with 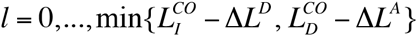 Thus, the sum of contributions by the fewest-indel histories is expressed as:

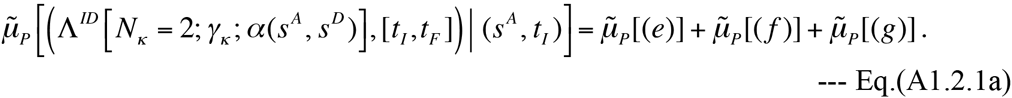

Here,

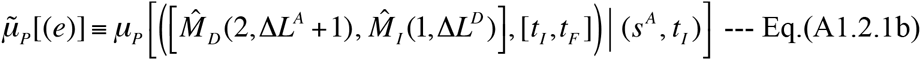

is the contribution from the single history of type (e),

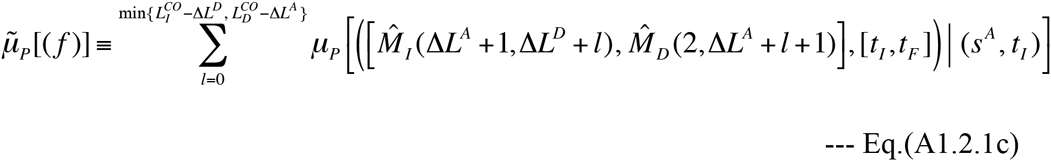

is the sum of contributions from the type (f) histories, and

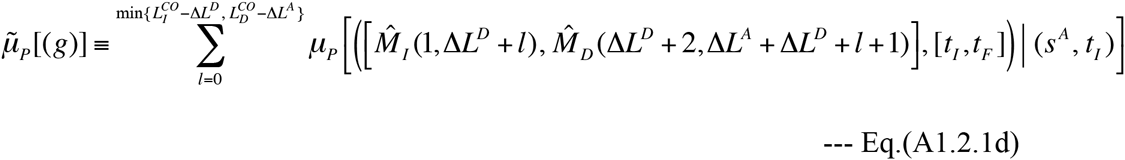

is the sum of contributions from the type (g) histories. To calculate the contribution from each of these histories, let 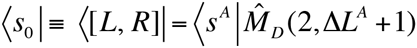 intermediate state of the type (e) history. And, as in case (ii), let 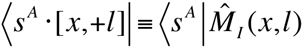 be the state type including the intermediate states of the histories of types (f) and (g). Then, the type (e) history’s contribution is:

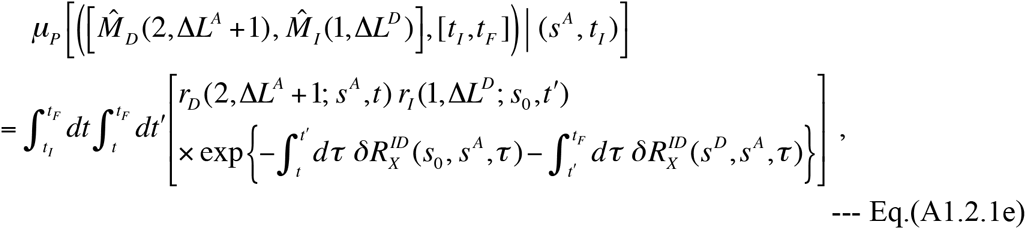

the contribution from each history of type (f) is:

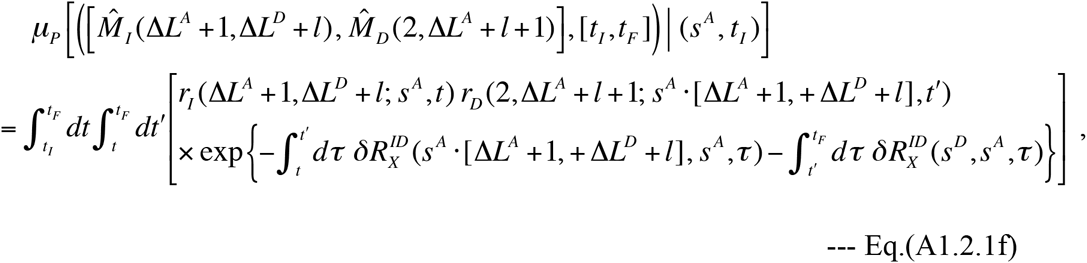

and each type (g) history’s contribution is:

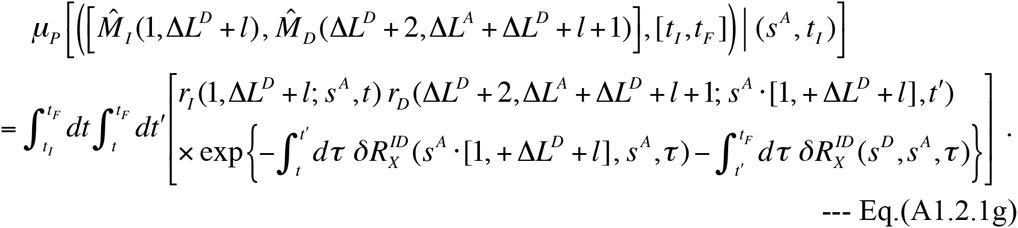

Under Dawg’s indel model (or, similarly, under the “long indel” model), we have 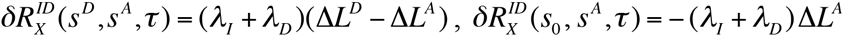 and 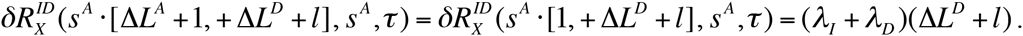 Using them, Eqs.(A1.2.1e,f,g) are calculated as:

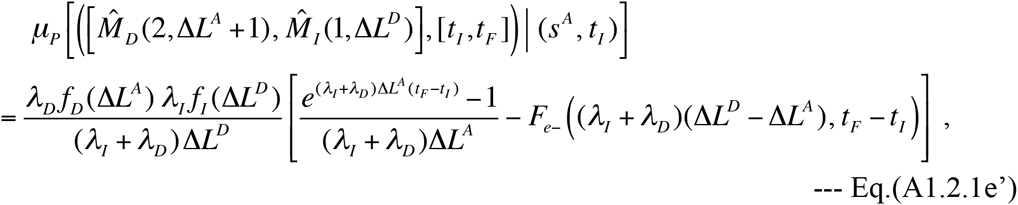

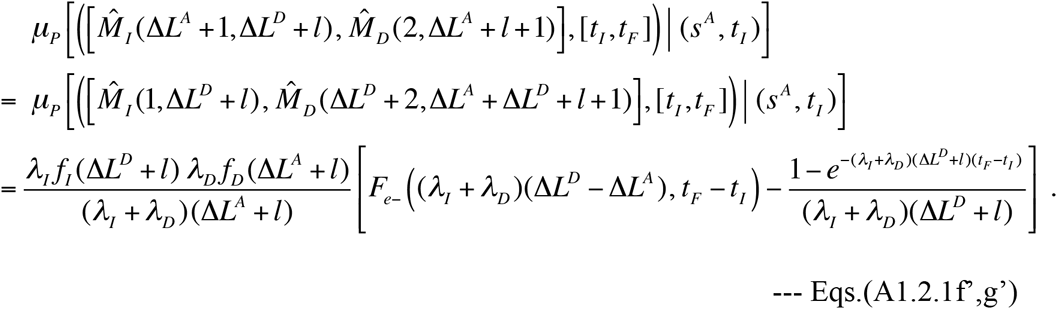

Here, we defined the function *F*_*e*_− (*x*, *t*) as:

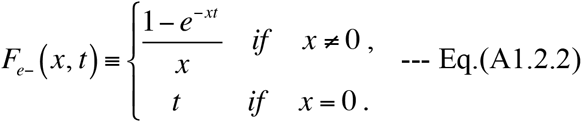

Meanwhile, each next-fewest-indel history is composed of three indel events, and classified into one of 6 broad types: (h) two successive deletions followed by an insertion; (i) a deletion, followed by an insertion, followed by a deletion; (j) an insertion followed by two successive deletions; (k) a deletion followed by two successive insertions; (l) an insertion, followed by a deletion, followed by an insertion; and (m) two successive insertions followed by a deletion. And these 6 broad types can be further sub-classified into 24 sub-types, as explained in the following.

Type (h) does not need to be sub-classified, and each history belonging to this type can be expressed as: 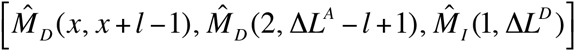 with *L* = 1,…, Δ*L*^*A*^ −1 and *x* = 2,…, Δ*L*^*A*^ − *L* + 2. Type (i) is sub-classified into three sub-types: (i-1) the case where the insertion is immediately on the right of the sites to be deleted, represented as 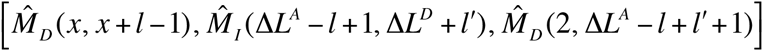 with *l*=1,…Δ*L*^*A*^−1, *x*=2,…,Δ*L*^*A*^−*l*+2, and 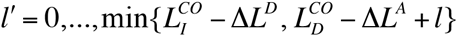 nd (i-2) the case where the insertion is immediately on the left of the sites to be deleted, represented as 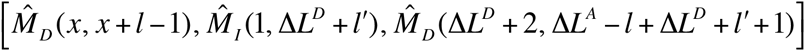 also with *l*=1,…Δ*L*^*A*^−1, *x*=2,…,Δ*L*^*A*^−*l*+2, and 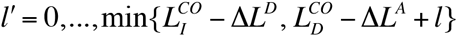 and (i-3) the case where the first event deletes all unpreserved ancestral sites, represented as 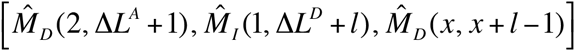 with 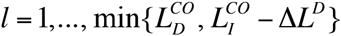 and *x* = 2,…, Δ*L*^*D*^ + 2. Type (j) is sub-classified into eight sub-types: (j-1) the case where the two deletions were on the right of the inserted sites that ended up in the descendant, represented as 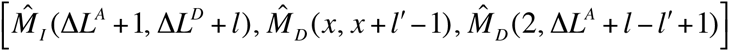 with 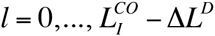, 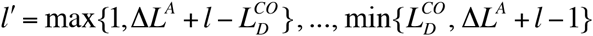, and *x*=2,…Δ*L*^*A*^+*l*−*l′*+*l*;(j-2) the case where the two deletions were on the left of the inserted sites that ended up in the descendant, represented as 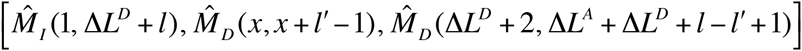 with 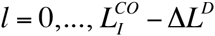, 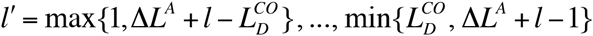 *x* = Δ*L* ^*D*^ + 2,…, Δ*L*^*A*^ + Δ*L*^*D*^ + *L* − *L*′+ 2; (j-3) the case where the first and the second deletions were on the left and in the middle, respectively, of the inserted sites that ended up in the descendant, represented as 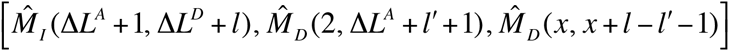 with 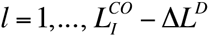 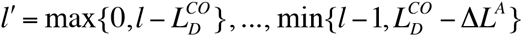, and *x*=3,…Δ*L*^*D*^+1Δ*L*^*D*^≥2 must always hold); (j-4) the case where the first and the second deletions were in the middle and on the left, respectively, of the inserted sites that ended up in the descendant, represented as 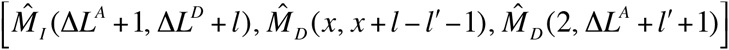 with 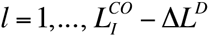 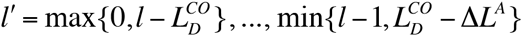 and *x*=Δ*L*^*A*^*l*′+3,…,Δ*L*^*A*^+Δ*L*^*D*^+*l*′+1 (Δ*L*^*D*^ ≥ 2 must always hold); (j-5) the case where the first and the second deletions were on the right and in the middle, respectively, of the inserted sites that ended up in the descendant, represented as 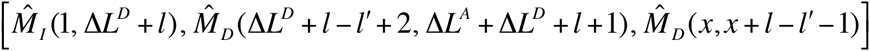 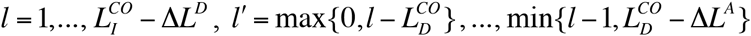 and *x*=3,…Δ*L*^*D*^+1 (Δ*L*^*D*^ ≥2 must always hold); (j-6) the case where the first and second deletions were in the middle and on the right, respectively, of the inserted sites that ended up in the descendant, represented as 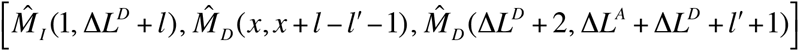 with 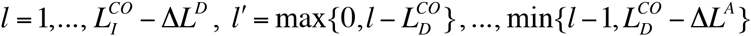, and *x*=3,…Δ*L*^*D*^+1 (Δ*L*^*D*^ ≥2 must always hold); (j-7) the case the first and the second deletions were on the right and on the left, respectively, of the inserted sites that ended up in the descendant, represented as 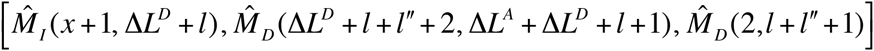 with 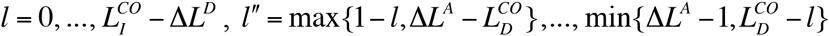, and *x* = max{0, *L*″},…, min{*L* + *L*″, Δ*L*^*A*^}; and (j-8) the case where the first and the second deletions were on the left and on the right, respectively, of the inserted sites that ended up in the descendant, represented as 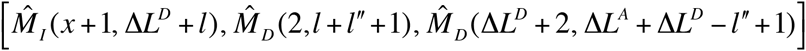 with 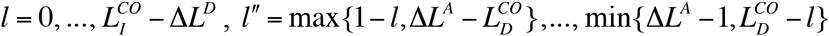 and *x*= max{0,*l*″},…,min{*l*+*l*″Δ*L*^*A*^}Type (k) does not need be sub-classified, and each history belonging to this type can be represented as: 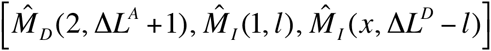 with *l*=1,…,*l*+1. Type (l) is sub-classified into three sub-types: (l-1) the case where the first insertion was on the immediate right of the left-flanking PAS, and at least a site inserted by the event survived, represented as 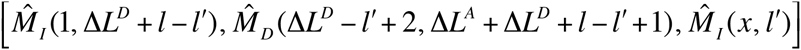 with 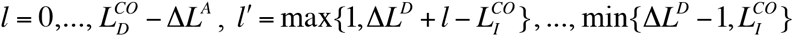, and *x*=1,…,Δ*L*^*A*^−*l*′+1;(l-2) the case where the first insertion was on the immediate left of the right-flanking PAS, and at least a site inserted by the event survived, 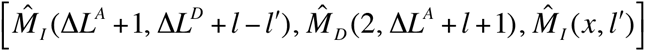 with 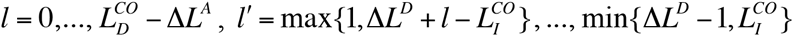, and *x*=1,…,Δ*L*^*A*^−*l*′+1; and and (l-3) the case where all sites inserted by the first event were deleted along with the unpreserved ancestral sites, represented as 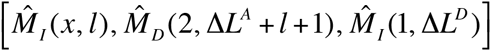 with 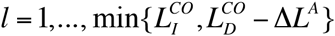 and *x*=1,…, Δ*L*^*A*^+1. Finally, type (m) is sub-classified into eight sub-types: (m-1) the case where the two insertions are on the left of the unpreserved ancestral sites, represented as: 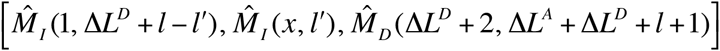 with 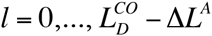, 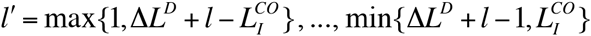 and *x* = 1,…, Δ*LD* + *L* − *L*′+1; (m-2) the case where the two insertions are on the right of the unpreserved ancestral sites, represented as 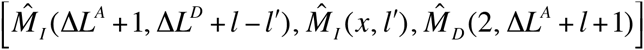 with 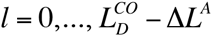, 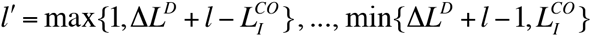. and *x* = Δ*L*^*A*^ +1,…, Δ*L*^*A*^ + Δ*L^D^* + *L* − *L*′+1; (m-3) the case where the first and the second insertions are in the middle and on the left, respectively, of the unpreserved ancestral sites, represented as 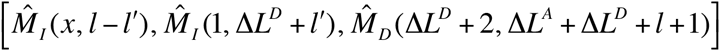 with 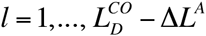, and 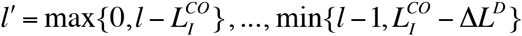 (Δ*L*^*A*^ ≥ 2 must always hold); (m-4) the case where the first and the second insertions are in the middle and on the left and in the middle, respectively, of the unpreserved ancestral sites, represented as 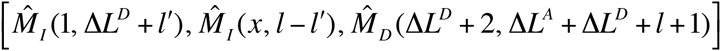 with 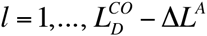, 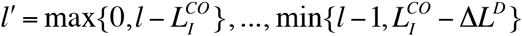, and *x* = Δ*L*^*D*^ + *L*′+ 2,…, Δ*L*^*A*^ + Δ*L*^*D*^ + *L*′ (Δ*L*^*A*^ ≥ 2 must always hold); (m-5) the case where first and the second insertions are in the middle and on the right, respectively, of the unpreserved ancestral sites, represented as 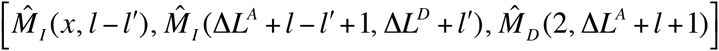 with *x*=2,…, Δ*L*^*A*^ 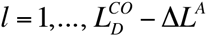, and 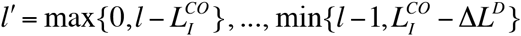 (Δ*L*^*A*^ ≥ 2 must always hold); (m-6) the case where the first and the second insertions are on the right and in the middle, respectively, of the unpreserved ancestral sites, represented as 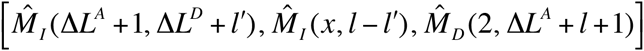 with *x* = 2,…, Δ*L*^*A*^, 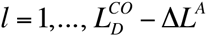, and 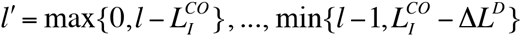 (Δ*L*^*A*^ ≥ 2 must always hold); (m-7) the case where the first and the second insertions are on the left and on the right, respectively, of the unpreserved ancestral sites, represented as 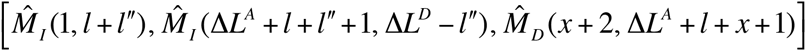 with 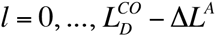 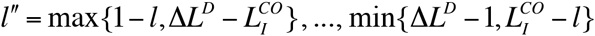, and *x* = max{0, *l*″},…, min{*l* + *l*″, Δ*L*^D^}; and (m-8) the case where the first and the second insertions are on the right and on the left, respectively, of the unpreserved ancestral sites, represented as 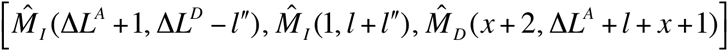 with 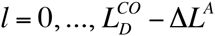 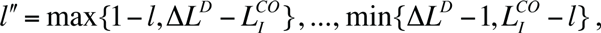, and *x* = max{0, *l*″},…, min{*l* + *l*″, Δ*L*^D^}.

Using this classification of the next-fewest-indel histories in case (iv), the sum of their contributions can be expressed as:

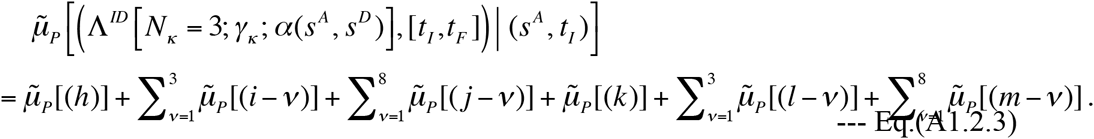

Here, each of 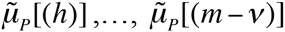 is the sum of contributions from the histories belonging to one of the 24 sub-types explained above. Each of them can be calculated by calculating the contribution from each constituent history according to the definition, Eq.(4.1.1b) of part I supplemented by Eq.(3.1.8b) of part I, and by summing the contributions over possible values of the length variables *l* and *l*′ (or *l*″) and over possible positions *x* when necessary, all specified above. Under a space-and time-homogeneous model, the indel rates become independent of the positions of the indels and of the exact state before the event, and the exit rate depends only on the sequence length (and possibly on the functional form of the deletion length distribution). This simplifies the calculation considerably. Especially, because each term is independent of the exact position of indels, a summation over the positions *x* is reduced to a simple multiplication by the number of possible positions. In addition, considerations on spatial symmetry reveal that some different sub-types actually give identical contributions to each other (detailed below). Moreover, under Dawg’s indel model or the “long indel” model, the increment of the exit rate is proportional to the difference in the sequence lengths, which further simplifies the calculation. Here, we give the results of contributions by individual histories under Dawg’s indel model.

But the results apply immediately to, *e.g*., the “long indel” model as well. We first give the general formula for the contribution of a three-event local history, and then we apply the formula to the histories of different sub-types.

First we consider a general three-event history, 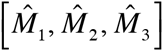, on the ancestral state S^A^=[*L*, 1,…Δ*L*^A^, *R*] that resulted in the descendant state 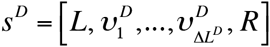 Here 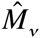 (with ν = 1, 2, 3) is either an insertion 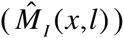 or a deletion 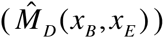. Let *δl_v_* (with ν = 1, 2, 3) be the sequence length change caused by the event 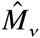; *δl_v_* is positive if 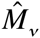 is an insertion and negative if 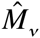 is a deletion. They satisfy: *δl*1 + *δl*2 + *δl*3 = Δ*L*_D_ − Δ*L*_A_. And let *r*(*δl*) be the space-and time-homogeneous rate of the indel event that changes the sequence length by *δl*. Under Dawg’s indel model, we have:

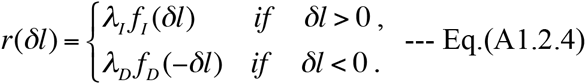

Then, according to Eq.(4.1.1b) of part I supplemented by Eq.(3.1.8b) of part I, the contribution from this three-event local history is expressed as:

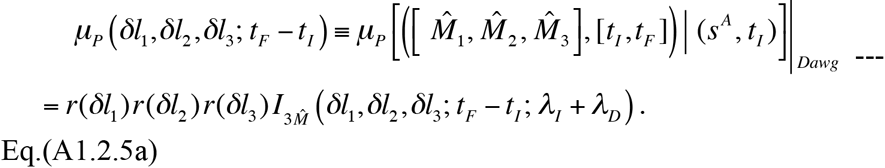

Here, 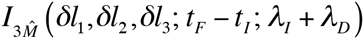 is a triple-time integral:

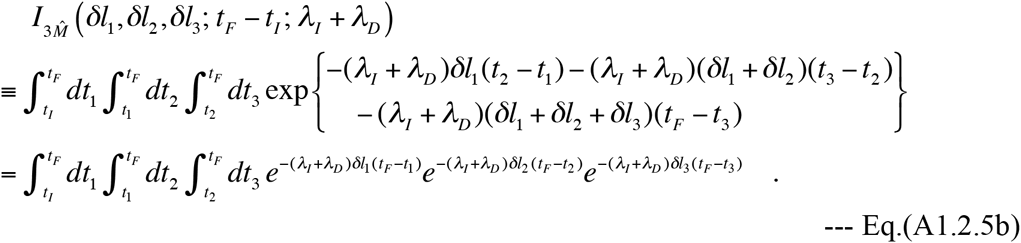

When *δl*2 + *δl*3 ≠ 0, it is calculated as:

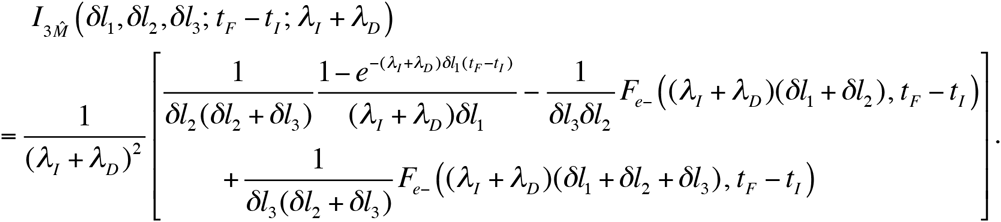

Here the function *F*_*e*_− (*x*, *t*) was already defined in Eq.(SR-8B.2). When *δl*2 + *δl*3 = 0, the triple-time integral is calculated as:

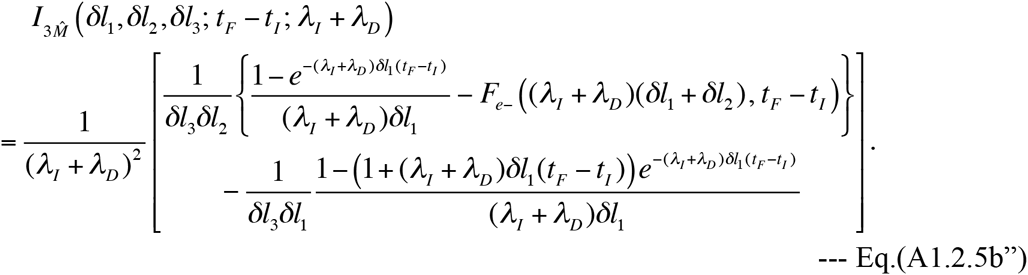

When 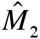 does not overlap 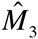, there should be a corresponding pair of events, 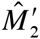 and 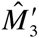 that satisfy 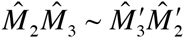 as one of the binary equivalence relations (Eqs.(2.3.3a-d) of part I). Under Dawg’s model, the effects of 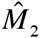 and 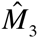 are *partially* factorable if we consider the joint contribution from the two local histories, 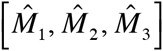 and 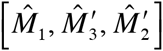. We will express this joint contribution as:

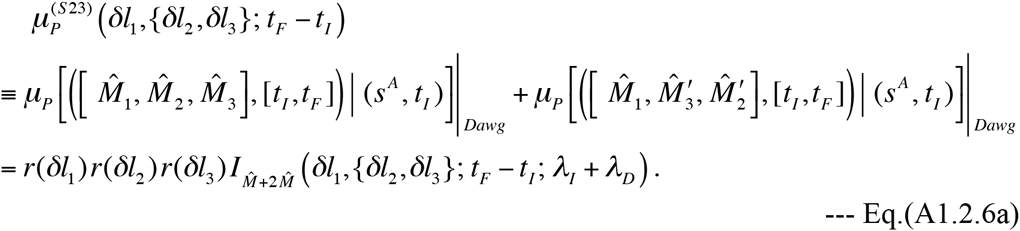

Here, 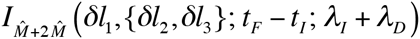 is a triple-time integral:

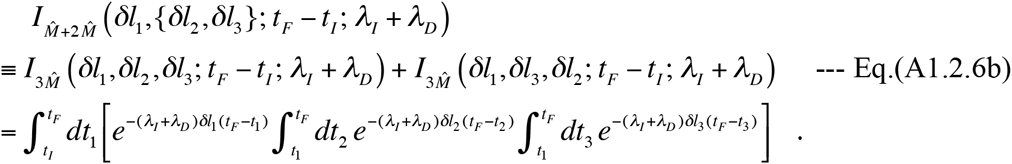

It is calculated as:

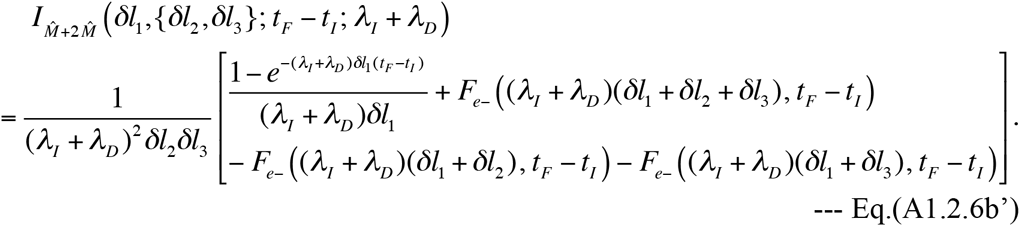

Similarly, when 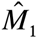 does not overlap 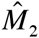, there should be a corresponding pair of events, 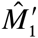 and 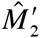, that satisfy the binary equivalence, 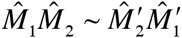 (as one of Eqs.(2.3.3a-d) of part I). When considering the joint contribution from the two local histories, 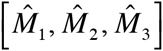 and 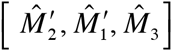, the effects of 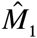 and 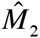 are *partially* factorable under Dawg’s model. We will express the joint effect as:

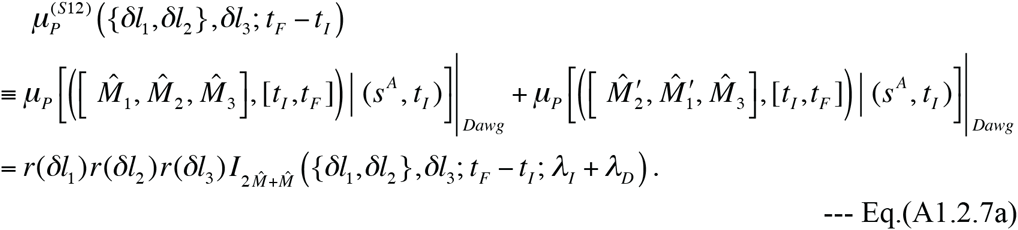

Here, 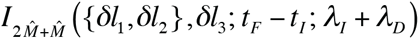 is a triple-time integral:

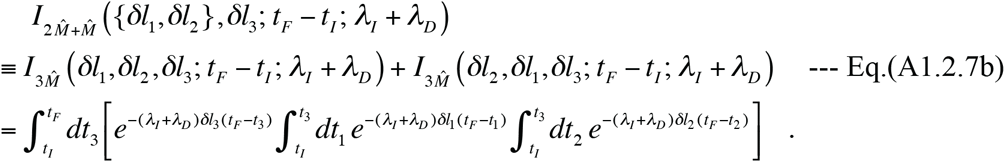

It could be calculated by taking advantage of the following relationship:

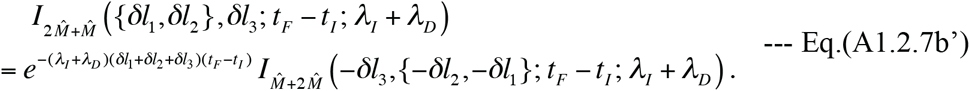

Now we can apply the general formulas, Eqs(A1.2.5a,b”), Eqs.(A1.2.6a,b’), and Eqs.(A1.2.7a,b’), to specific cases. First, the contribution from a type (h) history is:

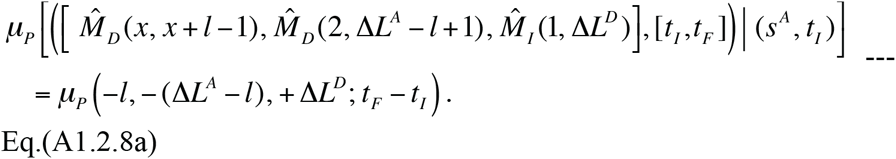

Second, by spatial symmetry, the contribution from a type (i-1) history is identical to that from the corresponding type (i-2) history. They are given by:

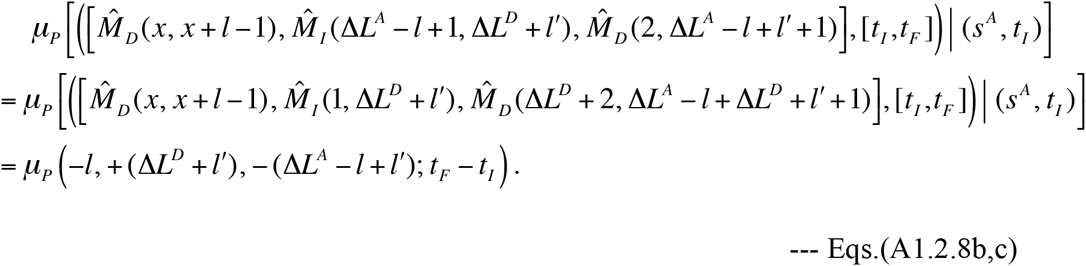

Third, the contribution from a type (i-3) history is:

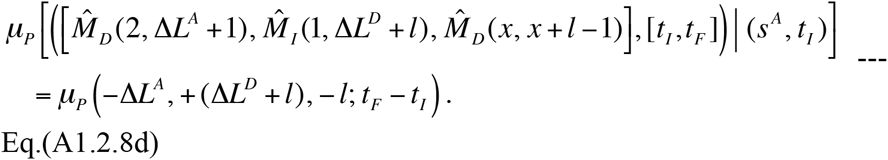

Fourth, by spatial symmetry, the contribution from a type (j-1) history is identical to that from the corresponding type (j-2) history. They are given by:

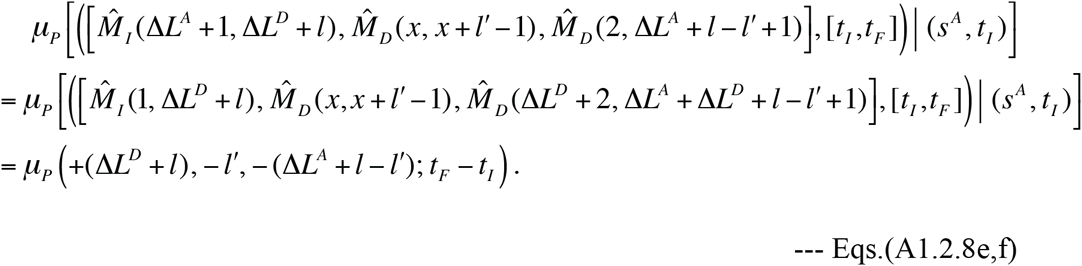

Fifth, we consider the joint contribution from a type (j-3) history and the corresponding type (j-4) history. By spatial symmetry, it is identical to the joint contribution from the corresponding type (j-5) and type (j-6) histories. They are given by:

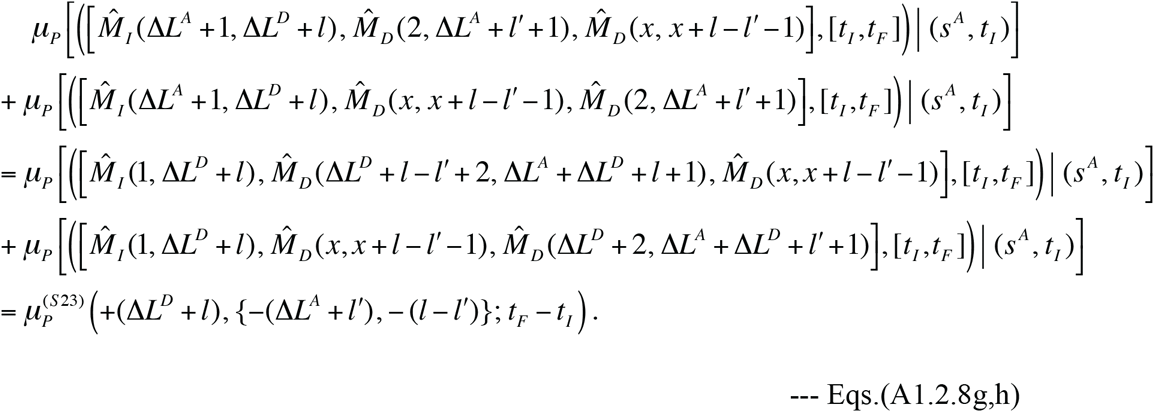

Sixth, consider the joint contribution from a type (j-7) history and the corresponding type (j-8) history. It is given by:

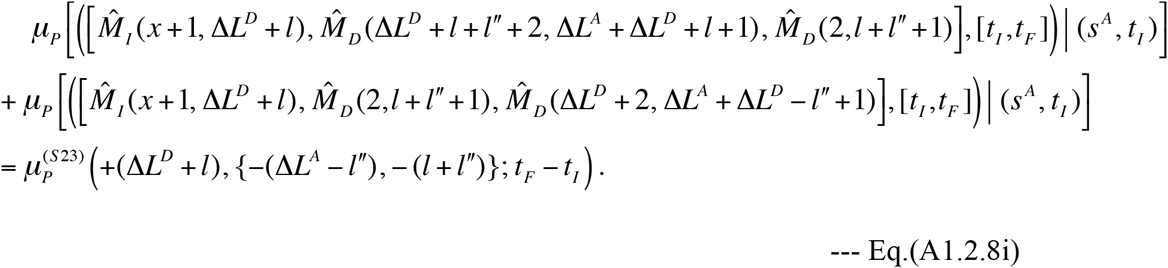

Seventh, the contribution from a type (k) history is:

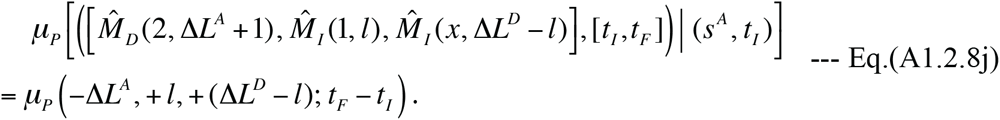

Eighth, by spatial symmetry, the contribution from a type (l-1) history is identical to that from the corresponding type (l-2) history. They are given by:

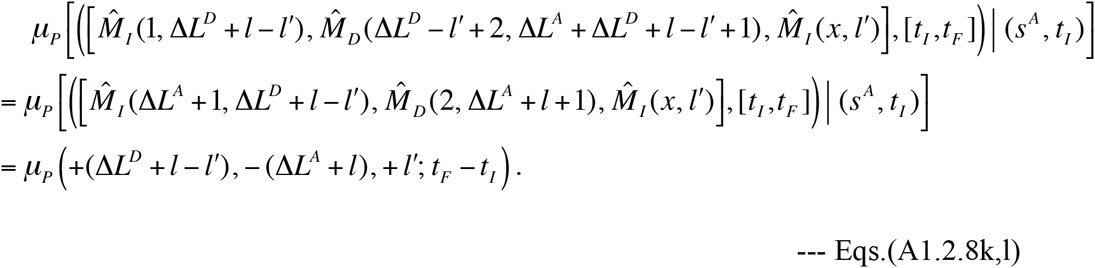

Ninth, the contribution from a type (l-3) history is:

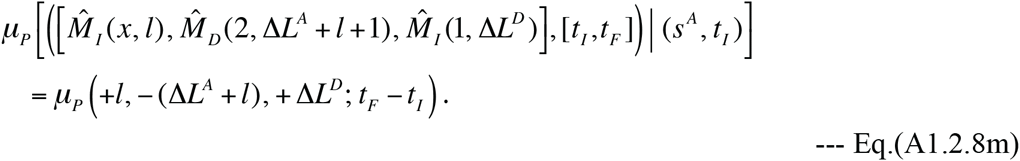

Tenth, by spatial symmetry, the contribution from a type (m-1) history is identical to that from the corresponding type (m-2) history. They are given by:

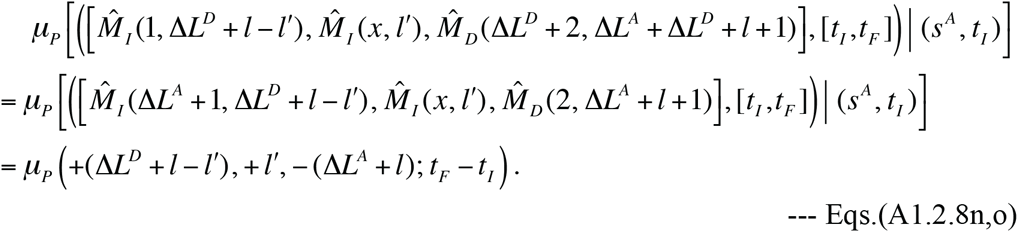

Eleventh, consider the joint contribution from a type (m-3) history and the corresponding type (m-4) history. By spatial symmetry, it is equal to the joint contribution from the corresponding type (m-5) and type (m-6) histories. They are given by:

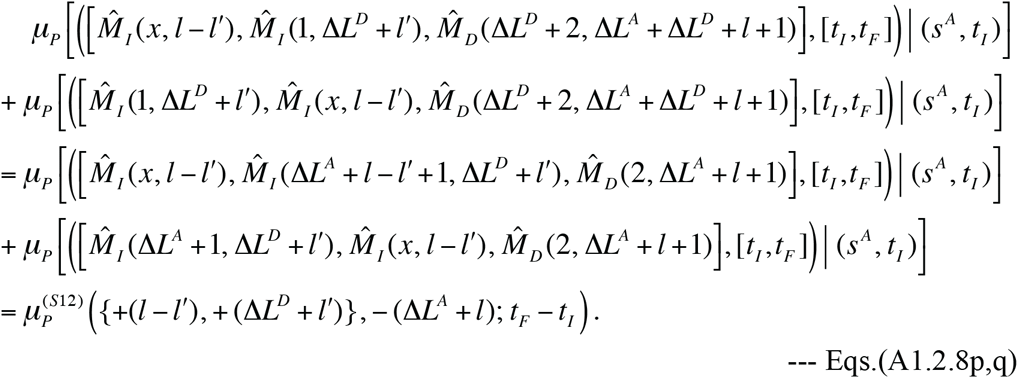

And, finally, twelfth, consider the joint contribution from a type (m-7) history and the corresponding type (m-8) history. It is given by:

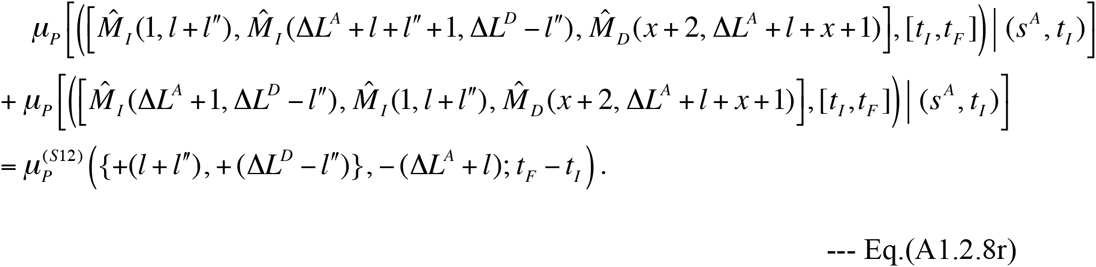

These equations, Eqs.(A1.2.8a-r), supplemented with the general formulas, Eqs.(A1.2.5,6,7), provide the elementary contributions from the individual next-fewest-indel local histories. By summing them up over the appropriate lengths and positions given above the general formulas, Eqs.(A1.2.5,6,7), we get the total contribution of the next-fewest-indel local histories.

#### A1.3. System of integral equations providing “exact” multiplication factors for case-(i) & (iii) local PWAs

In Section 1.2 of Results, we derived Eq.(1.2.7), which, supplemented with Eq.(1.2.8), gives a system of integral equations that can in principle provide “exact” multiplication factors for cases (i) and (ii), where there are a non-negative integer (Δ*L*^*A*^) of ancestral sites but zero descendant sites in between a pair of PASs. Here, following a similar line of procedures, we will derive a system of integral equations that can in principle provide “exact” multiplication factors *for cases (i) and (iii)*, where there are zero ancestral sites but a non-negative integer (Δ*L*^D^) of descendant sites in between a pair of PASs. For this purpose, we assume a setting very similar to that made above Eq.(1.2.6) for cases (i) and (ii). The only notable difference is that the ancestral and the descendant states here are *s*^*A*^ = [*L*, *R*] and 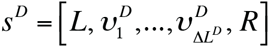 respectively, which will be denoted as *s*^*A*^ = 〈0| and *s*^*D*^| = 〈Δ*L*^*D*^| under the setting of (local) spatial homogeneity.

The starting point here is the fundamental integral equation, Eq.(3.1.2) of part I (Ezawa, Graur and Landan 2015a), for the stochastic evolution operator 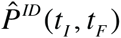, instead of Eq.(3.1.4) of part I for cases (i) and (ii). Similarly to above Eq.(1.2.6), we sandwich Eq.(3.1.2) of part I with 〈*s*^*A*^| and |*s*^*D*^ 〉, expand 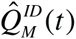 using its definition (*i.e*., Eq.(3.1.1c) of part I supplemented with Eqs.(2.4.2b’,c’) of part I), and ignore the effects of indels that are irrelevant to the cases under consideration. Then, we get:

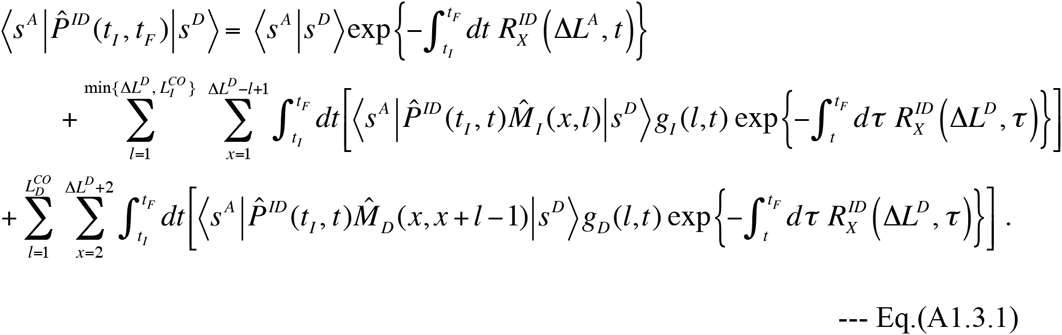

Here, when considering the domains of the summations, we used the fact that the ketvectors 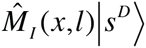 and 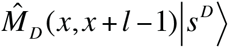 correspond uniquely to the states 〈Δ*l*^*D*^ − *l*| and 〈Δ*L*^*D*^ + *l*|, respectively. Next, we take advantage of this fact again, take account of the local homogeneity, and use the notation, 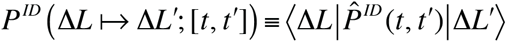, introduced above Eq.(1.2.7). Then, we can rewrite Eq.(A1.3.1) as:

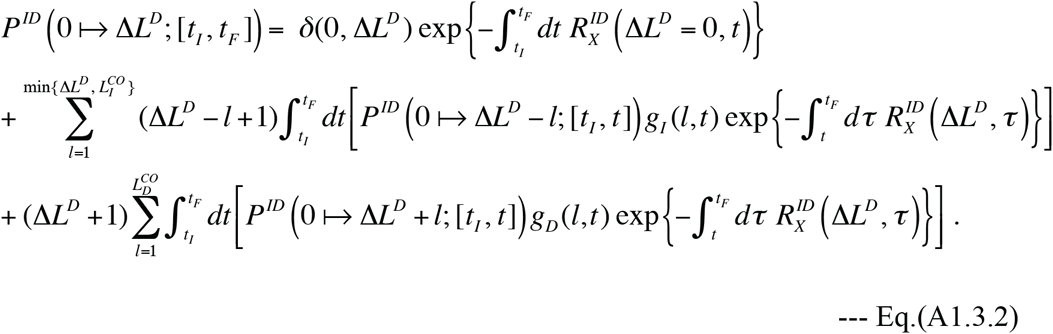

This gives the system of integral equations for the “exact” probabilities, *P*^*ID*^ (0 ↦Δ*L*^D^; [*t*, *t*]), with non-negative integers Δ*L*^D^ = 0,1, 2,…. This system of equations can be solved iteratively, again starting with the “zero-event approximation,” 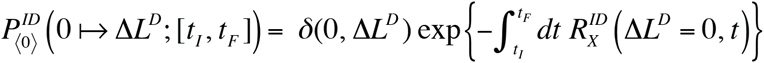 After a desired number (say, *N*_*ID*_) of iteration steps, the multiplication factor will be obtained similarly, and we have:

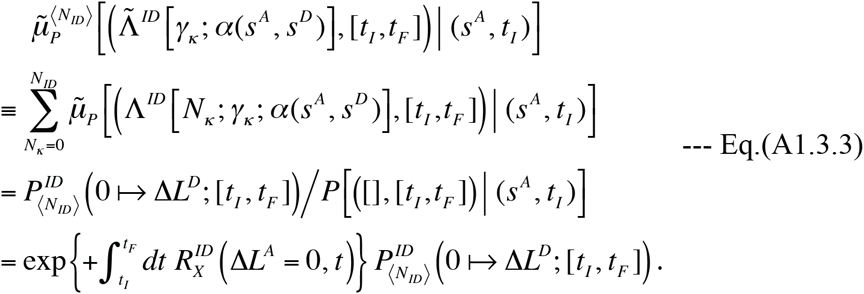

It should be noted that the exponent on the right-most hand side is the time integral of the exit rate of the ancestral state but not that of the descendant state. This is just due to the definition of the multiplication factor (Eq.(4.1.1b) of part I). The consideration of the time-and space-complexities of a naïve implementation of the iteration algorithm goes just as below Eq.(1.2.8) of Results. Just as Eq.(1.2.7), Eq.(A1.3.2) could also be numerically solved via a system of “two-sub-step” recursion relations:

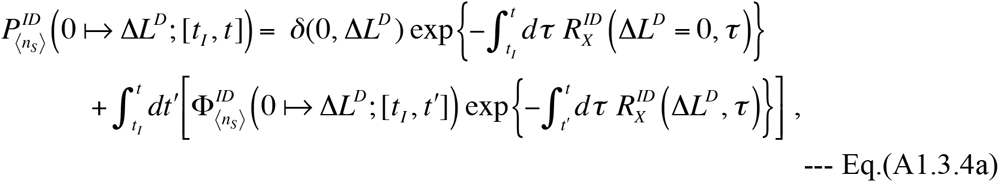

and

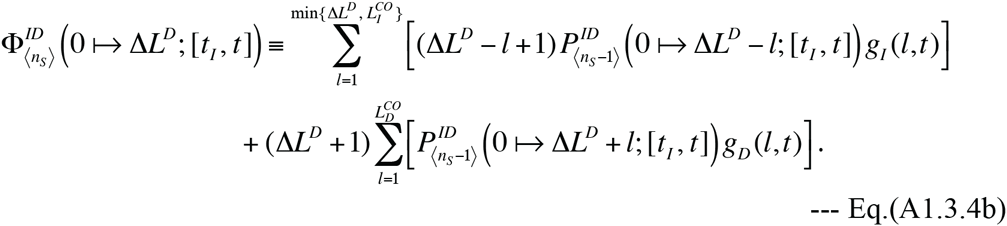

Similarly to Eqs.(1.2.9a,b), the algorithm to numerically solve this system of recursion relations also has the time-complexity of *O* (*N*^*ID*^ *L*^*CO*^ (*L*^*CO*^ + *N*_*P*_)*N*_*P*_) and the space-complexity of *O* (*L*^*CO*^ *N*_*P*_), and a single run of the algorithm (with *N*_*ID*_ steps) outputs 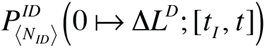 for all 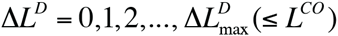 and at all time-points, 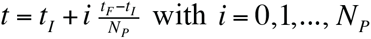

#### A2. Perturbation calculation of multiplication factors for MSAs: case (IV) in Section 1.3

Here we detail the calculation of the portion of the multiplication factor contributed by the fewest-indel events as well as that contributed by the next-fewest-indel events in case (IV). This case was considered in Section 1.3 of Results, and its situation is illustrated in Figure 4 E. In this case, the external sequence states were represented as: *S*_1_=[*L*,1,…,Δ*L*^*D*1^], *S*_2_=[*L*, 1,…,,*i*,*j*,…,Δ*L*^*D*1^, R], and S_3_=[*L*, *R*]with 1≤*i*+1<*j*≤Δ*L*^*D*1^+1 but (*i*, *j*)≠(0, Δ*L*^*D*1^+1)Also in this case,

*N*_min_[*C*_*k*_;α[*S*_1_, *S*_2_, *S*_3_]; *T*] and there were two fewest-indel local histories. One, which we call type (E) here, starts with the root state 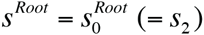 It is represented by Eq.(1.3.14a), *i.e*.,

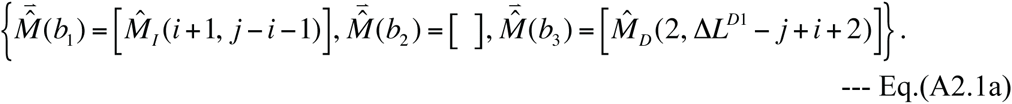

And the other, which we call type (F) here, starts with the root state

*s*^Root^ = *s*_1_ = [*L*, 1,…, Δ*L*^*D*1^, *R*], instead of 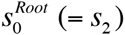. It is represented by Eq.(1.3.14b), *i.e.*,

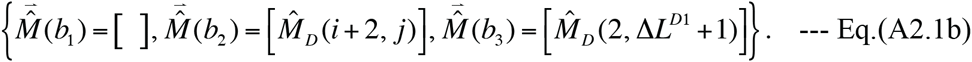

Thus the portion of the multiplication factor due to the fewest-indel histories can be written as:

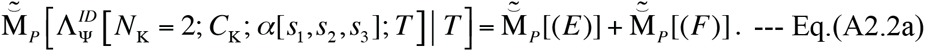

Here 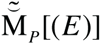 and 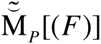 are the contributions from the local histories of type (E) and type (F), respectively. These contributions can be calculated according to the definition, Eq.(1.1.2b) supplemented by Eqs.(4.2.4b,6b,8) of part I (Ezawa, Graur and Landan 2015a), similarly to the calculation of 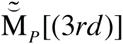 in case (III). First, the contribution from the type (E) history is:

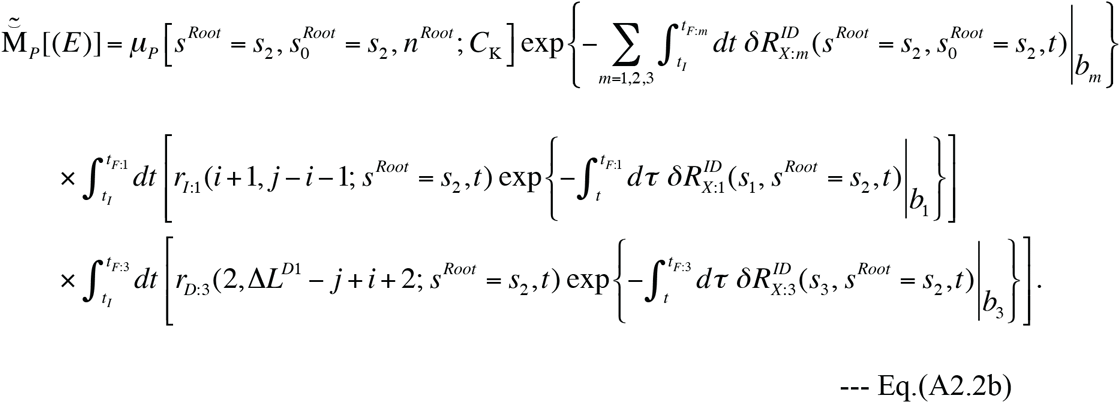

And the contribution from the type (F) history is:

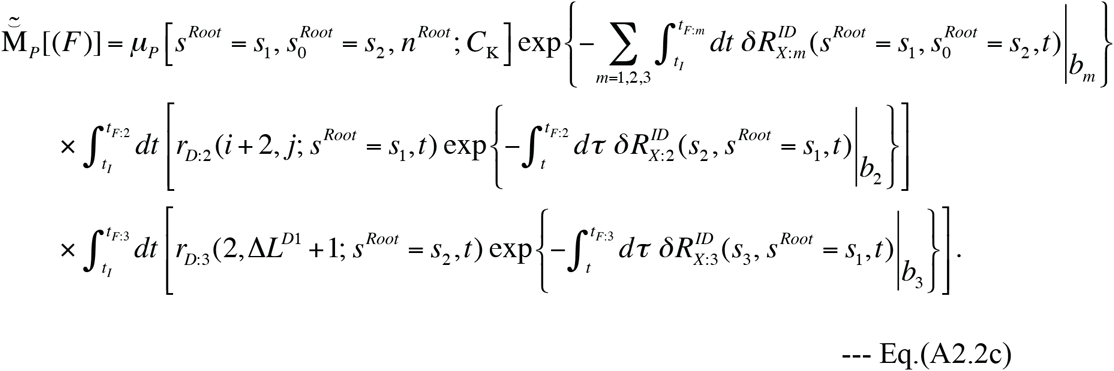

Assuming a uniform distribution of the ancestral sequence length, we have 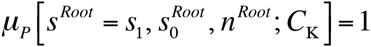 no matter what the root state (*s*^*Root*^) is. Then, under Dawg’s model, Eqs.(SR-9.2b,c) can be calculated as:

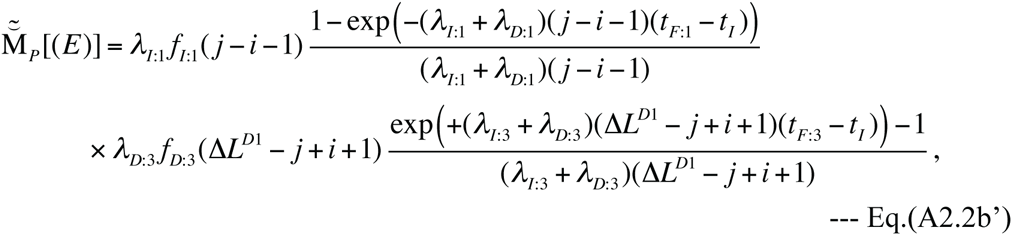

and

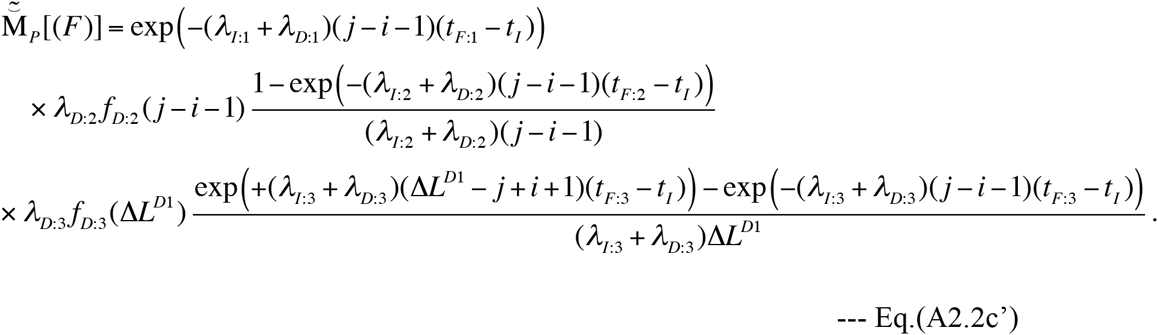

The next-fewest-indel local histories consist of three indels each, and can be broadly classified into four types according to the root sequence state: (G) the histories with 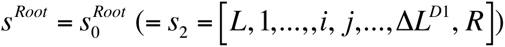 (H) those with *s*^Root^ = *s*_1_ = [*L*, 1,…, Δ*L*^*D*1^, *R*]; (I) the histories having the root states made from *s*_1_ by removing a single connected fraction out of the run of sites missing in *s*_2_, *i.e*., *s*^*Root*^ = [*L*, 1,…, *i*′, *j*′,…., Δ*L*^*D*1^, *R*](with *i*′ ≥ *i*, *i*′+2 ≤ *j*′ ≤ *j*, but (*i*′, *j*′) ≠ (*i*, *j*)); and (J) the histories having the root states made by adding some extra sites to *s*_1_, *i.e*., 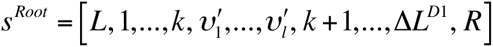 with *k* = *i*,…, *j* −1, 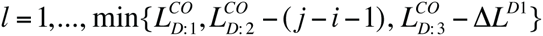 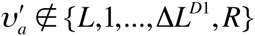 for ^∀^a ∈{1,…, *l*}, and 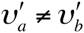 for ^∀^*a*≠*b*(∈ {1,…, *l*}). These four broad types are further sub-classified into 10 sub-types in total.

First, type (G) local histories are sub-classified into four sub-types: (G-1) the case where two successive insertions occur along *b*_1_, represented as 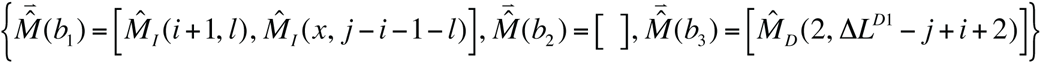 with *l* = 1,…, *j*−*i*−2 and *x* = *i*+1,…, *i*+*l*+1(*j*−*i* ≥ 3 must hold); (G-2) the case where a “long” insertion and a subsequent deletion occur along *b*_1_, represented as

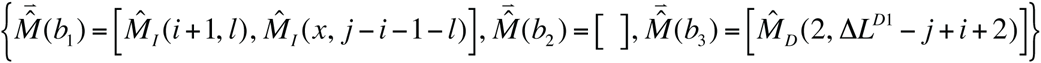

with 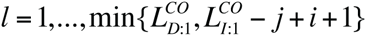 and *x* = *i*+2,…, *j*+1; (G-3) the case where two successive deletions occur along *b*_3_, represented as

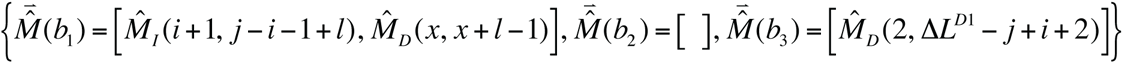

with *l* = 1,…, Δ*L*^*D*1^ − *j* + *i* and *x* = 2,…, Δ*L*^*D*1^ − *j* + *i* − l + 3 (Δ*L*^*D*1^ − *j* + *i* ≥ 1 must hold)and (G-4) the case where an insertion and a subsequent “long” deletion occur along *b*_3_, represented as

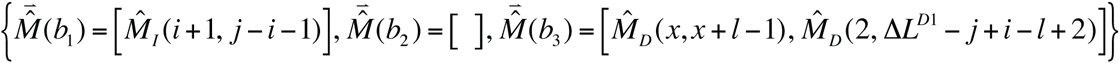

with 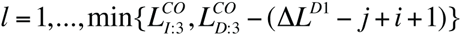 and *x* = 1,…, Δ*L*^*D*1^ − *j*+*i*+2.

Second, type (H) local histories are also sub-classified into four sub-types: (H-1) the case where two successive deletions occur along *b*_2_, represented as

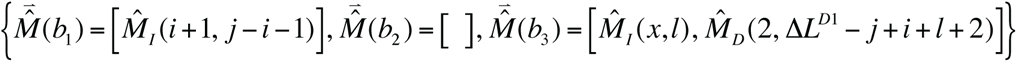

with *l* = 1,…, *j*−*i*−2 and *x* = *i*+2,…, *j*−*l*+1 (*j*−*i*≥3 must hold); (H-2) the case where an insertion and a subsequent “long” deletion occur along *b*_2_, represented as

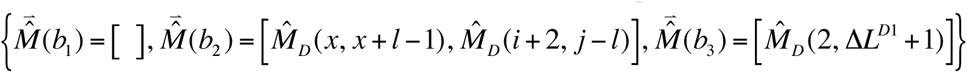

with 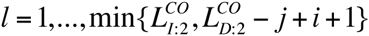 and *x* = *i*+1,…, *j*; (H-3) the case where two successive deletions occur along *b*_3_, represented as

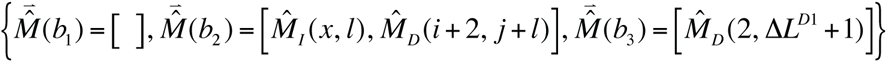

with *l* = 1,…, Δ*L*^*D*1^−1 and *x* = 2,…, Δ*L*^*D*1^ − *l*+2; and (H-4) the case where an insertion and a subsequent “long” deletion occur along *b*_3_, represented as

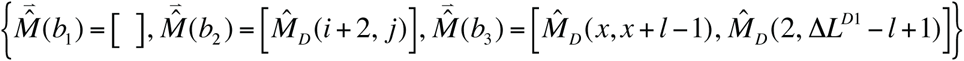

with 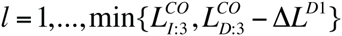 and *x*=1,…, Δ*L*^*D*1^+1.

Third, type (I) does not need be sub-classified, and each history of this type can be represented as:

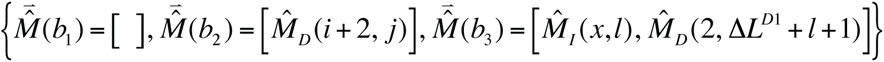

which is determined uniquely from the root state (*s*^*Root*^ = [*L*, 1,…, *i*′, *j*′,…, Δ*L*^*D*1^, *R*] with *i*′≥*i*, *i*′+2 ≤ *j*′ ≤ *j*, but also with (*i*′, *j*′) ≠ (*i*, *j*)). Fourth, type (J) does not need be sub-classified, either, and each history of this type can be represented as:

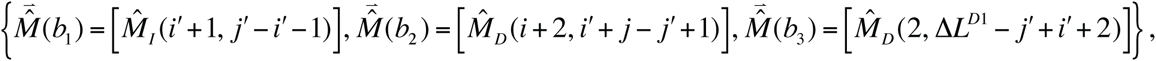

which, again, is determined uniquely from the root state 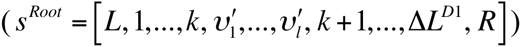.

Thus, the summed contributions from the next-fewest-indel local histories in case (IV) can be expressed as:

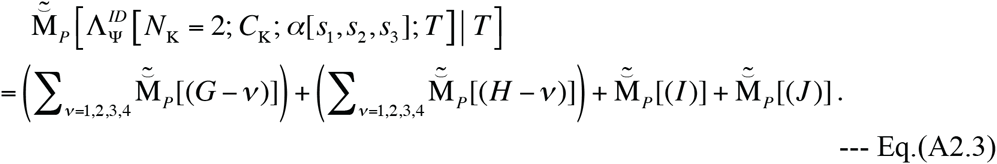

Here 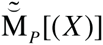 (with *X* = *G*−1,…, *J*) represents the summed contribution from the histories over one of the aforementioned 10 sub-types. Each such term can be calculated by summing the contributions from individual local histories belonging to each sub-type over the possible lengths and possible positions, if necessary. Under a space-homogeneous model, the summation over possible positions will be reduced to the multiplication by the number of possible positions (for each fixed set of lengths). And the contributions from individual local histories can be calculated according to the definition, Eq.(1.1.2b) supplemented by Eqs.(4.2.4b,6b,8) of part I, and taking advantage of, or by slightly modifying, Eqs.(1.2.4,5d,5e) and Eqs.(A1.1.1,2d,2e) for contributions from single-event and two-event local histories for PWAs. In the following, we will give the expressions for the contributions from individual local histories of respective sub-types under Dawg’s indel model.

Type (G-1):

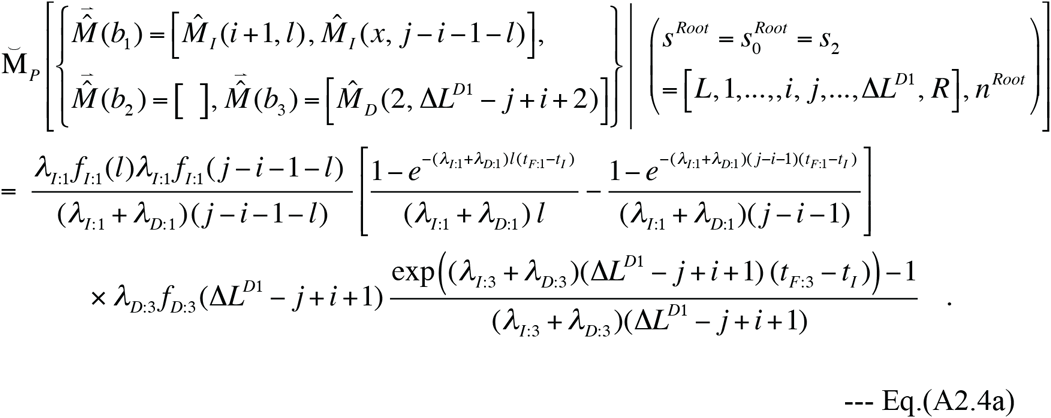

Type (G-2):

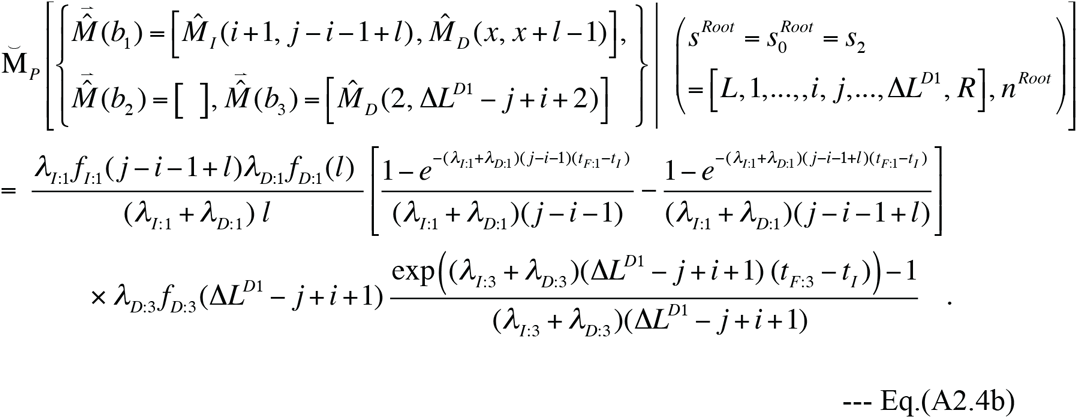

Type (G-3):

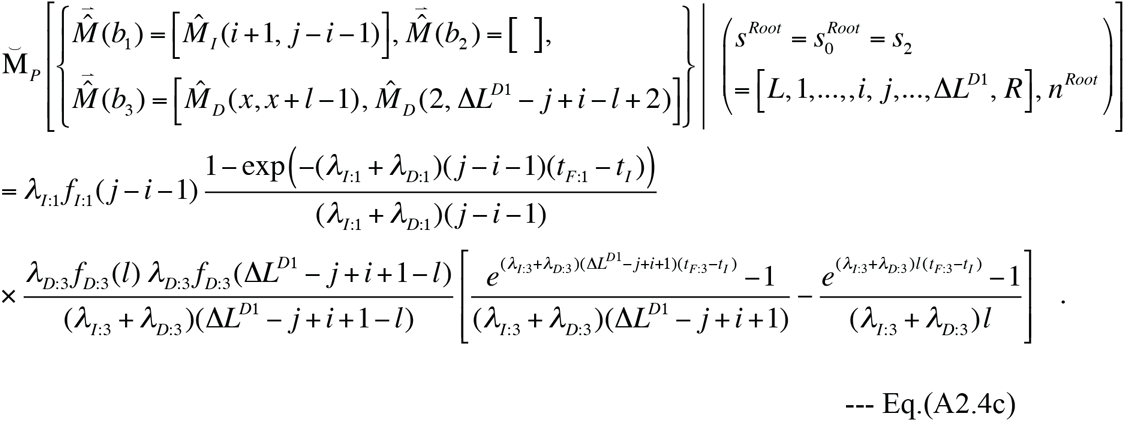

Type (G-4):

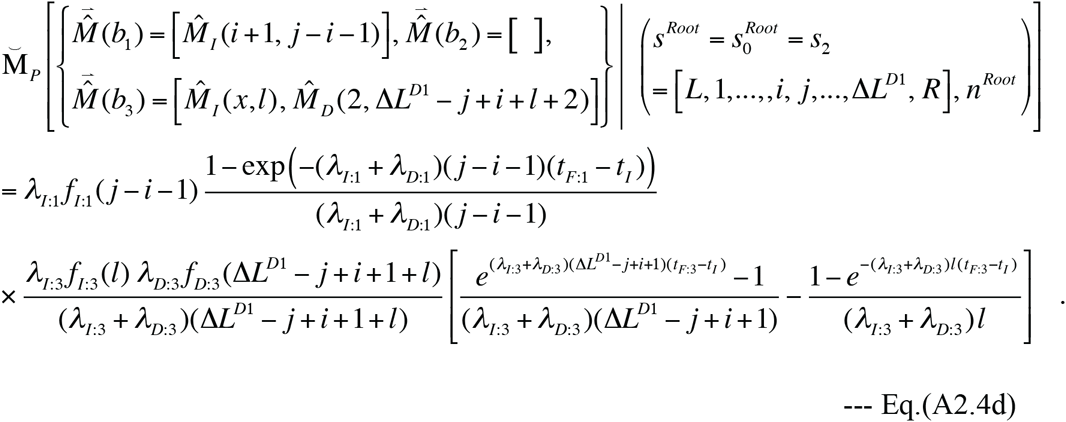

Type (H-1):

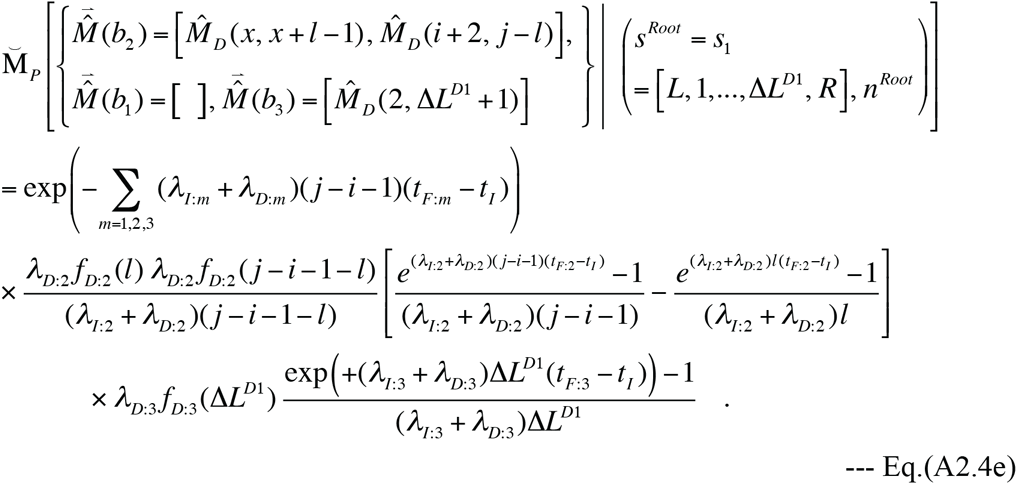

Type (H-2):

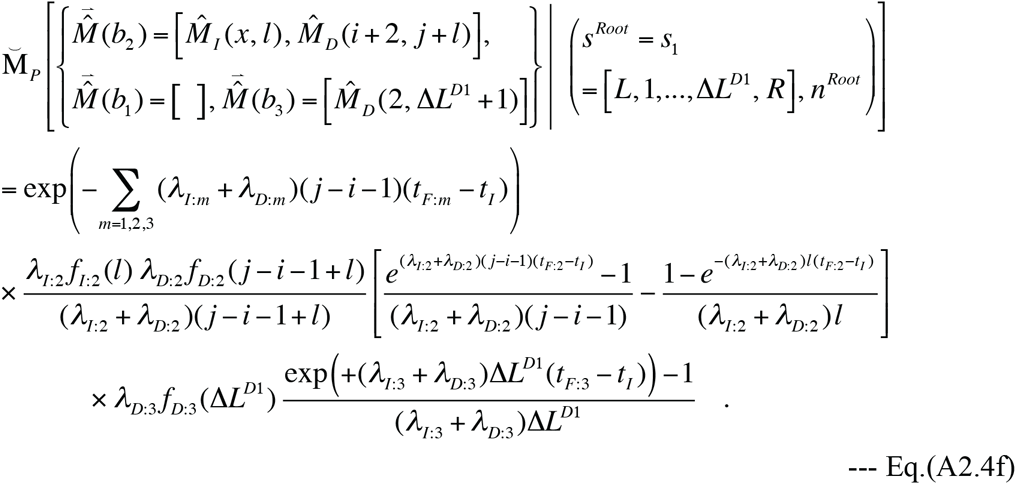

Type (H-3):

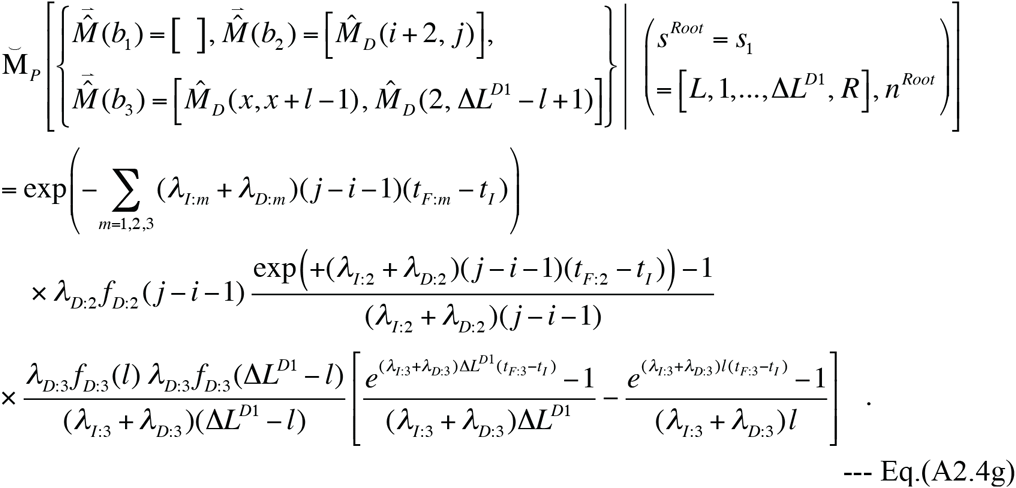

Type (H-4):

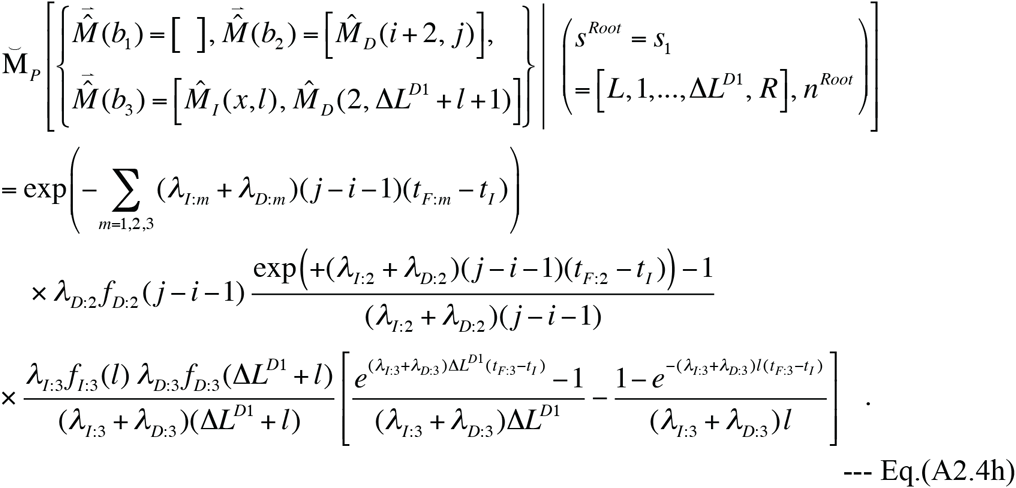

Type (I):

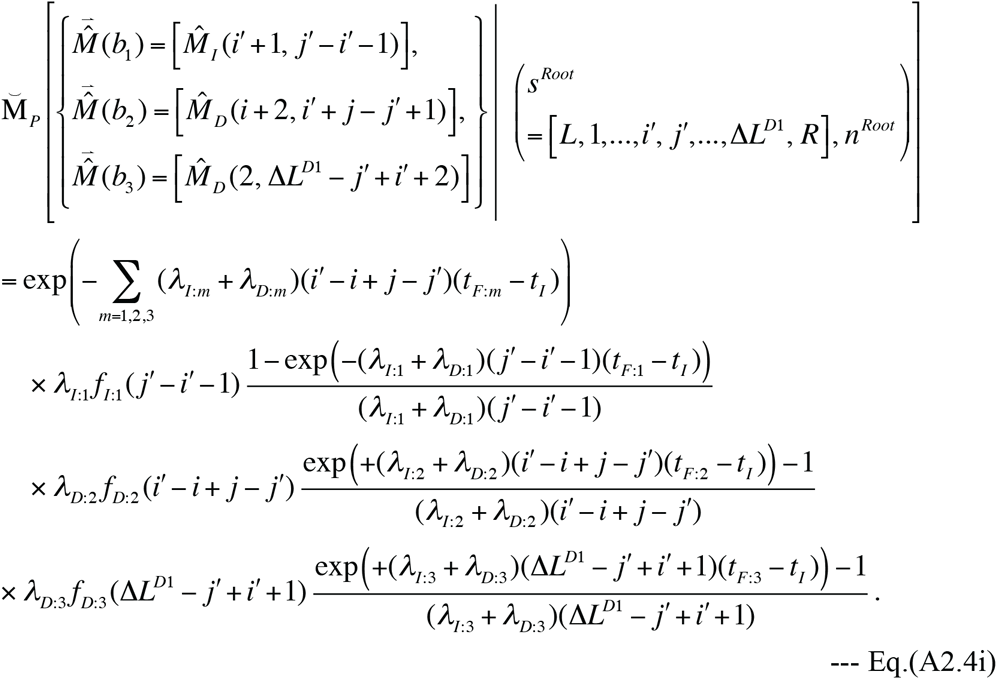

It should be noted that the right hand side of this equation depends on *i*′ and *j*′ only through *l*′ ≡ − *j*′−*i*′−1. Thus, it follows that there are *j*−*i*−*l*′ histories with the same *l*′ (and therefore the same probability) but with different *i*′’s (namely, *i*′ = *i*,…, *j*−1−*l*′).

Type (J):

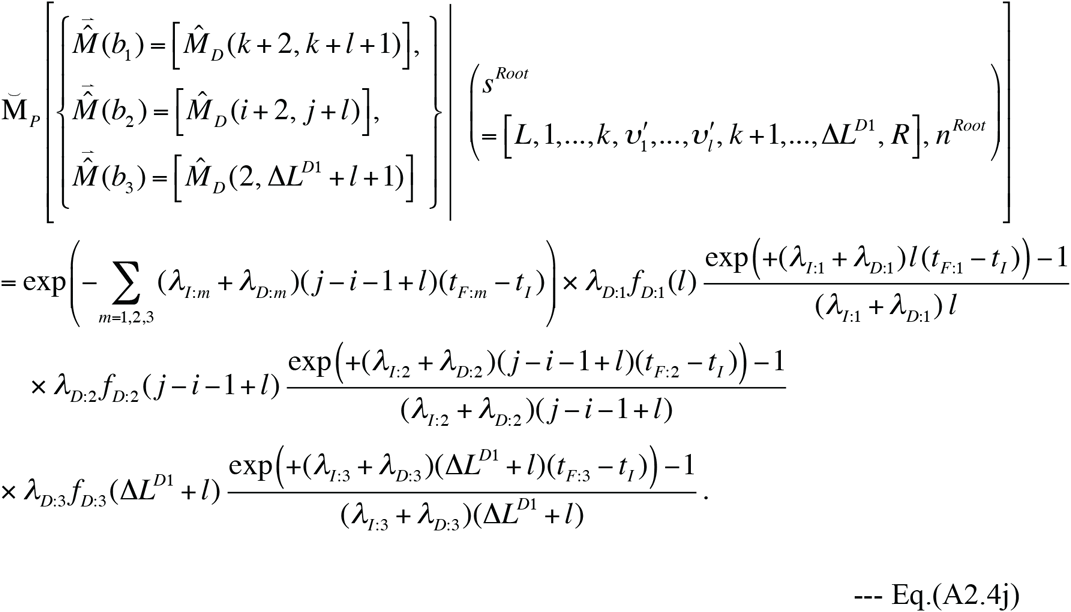

It should be noted that the right hand side of this equation does not depend on *k* whereas it does depend on *l*. Thus, it follows that there are *j*−*i* histories with the same *l* (and therefore the same probability) but with different *k*’s (namely, *k* = *i*,…, *j*−1).

Summing each of Eqs.(A2.4a-j) over the lengths and positions specified above Eq.(A2.3), we obtain 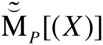 with *X* = *G*−1,…, *J*, the summation of which (according to Eq.(A2.3)) gives the total contribution from the next-fewest-indel local histories.

